# Allele-specific DNA methylation is increased in cancers and its dense mapping in normal plus neoplastic cells increases the yield of disease-associated regulatory SNPs

**DOI:** 10.1101/815605

**Authors:** Catherine Do, Emmanuel LP Dumont, Martha Salas, Angelica Castano, Huthayfa Mujahed, Leonel Maldonado, Arunjot Singh, Sonia C. DaSilva-Arnold, Govind Bhagat, Soren Lehman, Angela M. Christiano, Subha Madhavan, Peter L. Nagy, Peter H.R. Green, Rena Feinman, Cornelia Trimble, Nicholas P. Illsley, Karen Marder, Lawrence Honig, Catherine Monk, Andre Goy, Kar Chow, Samuel Goldlust, George Kaptain, David Siegel, Benjamin Tycko

## Abstract

**Background:** Mapping of allele-specific DNA methylation (ASM) can be a post-GWAS strategy for localizing regulatory sequence polymorphisms (rSNPs). However, the advantages of this approach, and the mechanisms underlying ASM in normal and neoplastic cells, remain to be clarified.

**Results:** We performed whole genome methyl-seq on diverse normal cells and tissues and three types of cancers (multiple myeloma, lymphoma, glioblastoma multiforme). After excluding imprinting, the data pinpointed 15,114 high-confidence ASM differentially methylated regions (DMRs), of which 1,842 contained SNPs in strong linkage disequilibrium or coinciding with GWAS peaks. ASM frequencies were increased 5 to 9-fold in cancers vs. matched normal tissues, due to widespread allele-specific hypomethylation and focal allele-specific hypermethylation in poised chromatin. Cancers showed increased allele switching at ASM loci, but disruptive SNPs in specific classes of CTCF and transcription factor (TF) binding motifs were similarly correlated with ASM in cancer and non-cancer. Rare somatic mutations affecting these same motif classes tracked with de novo ASM in the cancers. Allele-specific TF binding from ChIP-seq was enriched among ASM loci, but most ASM DMRs lacked such annotations, and some were found in otherwise uninformative “chromatin deserts”.

**Conclusions:** ASM is increased in cancers but occurs by a shared mechanism involving disruptive SNPs in CTCF and TF binding sites in both normal and neoplastic cells. Dense ASM mapping in normal plus cancer samples reveals candidate rSNPs that are difficult to find by other approaches. Together with GWAS data, these rSNPs can nominate specific transcriptional pathways in susceptibility to autoimmune, neuropsychiatric, and neoplastic diseases. Custom genome browser tracks with annotated ASM loci can be viewed at a UCSC browser session hosted by our laboratory (https://bit.ly/tycko-asm)

## Background

Genome-wide association studies (GWAS) have implicated numerous DNA sequence variants, mostly single nucleotide polymorphisms (SNPs) in non-coding regions, as candidates for mediating inter-individual differences in disease susceptibility. However, to promote GWAS statistical signals to biological true-positives, and to identify the functional sequence variants that underlie these signals, several obstacles need to be overcome. Multiple statistical comparisons demand stringent thresholds for significance, p<5×10^-8^ for a GWAS, and this level can lead to the rejection of biological true-positives with sub-threshold p-values [1]. A more fundamental challenge is identifying the causal regulatory SNPs (rSNPs) among the typically large number of variants that are in linkage disequilibrium (LD) with a GWAS peak SNP. Combined genetic-epigenetic mapping can address these challenges. In particular, identification of non-imprinted allele-specific CpG methylation dictated by cis-acting effects of local genotypes or haplotypes (sometimes abbreviated as hap-ASM but hereafter referred to simply as ASM), led us and others to suggest that mapping this type of allelic asymmetry could prove useful as a “post-GWAS” method for localizing rSNPs [2–12]. The premise is that the presence of an ASM DMR can indicate a bona fide regulatory sequence variant (or regulatory haplotype) in that genomic region, which declares itself by conferring the physical asymmetry (i.e. ASM) between the two alleles in heterozygotes. ASM mapping, and related but technically more demanding post-GWAS approaches such as allele-specific chromatin immunoprecipitation-sequencing (ChIP-seq) [13, 14] can facilitate genome-wide screening for disease-linked rSNPs, which can then be prioritized for functional studies. However, the unique advantages of ASM mapping, and its potential non-redundancy with other post-GWAS mapping methods, remain to be clarified.

Genome-wide analysis of ASM by methylation sequencing (methyl-seq) is also yielding insights to the fundamental mechanisms that shape DNA methylation patterns. Our previous data using bisulfite sequence capture (Agilent SureSelect) revealed ASM DMRs and methylation quantitative trait loci (mQTLs) in human brain cells and tissues, and in T lymphocytes, and uncovered a role for polymorphic CTCF and transcription factor (TF) binding sites in producing ASM [8]. Others have pursued similar approaches with progressively greater genomic coverage [10, 11], with substantial though partial overlap in the resulting lists of ASM DMRs [9], and with consistent conclusions regarding the importance of disruptive SNPs in CTCF and TF binding sites as a mechanism underlying ASM. However, since ASM can be somewhat tissue-specific and its mapping requires heterozygotes at one or more “index SNPs” in the DMR, constraints from the numbers of individuals and numbers of cell types have limited the harvest of high-confidence ASM DMRs. These factors have in turn limited the assessment of specific classes of TF and CTCF binding sites for their involvement in ASM and limited the yield of candidate rSNPs in disease-associated chromosomal regions. Further, while some studies of cancer samples have been done using targeted methyl-seq [15–18], the genome-wide features and mechanisms of ASM in human neoplasia have yet to be clarified.

To address these issues, we have expanded our previous methyl-seq dataset and carried out whole genome bisulfite sequencing (WGBS) on a new large series of human samples spanning a range of tissues and cell types from multiple individuals, plus three types of human cancers. We identify high-confidence ASM DMRs using stringent criteria, perform extensive validations, apply a multi-step analytical pipeline to compare mechanisms of ASM in normal and cancer cells, and assess the unique strengths of dense ASM mapping for finding mechanistically informative disease associated rSNPs.

## Results

### Mapping of high-confidence ASM regions in normal and neoplastic human samples

The biological samples in this study are listed in **Table S1**, and our experimental approaches for identifying ASM DMRs, testing ASM mechanisms, and nominating disease-associated rSNPs are diagrammed in **Figure S1A, B**. The sample set included diverse tissues and purified cell types from multiple individuals, with an emphasis on immune system cells, brain, carcinoma precursor lineages and several other normal tissues and cell types, plus a set of primary cancers including multiple myeloma, B cell lymphoma, and glioblastoma multiforme (GBM) (**Table S1** and **Figure S2**). Since placental trophoblast has epigenetic and biological similarities to cancers [19–23], we also analyzed unfractionated placental tissue (chorionic plate) and purified placental trophoblast. Agilent SureSelect methyl-seq is a sequence capture-based method for genome-wide bisulfite sequencing that covers 3.7 million CpGs, located in all RefSeq genes and concentrated in promoter regions, CpG islands, CpG island shores, shelves, and DNAse I hypersensitive sites. We previously applied this method to 13 human samples [8] and for the current study we added samples so that the final SureSelect series includes a total of 24 samples of normal tissues and purified cell types, plus one lymphoblastoid cell line (LCL; GM12878). All samples were from different individuals, except for a trio among the brain samples consisting of one frontal cortex (Brodmann area BA9) and two temporal cortex samples (BA37 and BA38) from the same autopsy brain (**Table S1**).

To further increase the number of samples and cell types, to obtain complete genomic coverage, and to include cancer samples, we performed WGBS on 81 human samples. As listed in **Table S1** and **Fig. S2**, the non-cancer tissues and cell types included a set of immune system cells (T cells, B cells, monocyte/macrophages, whole blood and whole reactive lymph node) from multiple individuals, whole and fractionated samples including purified villous cytotrophoblast and extravillous trophoblast from a term placenta, several normal liver samples, primary bladder and mammary epithelial cells from multiple individuals, and whole and fractionated (NeuN-positive neurons and NeuN-negative glia) samples from cerebral cortex of multiple autopsy cases, plus the GM12878 LCL. The WGBS series included 16 primary human cancers, comprising 3 B cell lymphomas, 7 multiple myeloma cases (CD138+ cells from bone marrow aspirates), and 6 cases of glioblastoma multiforme (GBM). While the two series were mostly distinct, 5 of the non-cancer samples were analyzed by both SureSelect and WGBS (**Table S1**).

Numbers of mapped reads and depth of sequencing are in **Table S1**, and numbers of informative (heterozygous) SNPs are in **Figure S2**. As a quality control, we performed Principle Component Analysis (PCA) using net methylation values for CpGs informative in both SureSelect and WGBS. This procedure revealed the expected segregation of samples according to cell and tissue type and cancer or non-cancer status. It also revealed some expected findings for cell lineages, particularly highlighting both similarities and differences in methylation patterns in the brain cells (whole cerebral cortex, glia, neurons) and the GBMs (**Fig. S3A**). As another aspect of the quality control, when the same biological samples were analyzed on SureSelect and WGBS, the corresponding data points clustered closely together by PCA (e.g. the GM12878 LCL; **Fig. S3A**).

Our analytical pipeline (**Figure S1A, B**) includes steps to identify and rank ASM DMRs for strength and confidence and utilize the resulting maps, together with public ENCODE and related data, for testing mechanistic hypotheses. For ASM calling, we separated the SureSelect and WGBS reads by alleles using SNPs that were not destroyed by the bisulfite conversion, and defined ASM DMRs by at least 3 CpGs with significant allelic asymmetry in fractional methylation (Fisher’s exact test p<0.05). We further required at least 2 contiguous CpGs with ASM, an absolute difference in fractional methylation of ≥20% between alleles after averaging over all covered CpGs in the DMR, and an overall difference in fractional methylation between alleles passing a Benjamini-Hochberg (B-H) corrected Wilcoxon p-value (false discovery rate, FDR) <.05 (**Fig. S1A, B**).

Using these cut-offs, we found a good yield of recurrent ASM regions (**Fig. S3B**), but also many more loci with ASM seen in only one sample (**Fig. S4A**). We utilized such rare or “private” ASM loci for analyzing per-sample ASM frequencies, but for our downstream analyses focused on mechanisms and disease associations, we required ASM in at least two samples. Using these stringent criteria, in the combined SureSelect and WGBS dataset, after removing known imprinted loci (see below), we found 15,114 recurrent ASM DMRs, tagged by 17,935 index SNPs, representing 0.7% of all informative SNP-containing regions with adequate sequence coverage. These data are tabulated using the ASM index SNPs as unique identifiers, and annotated for strength of allelic methylation differences, presence or absence of ASM for each of the various types of samples, chromatin states, TF binding motifs, LD of the ASM index SNPs with GWAS peak SNPs, and other relevant parameters, in **Table S2**, with parameter definitions in Table S3.

### ASM in imprinted chromosomal regions

While this study focuses mainly on non-imprinted (genotype- or haplotype-dependent) ASM, genomic imprinting also produces ASM, due to parent-of-origin dependent DNA methylation affecting a small number of imprinted chromosomal domains (∼150 genes). Therefore, we used the GeneImprint database [24] and manual annotations from the literature to flag imprinted gene regions, many of which showed ASM in the SureSelect and WGBS data, thus serving as positive internal controls for ASM detection (**Table S4).** Since a hallmark of parent-of-origin dependent ASM (i.e. imprinting) is 50/50 allele switching between individuals, to test for possible novel imprinted loci, we assessed allele switching frequencies for all loci that showed ASM in non-cancer samples from 10 or more different individuals, after excluding known imprinted regions (Methods). The number of ASM DMRs decreases steeply when they are required to be found in many individuals (**Fig. S4**) because identifying such loci requires both a high number of informative individuals and highly recurrent ASM. Accordingly, among the non-cancer samples 326 ASM DMRs (corresponding to 371 index SNPs) outside of known imprinted regions were identified as showing significant ASM in more than 10 individuals. Only 13/326 (4%) of this group of DMRs showed allele switching at a frequency of greater than or equal to 20% of individuals. In comparison, among ASM DMRs identified in our dataset and located in or near known imprinted genes, a large majority (9/13; 70%) showed high frequency allele switching, with an approximately 50:50 ratio, as expected for parental imprinting. These results suggest that, as expected, most of the ASM loci identified by our genome-wide analysis reflect non-imprinted ASM. Interestingly, even among the small number of highly recurrent ASM loci with frequent allele switching in normal cells and tissues and located outside of validated imprinted domains, some (e.g. *IGF2R*, *IGF1R*) have been reported as imprinted in humans with inconsistent findings or variability. This small group of loci (**Table S5)** are not pursued further here but will be of interest for future testing of parent-of-origin dependent behavior.

### Validations by cross-platform comparisons and targeted methyl-seq

Consistency in the methylation profiles of genomic regions covered by both SureSelect and WGBS is shown in **Figure S3** series-wide and in **Figure S5** for single DNA samples analyzed by both methods. In addition, tracks of net methylation comparing both methods in the same biological sample revealed similar patterns in regions that were covered by both methods (**Fig. S6)**. In the overall series, within the fraction of the genome that was adequately covered by both methods and contained informative SNPs, we found 2,005 (49.1%) shared ASM “hits”. This substantial but partial overlap is expected, given that most ASM loci show a significant allelic methylation bias in some but not all individuals (**Table S2**). In addition, some ASM DMRs passed our stringent criteria in SureSelect but not in WGBS due to the greater sequencing depth of SureSelect in some regions. The pairwise correlation of allelic methylation difference between the 5 samples assessed both by SureSelect and WGBS showed that a majority (80%) of the 1203 “discordant” but adequately covered ASM SNPs were suggestive but sub-threshold, showing either less than 3 CpGs passing ASM criteria or a sub-threshold p-value due to lack of depth and spatial coverage, or an allelic methylation difference in the same direction with magnitude >10% but less than 20%.

To assess the true-positive rate for ASM calling, we selected 27 ASM DMRs, distributed through the range of ASM strength and confidence scores, for targeted bisulfite sequencing (bis-seq). As summarized in **Table S6**, this validation procedure confirmed the presence of ASM in two or more independent biological samples, outside of those utilized for the genome-wide series, with no discordance in the observed direction of the allelic methylation bias between the genome-wide methylation sequencing data and the targeted bis-seq, in 22 of the 27 DMRs assayed (examples in **Figures S7-S10**). For 4 of the 5 remaining loci, the presence of ASM was confirmed by targeted bis-seq using DNA from a sample (index case) that had shown ASM in the primary SureSelect or WGBS series. The single non-validated ASM DMR had a weak overall rank, but other examples in the lower tertile of ranks were validated (**Table S6**). This high overall validation rate, 81% using biological samples outside the primary genome-wide series, and 96% including independent technical validations on index samples from the primary series, indicates a high true-positive rate of the genome-wide data. Nonetheless, although the current dataset provides dense and reliable maps of ASM it is still non-saturating; with inclusion of more individuals and greater sequencing depth, more ASM DMRs will be identified - particularly those tagged by rare SNPs or having a narrow cell type-specificity or low ASM magnitude.

### ASM is increased in cancers due to widespread allele-specific CpG hypomethylation and focal allele-specific CpG hypermethylation in regions of poised chromatin

As shown in **Figure 1A**, when the numbers of ASM index SNPs per sample based on WGBS were normalized to the numbers of informative SNPs and the samples classified by normal vs. cancer status the number of ASM DMRs in the cancers overall (multiple myeloma, lymphoma, GBM) was on average 5-fold greater than in the overall group of non-neoplastic samples (Wilcoxon p=1.7×10-^07^). Among the cancers, the absolute mean frequencies of ASM were greater in the multiple myeloma and lymphoma cases and weaker in the GBMs. The differences in the frequency of ASM between cancer and non-cancer were greater, especially for the GBMs, when compared using the cell lineage-matched samples; non-neoplastic B cells for comparing to the B cell lymphomas and multiple myelomas, and non-neoplastic glial cells for comparing to the GBMs. Compared to these matched normal cell types, the average fold increase in ASM was 5-fold for multiple myeloma, 8.5-fold for the B cell lymphomas, and 9-fold for the GBMs.

**Figure 1.**
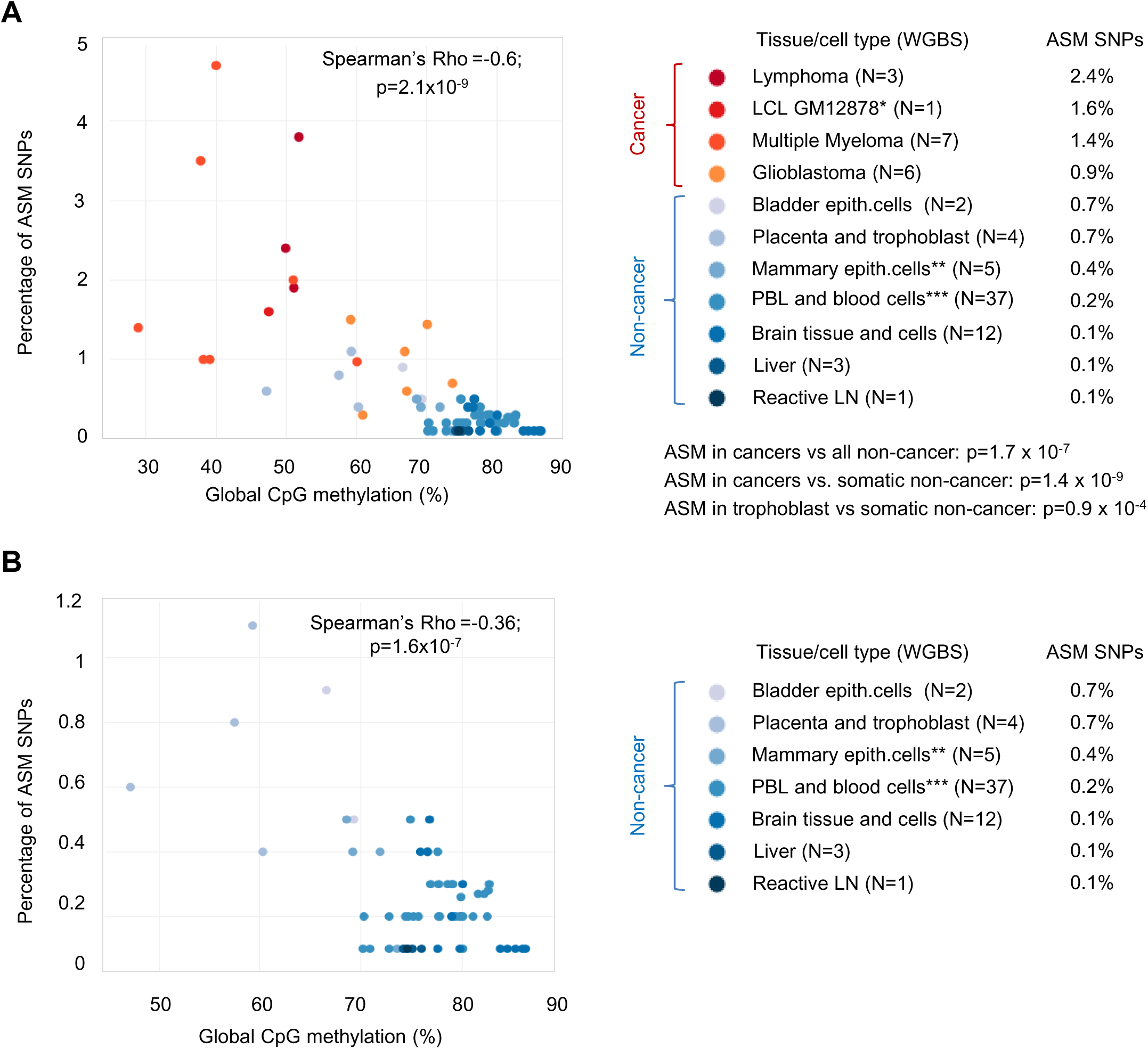
ASM is increased in cancers and correlates with global DNA hypomethylation. **A**, Relationship between global DNA methylation and the percentage of SNPs that reveal ASM in each sample, showing a strong inverse correlation between per sample ASM frequencies and global methylation levels. Cancer samples (color-coded in red scale) have higher ASM frequencies than nearly all the non-cancer samples (color-coded in blue scale), except for partial overlap of the GBM ASM frequencies with those of normal placental trophoblast and cultured bladder epithelial cells. When comparing the cancers to lineage-matched normal cell types, there is a 5 to 9-fold higher frequency of ASM in the cancers. The EBV-immortalized but euploid GM12878 LCL shows global hypomethylation and a high frequency of ASM. **B,** Zoomed-in graph showing that even among the non-cancer samples, those that are globally hypomethylated (i.e. placental trophoblast and cultured bladder epithelial cells) show relatively higher per-sample frequencies of ASM. P-values are from Wilcoxon tests. *EBV-immortalized GM12878 LCL is grouped with the cancers. **Mammary epithelial cell lines (N=3) and epithelium-rich normal breast tissue (N=2). ***Includes purified T cells, B cells and monocytes/macrophages (**Table S1**).

Since placental trophoblast is unique in having a cancer-like epigenomic and genomic profile [21–23], we also compared the per-sample ASM frequencies in cancers vs all normal somatic cells and tissues, excluding placenta and trophoblast, which revealed a 7-fold greater frequency of ASM in the cancers (Wilcoxon p=1.4×10^-9^), The EBV-transformed lymphoblastoid line (GM12878), which we had included to allow a direct reference to ENCODE data, showed a frequency of ASM in the mid-neoplasia range (**Fig. 1A**), which is important since much existing allele-specific mapping data, including expression and methylation quantitative trait loci (eQTLs, meQTLs) and allele-specific TF and CTCF binding by ChIP-seq (ASB) are from LCLs.

Given the well-known trend toward lower genome-wide (“global”) DNA methylation in human neoplasia (Gama-Sosa, Slagel et al. 1983, Feinberg, Gehrke et al. 1988), to evaluate mechanisms that could account for the gain of ASM in the cancers we asked whether there might be an inverse correlation between global methylation levels and frequencies of ASM. Global genomic hypomethylation was found in the GM12878 LCL and in the three types of primary cancers in our series (**Fig. 1A** and **Fig. S10**). As expected from prior studies by us and others [8, 21, 23], the placental tissue and purified trophoblast also showed global hypomethylation. Kernel density plots showed diffuse hypomethylation with nearly complete loss of the high methylation peak (fractional methylation >0.8) in lymphoma and myeloma compared to B cells, and a less dramatic but still significant hypomethylation in the GBMs compared to normal glia (**Fig. S11**).

Across the entire series of cancer and non-cancer samples, we found a strongly significant anti-correlation (i.e. inverse correlation) between per-sample ASM frequencies and global CpG methylation levels (Spearman’s Rho =-0.6; p=2.1×10^-9^). Importantly from a technical standpoint, this fundamental result was confirmed when we restricted our analysis to the WGBS data from a single sequencing facility using a single library preparation protocol (**Fig. S12A**). Arguing for global hypomethylation, not the malignant phenotype per se, as a main driving factor for increased ASM, the EBV-immortalized but euploid GM12878 LCL showed global hypomethylation and a high frequency of ASM, and among the non-neoplastic and non-immortalized samples, those that were relatively hypomethylated, namely the placental trophoblast and, to a lesser degree, the bladder epithelial cells that had been expanded in tissue culture, showed relatively higher per-sample frequencies of ASM (**Fig. 1B**).

To investigate how global hypomethylation could lead to increased ASM in cancers, we assessed the absolute and relative methylation levels of each of the two alleles across instances of ASM in the cancer samples, comparing myelomas and lymphomas to non-neoplastic B cells, and GBMs to normal glial cells. For each comparison, only the ASM-tagging index SNPs that were informative (heterozygous) in both cell types were considered, and we focused on loci showing ASM in the cancers but not in the cell lineage-matched non-neoplastic samples. We assessed the relative methylation levels of the low and high methylated alleles of these instances using a mixed linear model to estimate the average methylation level of each allele in each cell type taking into account the ASM magnitude in each cell type and the difference in ASM magnitude between cell types. As shown in **Figure 2** and **Figure S13,** this approach revealed that the average configuration was a relative loss of methylation (LOM) on one allele in the cancers. In 72% of cancer-only ASM occurrences in myelomas, 76% in lymphomas and 49% in GBMs, a strongly “hypermethylated/hypermethylated” configuration of the two alleles (“black/black”) in non-cancer became a “hypomethylated/hypermethylated” (“white-grey/black”) configuration in cancer (**Fig. 2**). The terminology here is a practical shorthand: “LOM” does not mean to imply that the normal cell types evolve into cancers; it is simply indicates the direction of the change in comparing the allelic methylation levels in the cancer vs cell lineage-matched non-cancer samples. Similarly, “cancer-only ASM” does not mean to imply that ASM at a given locus will never be detected in any non-cancer sample in future studies; it simply refers to the loci that have ASM in one or more cancer samples and in none of the non-cancer samples in the current dataset.

**Figure 2.**
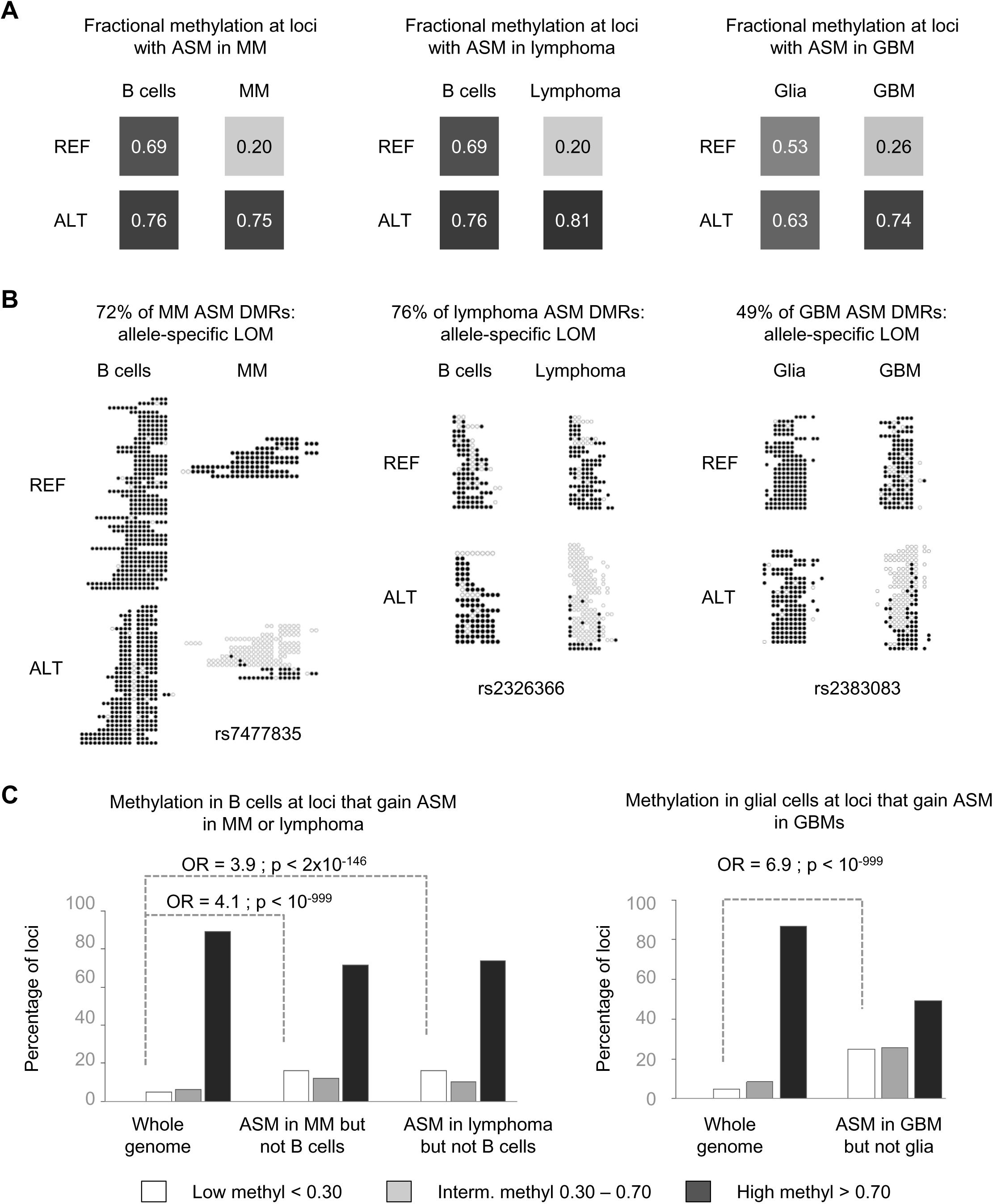
Gains of ASM in cancers due to widespread allele-specific LOM. **A**, Schematic showing the average configurations for allelic methylation levels in non-cancer and cancer samples at loci where ASM was observed only in cancer. Cancer samples are compared to the relevant non-cancer cell types. Average fractional methylation was estimated using a linear mixed model with random intercept and random slope (Methods). For each sample type, the squares represent the model estimate of the average fractional methylation in the low and high methylated alleles. **B**, Examples showing primary WGBS data. For the three types of cancers, the most frequent situation is an allele-specific LOM occurring in the cancers at loci that are highly methylated in the lineage-matched normal cell types. Rows are bisulfite sequence reads separated by REF and ALT allele. Methylated CpGs are black and unmethylated CpGs are white. **C**, Graphs showing distribution of net methylation in normal B-cells (left) and glia (right) grouped into 3 classes (low, intermediate and high methylation) at all informative CpGs (random expectation) and at CpGs where ASM was observed in the cancers but not in the matched non-cancer samples. While allele-specific LOM in cancer numerically accounts for most instances of cancer-only ASM (black bars; high methylation in the normal samples), relative to the background of global hypomethylation in the cancers it occurs less often than random expectation. In contrast, the smaller group of loci that have GOM leading to cancer-only ASM (white bars; low methylation in the normal samples) represent a significant enrichment over random expectation, given the globally hypomethylated genomic background of the cancers.

While the inverse correlation between per-sample ASM frequencies and global methylation is unequivocal, a multivariate regression analysis suggested that additional mechanisms might also be at play. This analysis showed that the anti-correlation between global methylation and per-sample ASM frequencies is partly independent of neoplastic status (p=9.4×10^-05^ after controlling for neoplastic status), and conversely, that the higher ASM frequencies in the cancers are only partly explained by global methylation levels (p=2.5×10^-04^ after controlling for methylation levels). In fact, while most of the cancer-only ASM loci conformed to the allele-specific LOM model, we found smaller but still substantial sets of loci (16% to 25% in the three cancer types) in which ASM in the cancers reflected allele-specific gains of methylation (GOM), relative to a biallelic low methylation configuration of the same regions in the lineage-paired normal samples (**Fig. 3** and **Fig. S14**).

**Figure 3.**
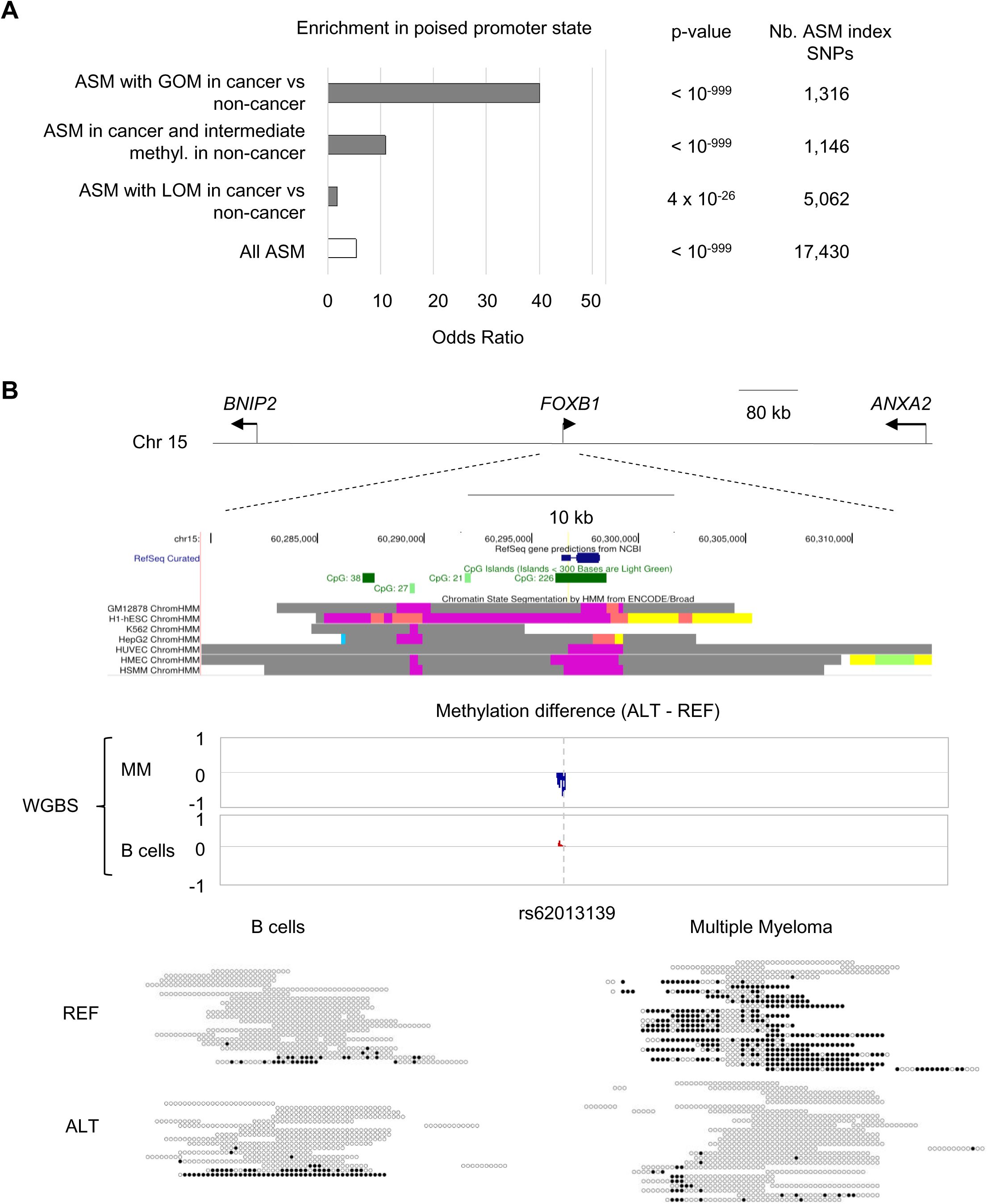
Gains of ASM in cancers due to allele-specific GOM at loci in poised chromatin. **A**, Graph showing enrichment in the poised promoter state as defined using ENCODE chromatin state segmentation by HMM. Although enrichment in poised promoter state is observed among ASM regions in general, this enrichment is dramatically increased among the subset of loci that show allele-specific GOM in cancers compared to cell lineage-matched normal samples. **B**, Map of the *FOXB1* locus showing an example of allele-specific GOM in multiple myeloma overlapping a CpG-island region with a poised promoter chromatin state (color coded purple). Methylation differences between alleles (index SNP rs62013139) are shown as a genome browser track and as WGBS reads for CD138+ multiple myeloma cells from a bone marrow aspirate, which show strong ASM with hypermethylation of the REF allele, and a paired peripheral blood non-neoplastic B cell sample from the same patient, which shows very weak ASM with slight hypermethylation of the ALT allele.

To further characterize this interesting set of loci with allele-specific GOM in the cancers, we compared the genomic and regulatory features among these loci to the background features of all informative loci using logistic regressions. As a comparison, we performed the same analyses for ASM loci that showed allele-specific losses of methylation in the cancers. This procedure revealed very strong over-representation of the poised “bivalent” promoter state among the ASM DMRs with allele-specific GOM in the cancers, compared both to ASM loci overall (O.R.=1.7; p=4.1×10^-26^) and to ASM loci with allele-specific LOM in the cancers (O.R.=40; p<10^-999^); **Fig. 3**. Poised promoters, as annotated by ENCODE chromatin state, are marked by the simultaneous presence of active histone marks, H3K4me3 and H3K4me2, and the repressive mark H3K27me3. Such regions are known to sometimes exist in a poised state in non-neoplastic stem cells [25] and can transition to a CpG-hypermethylated repressed state in cancer cells that acquire de-differentiated or stem cell-like phenotypes [26].

For completeness, using a similar statistical approach and mixed model for the set of ASM occurrences that were shared by cancer and non-cancer samples, we asked whether ASM might be not only more frequent in cancers, but also stronger. We found no significant differences in average ASM magnitude between the cancer and non-cancer ASM loci (**Fig S15**).

### Enrichment for chromatin states suggests mechanistic similarities between cancer and non-cancer ASM

Different chromatin states, and different classes of binding sites for TFs and CTCF, can be associated with specific patterns of CpG methylation [27–31]. Among the ASM DMRs found in the non-cancer samples, enrichment of active and poised promoter regions and enrichment of the poised/bivalent enhancer state are strong, the active transcription state is slightly enriched, and quiescent chromatin is depleted, relative to the background of adequately covered genomic regions (**Table 1**). This over-representation of promoter/enhancer elements among ASM DMRs suggests that ASM may contribute to inter-individual differences in gene expression – a conclusion that is supported by our observation of enrichment for eQTLs in ASM DMRs (**Table 1**). Using chromatin state data from the Roadmap Epigenomics project, which is available for T cells CD3, T cells CD4, T cells CD8, B cells, monocytes, cerebral cortex, and the GM12878 cell line, we tested chromatin state enrichment among ASM index SNPs separately in each of these tissues and cell types (as represented in our series, **Table S1**) and found that each of the major enrichments are shared across these tissues and cell types. This finding suggests that while ASM maps are partly tissue-specific the ASM is produced by shared underlying mechanisms.

Regarding our ability to address ASM tissue specificity in non-cancer samples, only SNPs informative in multiple tissues can be assessed. In addition, since ASM shows inter-individual variability, to correctly analyze ASM tissue specificity, each tissue needs to be informative in multiple heterozygous individuals. Using the ASM-by-cell type annotations (**Table S2**) we queried tissue specificity on the informative subset of 5,394 ASM SNPs where at least 3 samples per tissue (e.g. brain cells vs. blood cells, etc.) and at least 2 different tissues were informative. We found 1,519 ASM SNPs, representing 28% of the loci that could be tested, where the results suggest tissue specificity or at least some degree of tissue restriction. These findings need to be taken with caution since they are highly dependent on the number of informative samples. Also arguing for caution, when we performed pairwise correlations between ASM direction and magnitude using our set of paired samples from the same individuals (**Table S1**), the results suggested that some ostensibly tissue specific ASM DMRs in fact have a subthreshold allelic methylation bias, in the same direction, in the “non-ASM” tissue.

To assess similarities and differences in the characteristics of ASM in cancer vs. non-cancer we took two approaches: first, we tested for enrichment of chromatin states among ASM loci that were detected only in cancers (“cancer-only” ASM; observed in at least 2 cancer samples but in none of the non-neoplastic samples) and ASM loci detected in non-cancer samples (“normal ASM”; present in at least one non-cancer sample, but allowing ASM in cancers as well), separately using bivariate logistic regression and second, testing the differential enrichment between the two groups using multivariate regressions including the interaction term between ASM and cancer status. Both approaches showed that ASM DMRs in cancer and non-cancer show a parallel enrichment in all the strongly enriched chromatin states, albeit with some differences among the less strongly enriched features (**Table 1**). These findings suggest that the mechanisms leading to ASM are at least partly similar in non-neoplastic and neoplastic cells - a conclusion that is further supported by analysis of correlations of ASM with SNPs in CTCF and TF binding sites, described below.

### ASM correlates with allele-specific binding affinities of specific CTCF and TF recognition motifs in both cancer and normal samples

The hypothesis that allele-specific TF binding site occupancy (ASB) due to sequence variants in regulatory elements could be a mechanism leading to ASM has been supported by previous data from us and others [8, 10, 11]. To test this hypothesis using denser maps, and to ask whether this mechanism might underlie ASM in both normal and neoplastic cells, we analyzed the set of ASM loci for enrichment of sequence motifs recognized by classical TFs, and motifs recognized by CTCF, which defines the insulator chromatin state and regulates chromatin looping [32–34]. Previously we showed that ASM DMRs can overlap with strong CTCF ChIP-seq peaks and polymorphic CTCF binding sites [8, 35]. In our expanded dataset, we used atSNP to identify CTCF motif occurrences where the ASM index SNP not only overlaps a CTCF motif but also significantly affects the predicted binding affinity, as reflected in the Position Weight Matrix (PWM) score. For this analysis we required a significant difference in binding likelihood between the two alleles (FDR <0.05) and a significant binding likelihood (p <0.005) for at least one of the alleles (reflecting potential CTCF occupancy of at least one allele). We identified 3.075 ASM SNPs (17%) that significantly disrupted at least one of the canonical or ENCODE-discovery CTCF motifs [33, 36]. To estimate the random expectation of polymorphic CTCF motif occurrences in the genome (the background frequency), we ran atSNP on a random sample of 40,000 non-ASM informative SNPs (1:3 ASM vs non-ASM SNP ratio) and found that 8.4% of these non-ASM informative SNPs significantly disrupted a CTCF motif, corresponding to a substantial enrichment for disrupted CTCF motif among ASM SNPs (OR=2.3; p-value=10^-218^).

As noted in our previous smaller study [8], in the enlarged dataset this overall enrichment of CTCF motif-disrupting SNPs among ASM loci persists, albeit slightly weaker, when considering only non-CpG-containing polymorphic CTCF motif instances (OR=1.8; p=6.3×10^-34^). When testing enrichment separately for each of the 14 distinct ENCODE/JASPAR-defined CTCF motifs, we found significant enrichment for 13 of them (**Table S7**). Moreover, as shown in **Figure 4, Figure S16,** and **Table S8**, the difference in binding affinity score between alleles is significantly anti-correlated (i.e. inversely correlated) with the difference in methylation for three of these motifs, and these correlations persist after adjustment for the presence or absence of CpGs in the motif occurrences in a multivariate model. Thus, consistent with our previous conclusions in the smaller dataset, which required motif pooling [8], these results from individual motif classes show that the presence of a methylatable CpG in the CTCF binding motif is not required; rather, the essential feature is allele specific binding site occupancy.

**Figure 4.**
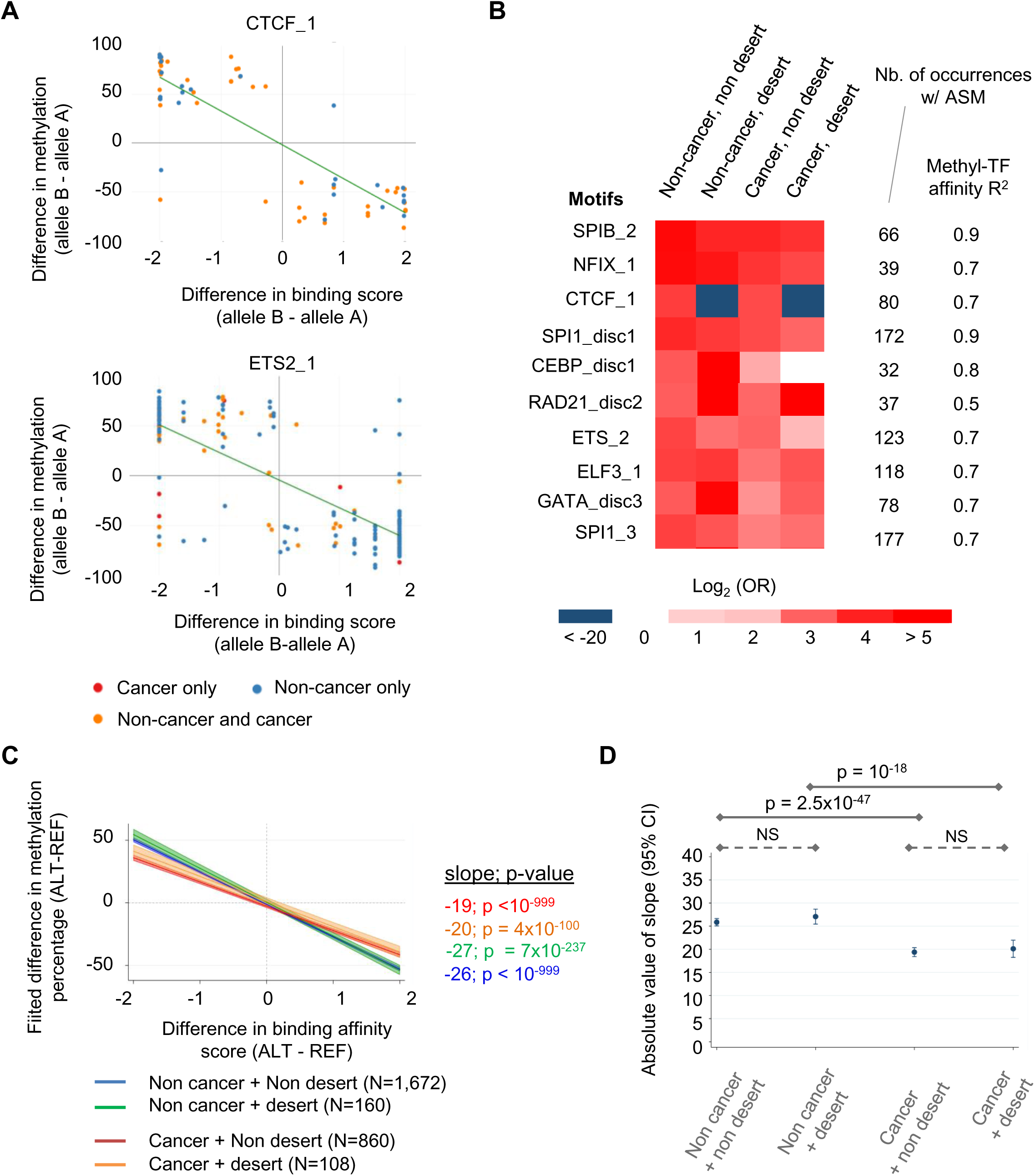
ASM is driven by allele specific CTCF and TF binding in both normal and neoplastic cells. **A**, X-Y plots showing examples of TF motifs with strong correlations between predicted allele-specific binding site affinities (estimated by PWM scores) and methylation differences across all occurrences showing ASM. These examples are among 179 significantly correlated motifs, listed in **Table S8**. Each data point represents one occurrence of the motif overlapping an ASM index SNP in cancer (orange) or non-cancer samples (blue). For occurrences showing ASM in multiple samples, allelic methylation differences were averaged across samples by sample type. R^2^ and B-H corrected p-values (FDR) were calculated using linear regression. **B,** A large majority of the polymorphic motifs with significant correlations between allelic methylation and predicted binding affinities are also statistically enriched among ASM regions (**Table S9**). The heatmap shows the enrichment or depletion, in log^2^(O.R.), for the top 10 enriched TF binding motifs among cancer or non-cancer ASM loci in regions defined as chromatin desert or non-desert (Methods). **C,** Significant correlations between allelic TF binding affinity scores and ASM in each of the 4 classes of ASM loci. The graph shows the fitted ASM difference on PWM score using a multivariate mixed model. The fitted line and its 95-confidence intervals (area) are shown for each ASM class; slopes were calculated by the marginal effects of the interaction term between PWM score and ASM class and were significantly different from zero. The correlations are similar in cancer ASM (in both non-desert and desert) compared to non-cancer ASM, with only small differences in the slopes for each class. **D,** Pairwise comparisons of the correlations in each of the 4 classes of ASM loci, Bonferroni-adjusted for multiple testing. While all the slopes are in a similar range, the correlations in the mixed model are weakest for cancer-only ASM loci, with a modest but statistically significant difference between the cancer vs non-cancer ASM classes, but not between desert and non-desert ASM loci. N: number of occurrences included in the mixed model.

Like CTCF, classical TFs could account for instances of ASM via ASB. When we scanned each ASM SNP for all ENCODE/JASPAR defined TF motifs [37], we found 17,022 (95%) recurrent ASM SNPs disrupting at least one TF binding site occurrence, compared to 32,790 (82%) in a randomized non-ASM background set (OR=3.9, p<10^-999^). Of these ASM SNPs, 12,853 overlapped at least one ENCODE DNase I hypersensitive site and 3,044 at least one ENCODE cognate ChIP-seq TF peak. From a panel of 2,263 TF motifs with at least 10 occurrences, we found 856 motifs with a specific enrichment (OR ≥ 2 and FDR corrected q-value<0.05, compared to the random sample of 40,000 non-ASM informative SNPs) among ASM DMRs (**Table S7**). Next, using linear regression of allele-specific affinity (PWM) score differences on allele-specific CpG methylation differences, we found 177 TF binding motifs, corresponding to 115 cognate TFs, where DNA methylation appears to be shaped by binding site occupancies (**Fig. 4, Fig. S14,** and **Tables S8** and **S9**). Among these motifs, 144 also showed significant enrichment among ASM loci (**Table S9**). Regarding the motif classes that are enriched among ASM index SNPs but do not show statistically significant correlations of PWM scores with ASM magnitude, it is likely that some simply have too few ASM occurrences in the current dataset to achieve significance in the correlation analysis.

Using stringent statistical criteria (FDR<0.05 and R^2^≥ 0.4), all but two of the TF motifs that were correlated with ASM show inversely correlated behavior, such that a relatively higher binding likelihood (stronger PWM score) correlates with CpG hypomethylation (**Table S8**, examples in **Figure 4** and **Figures S14 and S15).** Multivariate linear regression of the 158 (out of 177) significantly correlated motifs with at least three CpG-containing and three non-CpG-containing occurrences revealed that these inverse correlations between binding affinity scores and methylation levels persist after adjustment for the presence or absence of CpGs in the motifs. Like the findings for CTCF sites, these results suggest that ASM regions form around polymorphic TF binding sites because of allele-specific differences in binding site occupancy (ASB), not requiring a methylatable CpG in the binding motif.

Lastly and importantly, we tested for enrichment of TF and CTCF binding motifs and correlations of ASM with predicted binding affinities separately in the sets of ASM loci that were detected only in the cancers (including the GM12878 LCL) vs those found in the total group of non-cancer samples. We also analyzed the full set of ASM loci using a multivariate mixed model to test for interactions of normal vs cancer status with the TF binding site affinity to ASM strength correlations. The results showed that ASM loci in cancer and non-cancer samples have similar directions of the correlations of ASM with disruptive SNPs in the top-ranked classes of polymorphic TF binding motifs (**Fig. 4** and **Fig. S16**), which indicates sharing of this fundamental mechanism of ASM in normal and cancer cells. This key result was confirmed when we restricted our analyses to the data from a single sequencing facility using a single library construction method (**Fig. S12B, C**). However, the correlations between predicted TF binding site affinities and ASM amplitude were slightly weaker on average (shallower slope in the X-Y plot) among the cancer-only ASM loci (**Fig. 4** and **Fig. S16)**, and ASM-correlated motifs were not enriched among these loci **(Table 1).** These findings are explained by the presence of a subgroup of cancer-only ASM loci that show allele-switching (see below).

### Direct testing of the TF binding site occupancy mechanism of ASM

As a crucial validation, using our GM12878 SureSelect and WGBS data and the large number of ENCODE ChIP-seq experiments available for this cell line, we could directly ask whether ASM regions with or without polymorphic CTCF and classical TF binding sites exhibit allele specific binding of the cognate factors. Among the 2,102 high-confidence ASM index SNPs from our GM12878 data, 787 overlapped at least one ChIP-seq peak in this cell line and had enough ChIP-seq reads (>10X) to assess allele-specific binding of at least one ENCODE-queried TF. We found that 16.6% (131) of these ASM index SNPs showed ASB for at least one TF that could be assessed using available ENCODE data. As predicted from the binding site occupancy hypothesis for ASM, at 100 (76%) of these sites, considering both CTCF and TF motifs, the hypomethylated allele showed significantly greater occupancy. This percentage far exceeds random expectation (exact binomial test, p=1.2e10^-9^). Confirming this pooled analysis, among 9 TFs with more than 10 ASB occurrences associated with ASM, 7 examples, including the ELF1 (ETS-family) motif and others, showed a significant enrichment in ASM occurrences with an inverse correlation of predicted binding affinity with allelic CpG methylation (ASB-ASM instances with inverse correlation: 90%-100%, FDR <0.05).

### Somatic mutations in TF binding sites can produce ASM in human cancers

To more completely understand the features of ASM in cancers, and to further test the hypothesis that disruptive SNPs in TF binding motifs give rise to ASM, we searched for somatic mutations in the 4 multiple myeloma cases that were paired with non-neoplastic peripheral blood B cells from the same patients (analyzed using the same WGBS library protocol to ensure similar regional coverage depth) which served as germline reference sequences. We found somatic mutations at frequencies of 499 to 1,023 per case, and among these mutations from 6% to 17% were associated with gains of ASM (referred to here as “de novo ASM”, examples in **Fig. 5**). We next filtered out mutations situated within 1kb of known ASM index SNPs that had already been seen in other samples, since such instances might simply be uncovering normal ASM by conferring heterozygosity in regions that were non-informative in the patient’s germline sequence. Using the filtered list (410 de novo ASM occurrences), we asked whether the somatic mutations associated with de novo ASM might be disrupting TF binding motifs at a frequency greater than random expectation. We found a significant enrichment (O.R. >2 and p<0.05) for 54 TF binding motifs among the de novo ASM occurrences, compared to the representation of these motifs among all the somatic mutations that were not associated with ASM. Even more convincingly, we found that a majority (71%) of the TF binding site motif classes that were enriched among instances of de novo ASM belonged to the same motif classes that were enriched among the much larger set of ASM loci that were tagged by germline SNPs (O.R.=3.6, p-value=6.3×10^-4^; examples in **Fig. 5**). Thus, while mutation-associated de novo ASM does not make a large numerical contribution to the overall gains of ASM in cancer vs. non-cancer, this special phenomenon is informative in emphasizing the shared underlying mechanism (TF binding motif disruption or creation) for ASM in cancer and normal cells.

**Figure 5.**
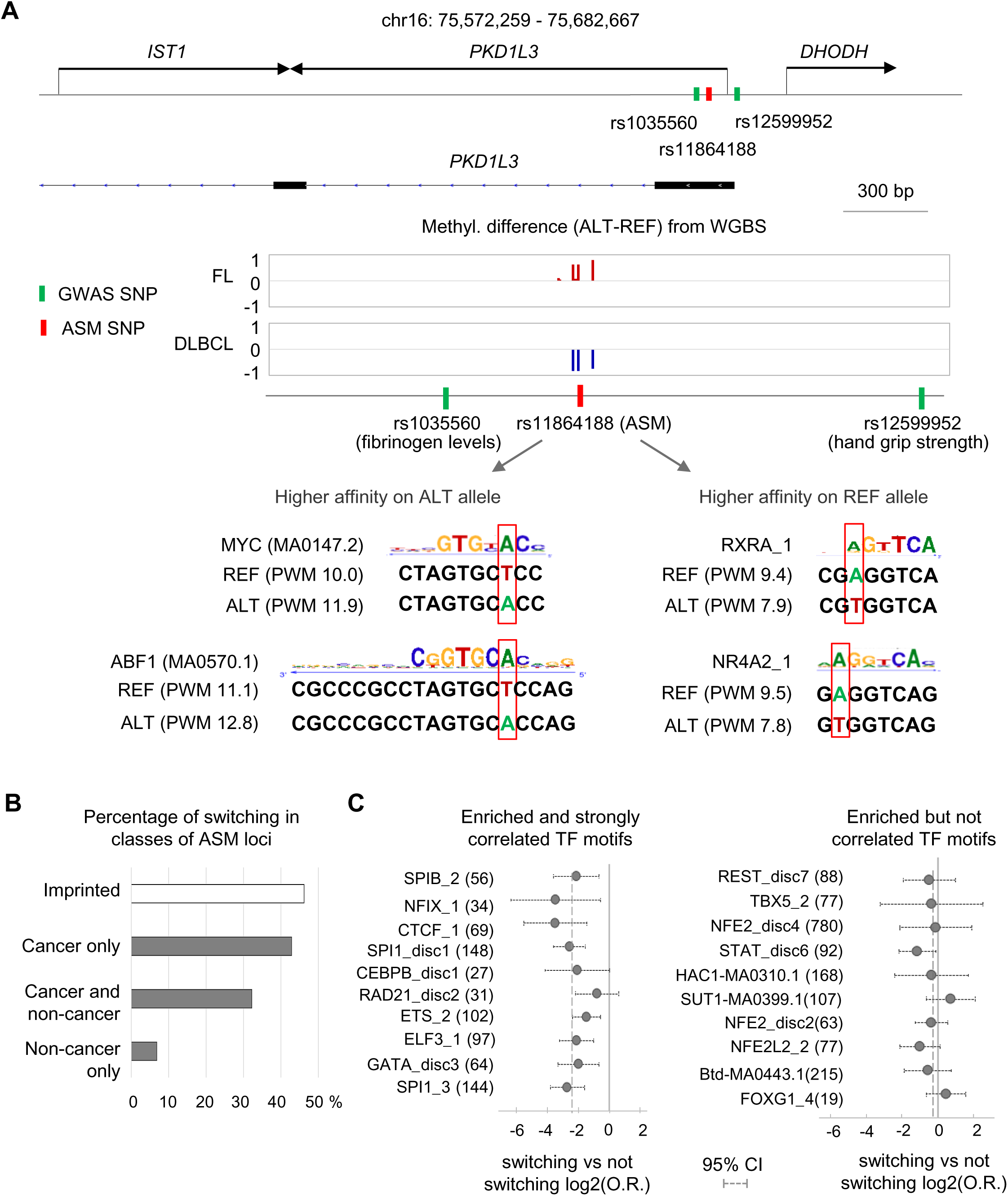
Examples of de novo ASM associated with somatic mutations in cancers. **A**, Map of the de novo ASM region centered on a somatic mutation at chr2:128663542 in case SUB006. The mutation is intergenic between *AMMECR1L* and *SAP130.* It overlaps a poised chromatin region and creates a de novo binding motif for the MAF TF, with a higher binding affinity on the mutated allele than on the reference (germline) allele. The graphical representation of the WGBS data shows strong ASM with less methylation on the mutated allele. The reference allele shows similar methylation (high) in both the myeloma and paired B cells. **B**, Map of the de novo ASM region centered on a somatic mutation at chr11:12692969, also in case SUB006. The mutation is intergenic downstream of *PARVA* and upstream of *TEAD1.* It overlaps an active enhancer chromatin region and creates a de novo binding motif for HOXA4 with a higher binding affinity on the mutated allele than on the reference (germline) allele. The graphical representation of the WGBS data shows strong ASM with lower methylation on the mutated allele. The reference allele showed a similar methylation (high) in both the myeloma and paired B cells. For these two motifs, the occurrences map to the negative strand and are oriented 3’ to 5’ per atSNP convention.

### ASM DMRs are found both in active chromatin and in quiescent “chromatin deserts”

For post-GWAS mapping of rSNPs much attention has been appropriately focused on cataloguing SNPs that are expression quantitative trait loci (eQTLs) and/or lie within regions of ASB. Such efforts are aided by databases such as AlleleDB for allele-specific marks [38–40], and RegulomeDB [41, 42], which highlights potential rSNPs in non-coding regions by assigning a score to each SNP based on criteria including location in regions of DNAase hypersensitivity, binding sites for TFs, and promoter/enhancer regions that regulate transcription. Our cross-tabulations indicate that, despite a strong enrichment in ASB SNPs among ASM index SNPs (**Table 1**), most of the ASM index SNPs (>95%) in our expanded dataset currently lack ASB annotations (**Table S2**). In addition, index SNPs for strong ASM DMRs sometimes have weak RegulomeDB scores (**Table S2**). Thus, from a practical standpoint with existing public databases, ASM mapping for identifying rSNPs appears to be largely non-redundant with other post-GWAS modalities.

To further assess the unique value of ASM mapping, we defined “chromatin desert” ASM regions as 1 kb genomic windows, centered on ASM index SNPs, that contained no DNAse peaks or only one DNAse peak among the 122 ENCODE cell lines and tissues, and no strong active promoter/enhancer, poised, or insulator chromatin state in any ENCODE sample. Less than 55% of such regions have SNPs listed in RegulomeDB, and when they are in that database they almost always (93%) have weak scores equal to or greater than 5 (**Table S2**). While most ASM loci map to active chromatin and are depleted in desert regions overall (**Table 1**), we find that 8% of ASM index SNPs in normal cells and 22% of cancer-only ASM SNPs are in chromatin deserts (**Table 1** and **Table S2**). Although deserts lack evidence of TF and CTCF binding in available databases, ASM DMRs found in these regions might be informative for localizing bona fide rSNPs if some desert regions contain cryptic binding motifs that were active (occupied) at an earlier point in the history of the cell.

To address this possibility, we asked whether correlations of ASM with disruptive SNPs in TF binding motifs might also pertain to ASM in desert regions. We analyzed the full set of ASM loci using a multivariate mixed model to test for interactions of normal vs cancer status and desert vs non-desert location (i.e. 4 classes of ASM loci) with the TF binding site affinity to ASM strength correlations. Some motifs, such as CTCF binding sites, were highly depleted in deserts and therefore excluded from the analysis, which was performed on the subset of 74TF motifs that had at least three occurrences per ASM class. The correlations, when significant (FDR<0.05), were in the same direction (inverse correlation of predicted binding affinity with allelic methylation) in all ASM classes. As expected from the findings above, we observed a slightly weaker correlation for canceronly ASM loci compared to ASM loci in non-cancer samples. However, no differences in the strength of the correlations were found when comparing ASM occurrences in desert versus non-desert locations, both for normal and cancer-associated ASM loci. The simplest hypothesis to explain these results is that ASM DMRs in desert regions are footprints left by rSNPs that disrupt cryptic TF binding sites that were active at some stage of normal or neoplastic cell differentiation (or de-differentiation) but are no longer active in available cells or tissue types. **Figure S17** shows examples of ASM DMRs in desert regions that contain disruptive SNPs in ASM-correlated ETS- and ERG-family TF binding motifs.

### Allele-switching at ASM loci is infrequent in normal samples but increased in cancers

Most of the ASM DMRs passed statistical cutoffs for ASM in less than half of the informative samples (**Table S2**), with variability not only between cell types and cancer status but also within a single cell type. Given the connection between TF binding site occupancies and ASM, one hypothesis to explain this variability invokes differences in intracellular levels of TFs (**Fig. S18A**). Alternatively, genetic differences (i.e. haplotype effects due to the influence of other SNPs near the ASM index SNP) could also play a role (e.g. **Fig. S18B)**. A more extreme form of variation was observed at some ASM loci, namely “allele switching” [8], in which some individuals have relative hypermethylation of Allele A while others show hypermethylation of Allele B, when assessed using a single index SNP. Some instances of allele switching reflect haplotype effects [8] or parental imprinting, but other occurrences might have other explanations. In this regard, a striking finding in the current dataset is that the frequency of allele switching among ASM loci in normal samples is low (14%), while the rate of allele switching is strikingly higher (43%) among cancer-only ASM loci (**Fig. 6A, B** and **Fig. S18C**). This finding suggests that biological states, here neoplastic vs non-neoplastic, can influence the stability of ASM, with greater epigenetic variability or instability in the cancers.

**Figure 6.**
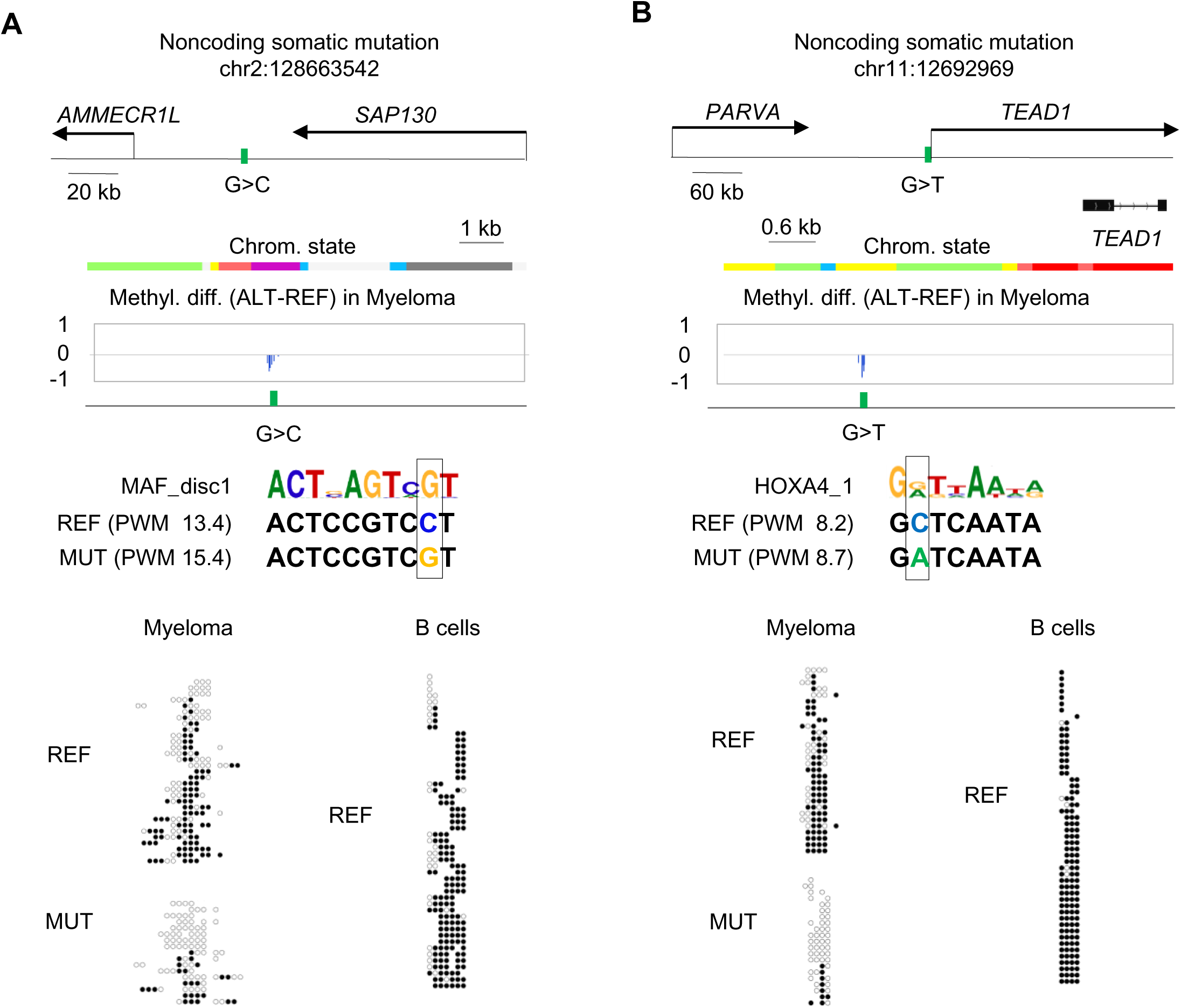
Increased allele switching at ASM loci in cancers. **A**, Map of the ASM region tagged by index SNP rs11864188 in the *PKD1L3* gene. The ASM shows switching with the ALT allele being hyper-methylated in the FL but with an opposite direction of the allelic methylation bias in the DLBCL. No ASM was detected in 23 non-cancer heterozygous samples. The rs11864188 SNP disrupts multiple TF motifs, some of which have opposite allele-specific predicted affinity differences. These motifs include MYC and ABF1 motifs with a higher affinity on the ALT allele and RXRA and NR4A2 motifs with a higher affinity on the REF allele. For this occurrence, the MYC motif maps the negative strand and is oriented 3’ to 5’. **B**, Frequency of switching, scored for ASM index SNPs that were heterozygous in at least 2 samples, is increased among cancer-only ASM loci (43%) compared to ASM loci found in normal samples (10%). As an internal control, informative ASM index SNPs mapping to imprinted regions showed 46% switching, approximating the expected 50% based on parent-of-origin dependent ASM. **C**, Enrichment analyses of CTCF and TF binding motifs among switching vs non-switching ASM loci: in the left panel, polymorphic motifs with very strong correlations between ASM magnitude and affinity score differences are found to be depleted among switching compared to non-switching ASM loci. In the right panel, polymorphic TF motifs enriched among ASM but with little or no correlations of predicted binding affinity with ASM magnitude showed little or no depletion among switching ASM loci. The solid vertical lines represent zero (no enrichment or depletion), and the dotted lines show the average values for depletion for each set of motifs.

To investigate this variability, we compared the features of ASM DMRs that showed allele switching versus those that did not. As shown in **Figure 6C**, the sets of ASM index SNPs for two classes of loci differed significantly in the relative representation of specific CTCF and TF binding motifs, such that the CTCF_1 motif and nearly all of the most strongly ASM-correlated classical TF binding motifs were markedly under-represented among the switching loci. Reinforcing this finding, ASM loci that were highly recurrent across multiple normal cell types and individuals showed a low frequency of switching, even when these loci had ASM in some cancers (**Fig. S19**).

These results suggest a working model that postulates two classes of binding motif occurrences. One group of motif occurrences stably bind their cognate TFs when the motif sequence is optimal but are sensitive to the effects of disruptive SNPs in the motif. These motif occurrences therefore show strong unidirectional correlations of PWM scores with ASM, independently of the neoplastic cellular phenotype. Another group of motif occurrences are postulated to have more labile binding of their cognate TFs, are sensitive to changes in the intracellular levels of their cognate factors, and can participate in ASM allele switching, via “TF competition”. According to this model (**Fig. S18C**), in situations with adequate chromatin accessibility, there can be replacement of one TF by another more highly expressed one that recognizes a nearby or overlapping DNA sequence motif. The credibility of this hypothesis is supported by the well-known over-expression of various oncogenic TFs in cancer cells, and by data indicating that global DNA hypomethylation in transformed cells is associated with increased chromatin accessibility at regulatory elements [43, 44].

### ASM index SNPs in LD or precisely coinciding with GWAS peak SNPs

For assessing the value of ASM as a signpost for rSNPs in disease-associated chromosomal regions, we defined lenient and stringent haplotype blocks by applying the algorithm of Gabriel et al [45], using 1000 Genomes data and employing D-prime (D’) values, both with standard settings utilizing high D’ and R-squared (R^2^) values to define “stringent” blocks (median size 5 kb) and with relaxed R^2^ criteria to define larger “lenient” blocks with a median size of 46 kb (**Figs. S20 and S21**). We also calculated R^2^ between each ASM index SNP and GWAS peak SNP to identify SNPs in the same haplotype block and with high R^2^, plus SNPs in strong LD located in genomic regions that lacked a haplotype block structure. We took this two-fold approach because (i) R^2^ can fail to identify SNPs in perfect LD when rare mutations have occurred over time on pre-existing common alleles in the population – a situation that can have a high D’, and (ii) for some loci, the combined effect of multiple regulatory SNPs, some with weak R^2^ values but high D’, might be responsible for net effects on disease susceptibility. Using our complete list of ASM DMRs and GWAS data from NHGRI-EBI, including both supra-threshold and suggestive peaks (p<10^-6^), we identified 1,842 ASM SNPs in strong LD (R^2^>0.8) or precisely coinciding with GWAS peak SNPs (**Table S2**). Highlighting mechanistic information from these ASM loci, among the ASM index SNPs in strong LD or precisely coinciding with GWAS peak SNPs, 1,450 disrupted ASM-enriched classes of CTCF or TF binding motifs and 310 disrupted significantly ASM-correlated CTCF or TF binding motifs.

#### ASM index SNPs in LD with GWAS peaks for autoimmune/inflammatory diseases

We found 275 ASM DMRs containing 305 index SNPs in strong LD (R^2^>.8) with GWAS peak SNPs for autoimmune and inflammatory diseases, or related traits such as leukocyte counts (**Table S10**) which corresponds to a moderate but significant enrichment (O.R.=1.5, p=1.2×10^-18^). Of these 305 index SNPs, 8 were in HLA genes and the remainder were in non-HLA loci. About half of these regions showed ASM in immune system cell types and/or B cell tumors (**Table S10**). Among these ASM index SNPs, 66 precisely coincided with GWAS peak SNPs, supporting the candidacy of these statistically identified SNPs as functional rSNPs. Moreover, 61 of the 305 ASM index SNPs altered strongly ASM-correlated CTCF or TF binding motifs, and 237 disrupted enriched classes of binding sites, thus providing mechanistic leads to disease-associated transcriptional pathways (**Tables S2** and **S10**). Interesting ASM index SNPs for this disease category, some precisely coinciding with GWAS peak SNPs and others in strong LD with these peaks include rs2145623, precisely coinciding with a GWAS peak SNP for ulcerative colitis, sclerosing cholangitis, ankylosing spondylitis, psoriasis and Crohn’s disease (nearest genes *PSMA6*, *NFKBIA*), rs840015 in strong LD with GWAS peak SNPs for celiac disease, rheumatoid arthritis, and hypothyroidism (nearest genes *POU2F1* and *CD247*), rs10411630 linked to multiple sclerosis (MS) via LD with GWAS peak SNP rs2303759 (nearest genes *TEAD2*, *DKKL1*, and *CCDC155*; **Fig. S7**), rs2272697 linked to MS via LD with GWAS peak SNP rs7665090 (nearest genes *NFKB1*, *MANBA*), rs2664280 linked to inflammatory bowel disease, systemic lupus erythematosus (SLE) and psoriasis via GWAS SNPs rs2675662 and rs2633310 (nearest genes *CAMK2G*, *PLAU*, *C10orf55*; **Fig. 7**), and rs6603785 which precisely coincides with a GWAS peak for SLE and hypothyroidism (nearest genes *TFNRSF4*, *SDF4*, *B3GALT6*, *FAM132A*, *UBE2J2*; **Fig. S22**). Each of these ASM index SNPs disrupts one or more strongly ASM-enriched or ASM-correlated TF binding motifs (**Table S10**).

**Figure 7.**
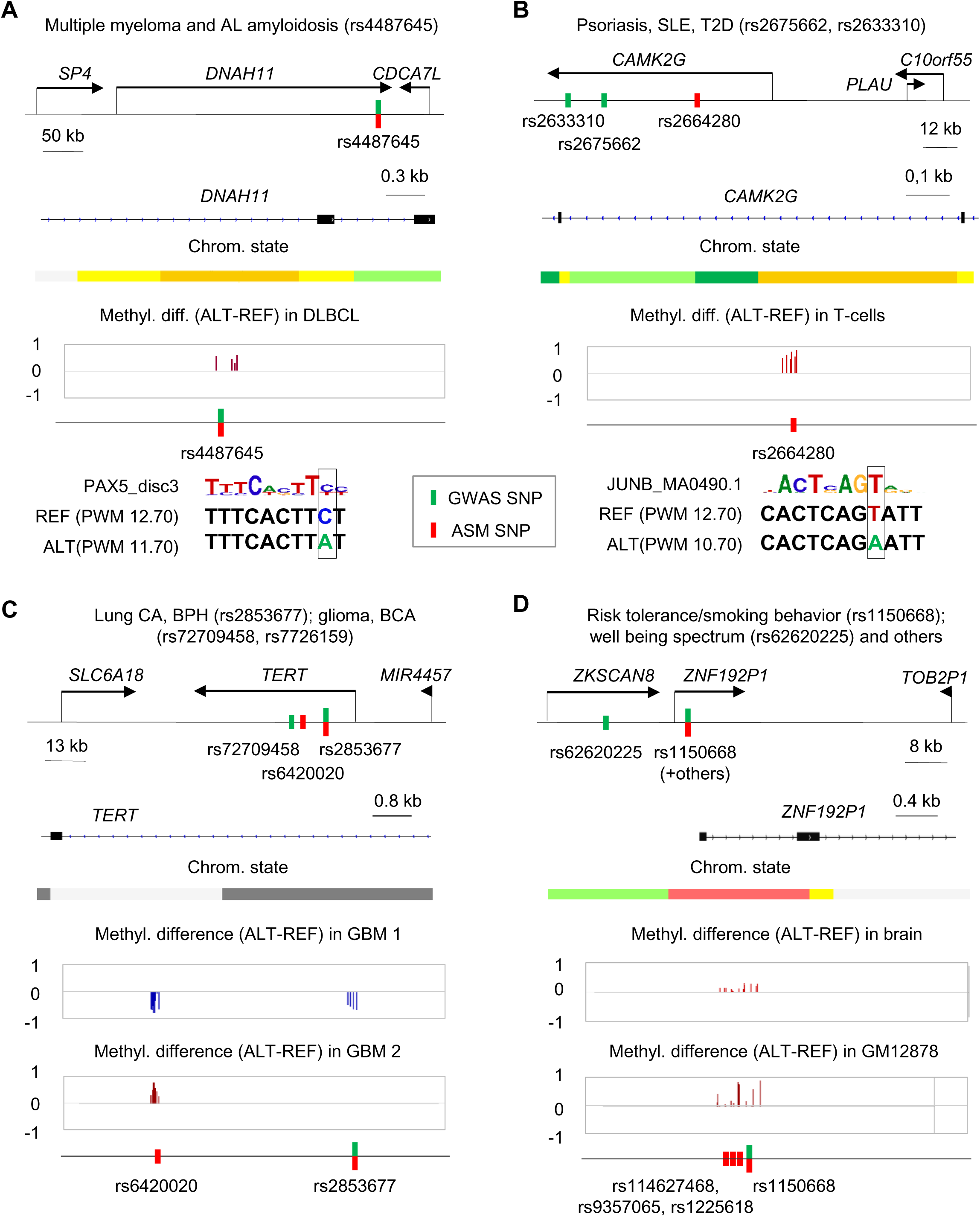
Examples of ASM index SNPs in strong LD or precisely coinciding with GWAS peaks. **A**, Map of the ASM DMR tagged by index SNP rs4487645, coinciding with a GWAS peak SNP for multiple myeloma (p=3.0×10^-14^; O.R.=1.38) and AL amyloidosis (p=2.0×10^-9^; O.R.=1.35). The ASM index SNP is in an enhancer region (yellow-coded chromatin state; GM12878 track) of the *DNAH11* gene on chromosome 7. This SNP disrupts a PAX5_disc3 TF binding motif on the ALT allele. The REF allele, with intact high-affinity motif, is relatively hypomethylated, as predicted by the TF binding site occupancy model. Additional motifs in this DMR are in **Table S2**. **B**, Map of the ASM region tagged by index SNP rs2664280, in strong LD with GWAS peak SNPs rs2675662 for psoriasis (p=3.0×10^-8^; O.R.=1.14) and rs2633310 for T2D (p=2.0×10^-8^; Beta=-.044). The ASM index SNP is in an enhancer region (yellow-coded chromatin state; GM12878) in the *CAMK2G* gene on chromosome 10. This SNP disrupts several AP1 binding motifs (JUNB shown) on the ALT allele, with higher binding affinity on the REF allele, which is relatively hypomethylated as predicted. For this occurrence, the motif maps the negative strand and is reported from 3’ to 5’. Additional motifs in this DMR are in **Table S2**. **C,** Map of the ASM DMRs tagged by the rs2853677 and rs6420020 index SNPs in the *TERT* gene on chromosome 5. The DMRs are in chromatin that is quiescent/repressed in most ENCODE samples (light and dark gray; K562 track), but this region is transcribed in undifferentiated H1-hESC. ASM for these DMRs was found only in cancer samples (GBMs). The index SNP rs2853677 is a GWAS peak SNP for non-small cell lung cancer and benign prostatic hyperplasia (p<10^-999^; O.R.=1.41 and p=2.0×10^-22^; O.R.=1.12 respectively). The other ASM index SNP, rs6420020 is in LD with GWAS peak SNPs for GBM (p=6.0×10^-24^; O.R.=1.68), breast carcinoma (p=3.0×10^-8^; O.R.=1.07), and chronic lymphocytic leukemia (p=6.0×10^-10^; O.R.=1.18). ASM allele switching is seen at rs6420020; candidate polymorphic TF binding motifs are in **Table S2**. **D,** Map of the ASM DMR tagged by multiple SNPs (rs114627468, rs9357065, rs1225618 and rs1150668) in the promoter region of the *ZNF192P1* pseudogene, flanked by coding genes in the *ZSCAN* family, on chromosome 6. ASM index SNP rs1150668 is a GWAS peak SNP for body height (p=2.0×10^-7^; Beta=-.060), smoking status (p=6.0×10^-15^; Beta=-.0086), smoking behavior (p=3.0×10^-8^; Beta=+.011), myopia (p=1.0×10^-11^; Beta=+6.78), and schizophrenia with autism spectrum disorder (p=8.0×10^-11^; O.R.=1.07). In addition, the 4 ASM index SNPs are in a stringently defined haplotype block containing GWAS peak SNP rs62620225, for multiple phenotypes including wellbeing spectrum (p=6×10^-12^; Beta=0.023). ASM in this DMR was observed in multiple tissues, including brain. The ASM index SNP rs1225618 is as an ASB SNP for TAF1; other ASM-correlated motifs disrupted by the index SNPs are in **Table S12**. Additional examples of disease-linked ASM loci are in **Figures S7-S9**, **S22**, and **S23**.

#### ASM index SNPs in LD with GWAS peaks for cancer susceptibility

We found 247 ASM DMRs containing 268 index SNPs in strong LD (R^2^>.8) with GWAS peak SNPs for cancer susceptibility or response to treatment (**Table S11**), which represents a moderate but significant enrichment (O.R.=1.5, p=3.2×10^-22^). Among these loci a large majority showed ASM in cancers or cell types that approximate cancer precursor cells (e.g. B cells for lymphoma and multiple myeloma, glia for GBM, mammary or bladder epithelial cells for carcinomas, normal liver for hepatocellular carcinoma, etc.) and/or in T cells, which are relevant to cancer via immune surveillance. In these DMRs, 60 of the ASM index SNPs precisely coincided with the GWAS peak SNPs, supporting the candidacy of these statistically identified SNPs as functional rSNPs. Among the 268 ASM index SNPs, overlapping groups of 40 and 207 index SNPs altered ASM-correlated or enriched binding motifs, respectively, providing mechanistic leads to disease-associated transcriptional pathways (**Table S11**). Interesting ASM index SNPs for this disease category include rs398206 associated with cutaneous melanoma and nevus counts via strong LD with GWAS SNPs rs416981 and rs45430 (nearest genes *FAM3B*, *MX2*, *MX1*), rs4487645 precisely coinciding with a GWAS peak SNP for multiple myeloma and immunoglobulin light chain amyloidosis (nearest genes *SP4*, *DNAH11*, *CDCA7L*; **Fig. 7**), rs3806624 precisely coinciding with a GWAS peak SNP for B cell lymphomas and multiple myeloma (nearest gene *EOMES*; **Fig. S23**), rs2853677 linked to lung cancer and other malignancies, as well as benign prostatic hyperplasia, via strong LD with several GWAS peak SNPs (genes *SLC6A18*, *TERT*, *MIR4457*, *CLPTM1L*; **Fig. 7**), and rs61837215 linked to breast cancer via LD with GWAS peak SNP rs2754412 (nearest genes *HSD17B7P2*, *SEPT7P9*, *LINC00999*; **Fig. S23**). Potentially informative examples in lenient blocks include rs2427290 linked to colorectal cancer via GWAS peak SNP rs4925386 (nearest genes *OSBPL2*, *ADRM1*, *MIR4758*, *LAMA5*, *RPS21*, *CABLES2*; **Fig. S8**), and rs2283639 linked to non-small cell lung cancer via GWAS peak SNP rs1209950 (nearest genes *LINC00114*, *ETS2*, *LOC101928398*; **Fig. S9**). Each of these index SNPs disrupts one or more strongly ASM-enriched and/or correlated TF binding motifs (**Table S11**).

#### ASM index SNPs in LD with GWAS peaks for neuropsychiatric traits and disorders and neurodegenerative diseases

We found 210 ASM DMRs containing 225 index SNPs in strong LD (R^2^>.8) with GWAS peak SNPs for neurodegenerative, neuropsychiatric, or behavioral phenotypes (**Table S12**). Among these ASM DMRs, about 15% showed ASM in brain cells and tissues (cerebral grey matter, neurons, glia), and a larger percentage showed ASM in immune system cell types. Both can be disease relevant, since studies have linked brain disorders not only to neuronal and glial cell processes but also to the immune system [46]. In addition, many loci in this list showed ASM in GBMs, which have partial glial and neuronal differentiation and may be revealing genetic variants that can affect early neuronal proliferation and differentiation. In these DMRs, 52 of the ASM index SNPs precisely coincided with the GWAS peak SNPs, supporting a functional regulatory role for these genetic variants, and overlapping groups of 36 and 183 index SNPs altered ASM-correlated or enriched binding motifs, respectively, providing mechanistic leads to disease-associated transcriptional pathways. Some interesting examples in strong LD with GWAS peaks for this general disease category (**Table S12**) include rs1150668 linked to risk tolerance/smoking behavior and wellbeing spectrum via GWAS peak SNPs rs1150668 (coinciding with the ASM index SNP) and rs62620225 (nearest genes *ZSCAN16*, *ZKSCAN8*, *ZNF192P1*, *TOB2P1*, *ZSCAN9*; **Fig. 7**), rs2710323 that coincides with a GWAS peak SNP for schizoaffective disorder, anxiety behavior, bipolar disorder and others (nearest genes *NEK4*, *ITIH1*, *ITIH3*, *ITIH4*, *MUSTN1*, *MIR8064*, *TMEM110*; **Fig. S22**), rs4976977 linked to intelligence measurement, anxiety measurement, schizophrenia, and unipolar depression via strong LD with GWAS peak SNP rs4976976 (nearest genes *MIR4472-1*, *LINC00051*, *TSNARE1*), rs667897 linked to Alzheimer’s disease via GWAS peak SNP rs610932 (nearest genes *MS4A2*, *MS4A6A*) and rs13294100, which coincides with a GWAS peak SNP for Parkinson’s disease (nearest gene *SH3GL2*). Each of these index SNPs disrupts one or more strongly ASM-enriched and/or correlated TF binding motifs (**Table S12**).

### Visualization of the ASM mapping data as annotated genome browser tracks

In addition to the three major disease categories detailed above, we found several hundred ASM index SNPs in strong LD with GWAS peaks for pharmacogenetic phenotypes or for cardiometabolic diseases and traits (e.g. rs2664280 linked to Type 2 diabetes mellitus via GWAS SNP rs2633310, **Fig. 7**). The final set of high-confidence recurrent ASM loci averaged 5 ASM DMRs per Mb of DNA genome wide. We provide the data both in tabular format (**Table S2**) and as annotated genome browser tracks that include the most useful parameters for each ASM index SNP. These parameters include ASM confidence and strength ranks, cell and tissue types with ASM, cancer vs normal status of the samples with ASM, and presence or absence of enriched CTCF or TF binding motifs and/or motifs with significant correlations of ASM strength with allele-specific differences in predicted binding affinity scores. An example of a 500 kb region of chromosome 19 containing 5 ASM DMRs, with ranks ranging from strong to weak and the strongest one encompassing a CTCF-bound insulator element, is in **Fig. S24**. These tracks (see Availability of Data) can be displayed, together with other relevant tracks, including chromatin structure for mechanistic studies and the GWAS catalogue track for potential disease associations, in UCSC Genome Browser sessions [47].

## Discussion

These data from dense mapping of ASM in normal human cell types and tissues, plus a group of cancers, identify 17,935 index SNPs in 15,115 DMRs that show strong and recurrent non-imprinted ASM, of which a substantial subset map within haplotype blocks that contain GWAS peaks for common diseases and related traits. In this study we focused on finding strong and high-confidence ASM DMRs, each containing multiple CpGs passing ASM criteria, and each detected in at least two independent samples. Thus, we sought to maximize true-positive findings, which were borne out by a high validation rate using targeted bis-seq. In addition to the value of these data for disease-focused post-GWAS studies, this high yield of stringently defined ASM DMRs, and inclusion of both cancer and normal cell types and tissues, allowed us to test mechanistic hypotheses for the creation of allele-specific CpG methylation patterns in ways that have not been feasible with prior datasets.

A recent study by Onuchic et al. using Human Epigenome Project (HEP) data provided a map of ASM SNPs based on 49 WGBS from 11 donors (non-cancer tissues) and 2 cell lines [11]. Using their publicly accessible processed data, we identified a set of strong ASM SNPs that pass similar effect size and p-value criteria as in our analysis (>20% methylation difference and corrected p-value<0.05). However, for harvesting these candidate ASM loci from their dataset, we did not require multiple CpGs in each sequence contig to show ASM, since although in our criteria this is a requirement, it was not utilized as a criterion by Onuchic et al. Overall, 50% of our informative SNPs were also informative in the HEP dataset and 31% of our ASM index SNPs passed the above criteria for ASM in the HEP WGBS data. Given the differences in analytical methods, and the differences in numbers and tissue types of the individuals analyzed, this is an encouraging convergence of findings. At the same time, this comparison indicates that our dataset adds substantial new information. With even greater numbers of individuals (informative heterozygotes at more SNPs), additional cell and tissue types, and greater depth of WGBS, additional loci with ASM will be identified. Our data already reveal a large component of rare or “private” ASM. Indeed, some of the ASM loci identified and validated by targeted bis-seq in our previous smaller study [8] are not included in our current list of recurrent ASM DMRs because they passed ASM criteria in only one individual. Conversely, as expected based on the requirement for multiple individuals when using a methylation QTL (mQTL) approach to detect ASM, the current ASM dataset now encompasses a larger percentage of the set of mQTLs identified in that prior study.

Allele specific binding of TFs and CTCF has been detected at up to 5% of assessed genomic sites [39], and the data provided here bolster and refine previous results from us and others [8, 9, 11, 28, 30, 31, 48] implicating a major role for binding site occupancies in shaping both net and allele-specific DNA methylation patterns in human cells. The harvest of large numbers of strong and high-confidence ASM occurrences in this study facilitated our analysis of individual (not pooled) binding motifs, thereby producing a statistically robust list of specific ASM-correlated CTCF and TF binding motifs, nearly all of which show anti-correlated (i.e. inversely correlated) behavior in which greater predicted binding site affinity and site occupancy tracks with less methylation of CpGs on that allele - which can be heuristically understood as protection of the occupied binding site from methylation.

The set of CTCF and TF binding motifs that we find to be strongly correlated with ASM when they contain disruptive SNPs overlaps only partly with the ASM-correlated motifs identified in the HEP study [11]. Encouragingly, certain classes of motifs emerge as significantly correlated in both studies. However, in addition to some differences in the identities of the most strongly correlated and enriched motifs, a general difference between the conclusions of the two studies concerns the numbers of motifs showing positive vs negative directions of the correlations. The HEP investigators reported a substantial minority subset (approximately 30%) of motifs for which higher predicted binding affinity was found to correlate with greater CpG methylation (i.e. positively or directly correlated behavior). In our dataset, using our ASM criteria and analytical pipeline, we find a nearly complete absence of such occurrences. All but 1 of the 144 motifs that are both enriched and significantly ASM-correlated (**Table S9**) show an inversely correlated direction of the relationship, such that higher predicted binding affinity (greater predicted binding site occupancy) tracks with relative CpG hypomethylation. When we only require ASM correlation, without enrichment as a criterion (**Table S8**), we find 175 motifs with this inversely correlated behavior, but only two motifs with positively correlated behavior in which greater predicted binding site occupancy tracks with CpG hypermethylation. Our combined ASM and ASB analysis, using ENCODE ChIP-seq data in the GM12878 LCL, also showed a strong enrichment of inverse correlations between TF binding and allelic CpG methylation levels. Interestingly however, in our small set of two positively correlated motifs we find the YY1 binding motif, which was also found by the HEP investigators in their positively correlated subset. This finding makes biological sense since the YY1 TF, acting as a component of the PRC2 polycomb repressive complex, can attract CpG methylation, at least partly through recruitment of DNA methyltransferases [49].

An advance in the current study is our ability to test and compare mechanisms of ASM in normal and neoplastic cells. We observed a dramatic increase in per sample ASM frequencies, on average, in the primary cancers compared to cell lineage-matched normal cells and to non-cancer samples overall. This increase was paralleled by a more modest but still significant increase in ASM frequency in whole placental tissue and in purified trophoblast, which, as shown here and in other studies [8, 21, 23], have global CpG hypomethylation similar to cancers. Special aspects of ASM detected in the cancers included allele-specific hypomethylation genome-wide and allele-specific hypermethylation at loci in poised chromatin, as well as relatively increased ASM in chromatin desert regions and increased allele-switching at ASM loci. Despite these differences, our findings from testing for enrichment of TF and CTCF binding motifs and correlations of ASM with disruptive SNPs in these motifs clearly indicate that the same binding site occupancy mechanism pertains in both normal and cancer-associated ASM. A striking additional result that supports this shared mechanism, and which may have important implications for cancer biology, is our finding of de novo ASM affecting CpGs clustered around somatic point mutations in cancer cells. The key mechanistically informative feature of this de novo ASM is that it preferentially occurs around mutations that disrupt or create the same classes of TF binding motifs that are linked to ASM in normal cells. While this topic will need future work, we can speculate that some of these non-coding mutations, which declare their functionality by producing the observed de novo ASM, might play roles in cancer biology through effects on gene expression. A possible example is the *TEAD1* gene, which is known to be over-expressed in aggressive and treatment-refractory cases of multiple myeloma [50] and which showed somatic mutation-associated de novo ASM in its promoter/enhancer region in a multiple myeloma case in our series (**Fig. 6**).

Based on the shared general mechanism of ASM in cancer and normal cells, an important practical conclusion is that analyzing combined series of cancer cases plus non-cancer samples increases the power of ASM mapping for finding mechanistically informative rSNPs. In conjunction with GWAS data these rSNPs can point to genetically regulated transcriptional pathways that underlie inter-individual differences in susceptibility not only to cancers but also to nearly all common human non-neoplastic diseases. Due to the LD structure of the genome, GWAS peaks by themselves can only point to disease-associated haplotype blocks, with all SNPs in strong LD with the causal SNP(s) showing similar correlations to the phenotype. Therefore, additional types of evidence are needed before causal roles can be attributed to GWAS peak SNPs or to other SNPs in strong LD with them. ASM mapping can pinpoint candidate rSNPs that declare their presence by conferring the observed physical asymmetry in CpG methylation between the two alleles. The key finding that supports such mapping for biologically meaningful rSNP discovery is the one above, namely that ASM is caused by disruptive SNPs in TF and CTCF binding sites.

This informative situation is highlighted by our findings for ASM index SNP rs4487645 (**Fig. 7**), which coincides with a GWAS peak for AL amyloidosis and multiple myeloma and disrupts an ENCODE PAX5 discovery motif (PAX5_disc3) that is significantly enriched among ASM loci. Since the PAX5 TF is a master regulator of B cell development [51], these ASM mapping data are post-GWAS evidence suggesting involvement of a relevant biological pathway in susceptibility to multiple myeloma, a B cell malignancy. That the ASM at this locus was specifically found in a sample of DLBCL (another type of B cell cancer) highlights the usefulness of including primary tumor samples in ASM mapping. Another example is the ASM index SNP rs2283639, is linked to lung cancer GWAS peak SNP rs1209950. This ASM index SNP is situated in the promoter/enhancer region of the *ETS2* gene, where it disrupts an ASM-enriched ETS1_3 TF binding motif (**Fig. S9**). A promising example in a non-neoplastic disease, is provided by ASM index SNP rs2664280, which disrupts multiple ASM-enriched and ASM-correlated JUNB and AP1 binding motifs (all with greater predicted binding affinity on the REF allele) and is in strong LD with a GWAS peak SNP for psoriasis (**Fig. 7**). For this example, the ASM was found in T cells, which are relevant for psoriasis, and the candidacy of the JUNB motif disruption as a biological explanation for the disease association is supported by other evidence for involvement of AP1-dependent transcriptional changes in this disease [52]. These situations can be tested further by functional experiments such as CRISPR/Cas9-mediated DNA deletions in ASM DMRs and mutations of ASM index SNPs in appropriate cell types.

Lastly, regarding the non-redundancy of ASM mapping as a post-GWAS approach, while SNPs with experimental evidence for ASB are strongly enriched among the ASM loci reported here, more than 90% of the ASM index SNPs harvested in this study lack currently available ASB annotations. Thus, maps of ASM, which are readily generated from large archival collections of DNA samples, can provide information about rSNPs that has not emerged from other types of mapping data, such as ChIP-seq for ASB, which require whole cells or tissue samples and are more technically difficult to obtain. That ASM data are largely non-redundant with other post-GWAS modalities (ASB, chromatin states and chromatin accessibility, eQTLs) is further highlighted by our observation of ASM DMRs in chromatin deserts. Our finding of similar correlations of ASM with disruptive SNPs in specific TF binding motifs in both non-desert and desert regions suggests that mapping ASM in deserts can pinpoint candidate rSNPs in cryptic TF binding sites, which were presumably active at earlier stages of cell differentiation and have left “methylation footprints” that can be detected as ASM but cannot be found using other mapping methods.

## Conclusions

We mapped ASM genome-wide in DNA samples including diverse normal tissues and cell types from multiple individuals, plus three types of cancers. The data reveal 15,115 high-confidence ASM regions, of which 1,842 contain SNPs in strong LD or precisely coinciding with GWAS peaks for human diseases and traits. We find that ASM is increased in cancers, due to widespread allele-specific hypomethylation and focal allele-specific hypermethylation in regions of poised chromatin, with cancer-associated epigenetic variability manifesting as increased allele switching. We also report rare but informative de novo ASM due to somatic mutations in TF binding sites in cancers. Despite these cancer-specific phenomena, enrichment and correlation analyses indicate that disruptive SNPs in specific classes of CTCF and TF binding motifs are a shared mechanism of ASM in normal and cancer cells, and that this mechanism also underlies ASM in “chromatin deserts”, where other post-GWAS mapping methods have not been informative. We provide our dense ASM maps as genome browser tracks and show examples of ASM index SNPs that are in LD with GWAS peaks and disrupt TF binding motifs, thereby nominating specific transcriptional pathways in susceptibility to autoimmune diseases, neuropsychiatric disorders, and cancers.

## Materials and methods

### Human cells and tissues

Human tissues and cell types analyzed in this study are listed in **Table S1**. The Agilent SureSelect series included 9 brain (cerebral cortex), 6 T cell (CD3+), 3 whole peripheral blood leukocyte (PBL), 2 adult liver, 1 term placenta, 2 fetal heart, 1 fetal lung, and one ENCODE lymphoblastoid cell line (LCL; GM12878). All samples were from different individuals, except for a trio among the brain samples consisting of one frontal cortex (Brodmann area BA9) and two temporal cortex samples (BA37 and BA38) from the same autopsy brain. We performed WGBS on 16 normal T cell preparations (10 CD3+, 4 CD4+, and 2 CD8+), 10 B cell samples (CD19+), 7 monocyte (CD14+) and 2 monocyte-derived macrophage samples, 2 PBL, 1 reactive lymph node, 4 fractionated samples from a term placenta (whole tissue from the chorionic plate, purified villous cytotrophoblast from chorionic plate and basal plate, and extravillous trophoblast from basal plate), 3 adult liver, 2 primary bladder epithelial cell cultures, 2 epithelium-rich non-cancer tissue samples from breast biopsies, 3 primary mammary epithelial cell cultures, 3 frontal cerebral cortex grey matter samples, 6 NeuN+ FANS-purified cerebral cortex neuron preparations, 4 NeuN-FANS-purified cerebral cortex glial cell preparations, 1 LCL (GM12878), 3 B cell lymphomas (1 follicular and 2 diffuse large B cell type), 7 multiple myeloma cases (CD138+ cells from bone marrow aspirates), and 6 cases of glioblastoma multiforme (GBM). The glia samples were paired with neuron preparations from the same autopsy brains, and several of the B cell, PBL, monocyte/macrophage, and T cell samples were from the same individuals (**Table S1**). In the combined series, 5 samples were assessed by both SureSelect and WGBS (**Table S1**). Peripheral blood samples were obtained with informed consent, and CD3+ T-lymphocytes, CD19+ B-lymphocytes and CD14+ monocytes were isolated by negative selection using RosetteSep kits (Sigma). Macrophages were produced from monocytes by culturing in RPMI with 20% fetal calf serum with 50 ng/ml M-CSF for one week as described [53]. Fractionation of villous cytotrophoblast and extra-villous trophoblast from a term placenta was carried out as previously described [54]. All other non-neoplastic primary human tissues were obtained from autopsies. Neuronal and glial cell nuclei were prepared from autopsy brains using tissue homogenization, sucrose gradient centrifugation and fluorescence activated nuclear sorting (FANS) with a monoclonal anti-NeuN antibody [55] and documented for purity of cell types by immunostaining of cytospin slides, as shown previously [56]. Biopsy samples of human lymphomas and GBMs, and CD138+ multiple myeloma cells isolated from bone marrow biopsies by positive selection on antibody-conjugated magnetic beads (Miltenyi Biotec), were obtained with I.R.B. approval in a de-identified manner from the Tissue Biorepository of the John Theurer Cancer Center. Absence of circulating myeloma cells in the paired B cell samples was verified by cytopathology and by the absence of DNA copy number aberrations that were seen in the multiple myeloma cells. Among the 6 GBMs, we did not detect cases with a strong CpG island hypermethylator phenotype (CIMP) as defined by Noushmehr et al [57], which is expected given that CIMP is more frequent in low-grade gliomas than in high-grade GBMs. In surgical specimens, GBM cells are mixed with non-neoplastic glial and vascular cells, but the presence of malignant cells in each GBM sample was confirmed by histopathology on sections of the tissue blocks and was verified by assessing DNA copy number using normalized WGBS read counts [56], which revealed characteristic GBM-associated chromosomal gains and losses. The GM12878 lymphoblastoid cell line was purchased from Coriell, primary cultures of non-neoplastic human urinary bladder epithelial cells were purchased from A.T.C.C., and primary cultures of non-neoplastic human mammary epithelial cells were purchased from Cell Applications, Inc. and ScienCell Research Laboratories.

### Agilent SureSelect Methyl-seq and WGBS

We used the Agilent SureSelect methyl-seq DNA hybrid capture kit according to the manufacturer’s protocol to analyze methylomes in a total of 27 non-neoplastic cell and tissue samples (**Table S1**). In this protocol, targeted regions (total of 3.7M CpGs) including RefGenes, promoter regions, CpG islands, CpG island shores, shelves, and DNAse I hypersensitive sites are sequenced to high depth. DNA was sheared to an average size of 200 bp and bisulfite converted with the EZ DNA methylation kit (Zymo). Paired end reads (100, 150 or 250 bp) were generated at the Genomics Shared Resource of the Herbert Irving Comprehensive Cancer Center of Columbia University, with an Illumina HiSeq2500 sequencer.

For analyzing complete methylomes in the normal and tumor samples, plus the GM12878 LCL, WGBS was performed at the New York Genome Center (NYGC), MNG Genetics (MNG) and the Genomics Shared Resource of the Roswell Park Cancer Institute (RPCI), as indicated in **Table S1**. The NYGC used a modified Nextera transposase-based library approach. Briefly, genomic DNA was first tagmented using Nextera XT transposome and end repair was performed using 5mC. After bisulfite conversion, Illumina adapters and custom bisulfite converted adapters are attached by limited cycle PCR. Two separate libraries were prepared and pooled for each sample to limit the duplication rate and sequenced using Illumina X system (150 bp paired-end). WGBS performed at MNG used the Illumina TruSeq DNA Methylation Kit for library construction according to the manufacturer’s instructions and generated 150 bp paired end reads on an Illumina NovaSeq machine. WGBS performed at RPCI utilized the ACCEL-NGS Methyl-Seq DNA Library kit for library construction (Swift Biosciences) and generated 150 bp paired end reads on an Illumina NovaSeq.

### Read mapping, SNP calling, and identification of ASM DMRs

Our analytical pipeline is diagrammed in **Figure S1**. Compared with our previous study [8], updates included improvements in sequence processing, updated database utilization and increased stringency for SNP quality control, assignment of both strength and confidence scores to ASM index SNPs, use of updated ENCODE and JASPAR databases [36, 58] for scoring the effects of the ASM index SNPs on predicted TF binding affinities, and utilization of haplotype blocks and LD criteria, instead of simple distance criteria around GWAS peaks for nominating disease-associated rSNPs in ASM DMRs. After trimming for low-quality bases (Phred score<30) and reads with a length <40 bp with TrimGalore, the reads were aligned to the human genome (GRCh37) using Bismark [59] with paired end mode and default setting allowing about 3 mismatches in a 150 bp read. For the SureSelect methyl-seq samples, unpaired reads after trimming were aligned separately using single end-mode and the same settings. Duplicate reads were removed using Picard tools [60] and reads with more than 10% unconverted CHG or CHH cytosines (interquartile range: 0.1-2.2% of mapped reads; median 0.14%) were filtered out. Depth of sequencing for each sample in **Table S1**, with metrics calculated using Picard tools. SNP calling was performed with BisSNP [61] using default settings, except for the maximum coverage filter set at 200 to encompass deep sequencing while avoiding highly repetitive sequences, and quality score recalibration. SNP calling was carried out using human genome GRCh37 and dbSNP147 as references. For ASM calling, only heterozygous SNPs are informative. We filtered out heterozygous SNPs with less than 5 reads per allele. In addition, SNP with multiple mapping positions were filtered out, as well as SNPs with more than one minor allele with allele frequency>0.05. Informative SNPs were defined as heterozygous, bi-allelic and uniquely mapped SNPs that did not deviate significantly from Hardy-Weinberg equilibrium based on exact tests (FDR<0.05 by HardyWeinberg R package) and were covered by more than 5 reads per allele. In addition, we filtered out any informative regions mapping ENCODE defined “blacklisted” regions [62]. Informative regions were defined as regions with overlapping reads covering at least one informative SNP. Bisulfite sequencing converts unmethylated C residues to T, while methylated C residues are not converted. Therefore, for C/T and G/A SNPs the distinction between the alternate allele and bisulfite conversion is possible only on the non-C/T strand. For SureSelect methyl-seq, since only negative stranded DNA fragments are captured, G/A SNPs were filtered out; for WGBS, C/T and G/A SNPs were assessed after filtering out reads mapping to the C/T strand.

ASM calling was performed after separating the SNP-containing reads by allele. For each heterozygous SNP, all reads overlapping the 2 kb window centered on the SNP were extracted using Samtools. Given the median insert size of our libraries (∼200 bp), the use of a 2 kb window instead of the SNP coordinate allows extraction, in most cases, of both paired ends even if the SNP is only covered at one of the ends. SNP calling is performed on each paired read and read IDs are separated into two files as reference (REF) and alternate (ALT) alleles using R. After Bismark methylation extractor is applied, CpG methylation calls by allele are retrieved using allele tagged read IDs. Paired reads with ambiguous SNP calling (i.e., called as REF allele on one paired end and ALT allele on the other) were discarded. For Nextera WGBS, due to the fill-in reaction using 5mC following DNA tagmentation which affects the 10 first base pairs (bp) on 5’ of read 2, methylation calling for Cs mapping to these bp were not considered. In addition, a slight methylation bias due to random priming and specific to each library kit was observed within the last 2 bp on 3’ of both paired ends for Nextera WGBS, within the first 10 bp on 5’ of both paired ends and the last 2 bp on 3’ of read 2 for TruSeq WGBS, and within the first 10 bp on 5’ of read 2 for ACCEL-NGS WGBS. Therefore, methylation calls in these windows were ignored.

To further increase the stringency and accuracy of ASM calling, only regions with at least 3 CpGs covered by more than 5 reads per allele were considered. ASM CpGs were then defined as CpGs with Fisher’s exact test p-value <0.05 and ASM DMRs were defined as regions with ≥20% methylation difference after averaging all CpGs covered between the first and last CpGs showing ASM in the region, a Wilcoxon p-value corrected for multiple testing by the B-H method <0.05 (FDR at 5%) and more than 3 ASM CpGs including at least 2 consecutive ASM CpGs. CpGs destroyed by common SNPs (maf>0.05) were filtered out from both CpG and DMR level analyses. Very close or overlapping DMRs (<250 intervening bp) were merged into one unique DMR.

We ranked the ASM SNPs using two approaches, one based on confidence/recurrence criteria and the other on percent difference in methylation of the two alleles (ASM strength). For the confidence rank, we used the geometric mean of the average coverage of each allele, the number of samples showing ASM, and the percentage of these samples among all heterozygous (informative) samples. For the strength rank, we used the geometric mean of the methylation difference, number of ASM CpGs and percentage of ASM CpGs among all covered CpGs. An overall rank was calculated using the geometric mean of these two ranks. ASM DMRs dictated by multiple index SNPs were ranked by the top-scoring SNP. ASM calling and ranking were performed using R and Stata 15. We used the GeneImprint database to flag and exclude from downstream analyses all ASM DMRs that mapped within 150Kb windows centered on the transcription starting site of all known high confidence imprinted genes, including in this list the *VTRNA2-1* gene, which we have previously shown to be subject to parental imprinting in trio samples [35] and which showed frequent allele switching in normal samples in the current dataset, consistent with imprinting (**Table S4**).

Lastly, although varying levels of non-CpG methylation (mCH) have been observed in human and mouse tissues, and this non-canonical methylation appears to have unique sub-chromosomal distributions and biological functions [63], for clarity the current report is focused only on ASM affecting classical CpG methylation. Nonetheless, giving confidence in our dataset, we found mCH to be higher in the purified cerebral cortical neurons, (2.4% +/- 0.9%, N=16) than in the non-neuronal samples (0.47% +/- 0.54%, N=43), which is consistent with findings from another laboratory [64, 65].

### Detection of somatic mutations

Somatic mutation calling was performed on the 4 multiple myeloma samples for which paired normal peripheral blood B cells from the same individuals had been bis-sequenced using the same library preparation (ACCEL-NGS WGBS). We used BisSNP (with the same setting as for SNPs but without providing reference SNP dataset) to call all heterozygous variants for both myeloma and normal B cells samples. We then filtered out any variants reported as germline SNPs by DbSNP147. Variants mapping to ENCODE blacklisted regions were removed, and we next filtered out any variants that were present in the paired B cell samples. We used a sequencing coverage requirement for candidate mutations in the myeloma cases of 10X per allele (wild-type, mutant).

### Targeted bisulfite sequencing (bis-seq) for validations of ASM

Targeted bis-seq was utilized for validation of ASM regions. PCR primers were designed in MethPrimer [60], and PCR products from bisulfite-converted DNA samples were generated on a Fluidigm AccessArray system as described previously [8], followed by sequencing on an Illumina MiSeq. PCRs were performed in triplicate and pooled to ensure sequence complexity. ASM was assessed when the depth of coverage was at least 100 reads per allele. While the absolute differences between methylation of the two alleles are not exaggerated by deep sequencing, the p-values for these differences tend to zero as the number of reads increases. Therefore, to avoid artificially low p-values, we carried out bootstrapping (1000 random samplings, 50 reads per allele), followed by Wilcoxon tests for significance. Samplings and bootstrapping were performed using R. The tested ASM loci and amplicon coordinates are in **Table S6**.

### Annotation and enrichment analysis of ASM loci

To annotate ASM and informative SNPs, we defined small (1000 bp) and large (150 kb) windows centered on each index SNP. The small windows were used to assess mechanistic hypotheses involving local sequence elements and chromatin states and the large windows were used for functional annotation (genes and GWAS associated SNPs). We used BedTools to intersect the genomic coordinates of ASM windows to the coordinates of the annotation sets. From the UCSC Genome Browser (GRCh37 assembly) we downloaded RefSeq annotations, DNase hypersensitive sites, TF peaks by ChIP-seq, and chromatin state segmentation by HMM in ENCODE cell lines[66]. We allowed multiple chromatin states at a single location when different states were present in different cell lines. Distances between ASM loci and genes were calculated from the transcription start sites. Regulome scores were downloaded from RegulomeDB [42]. For each relevant feature, enrichment among ASM index SNPs compared to the genome-wide set of nformative SNPs (SNPs that were adequately covered and heterozygous in at least 2 samples) was tested using bivariate logistic regressions. To compare characteristics of ASM observed only in cancer samples (“cancer-only ASM”) vs ASM observed in at least one non-cancer sample (“normal ASM”), these analyses were stratified by cancer status. To assess enrichment for chromatin states among ASM loci that were found only in cancers or only in non-cancer samples, with the occurrences divided into subsets according to the direction of the change in methylation in the cancers compared to cell lineage-matched normal samples, we used the same approach but considering only the sets of heterozygous SNPs informative in both myelomas and B cells, or lymphomas and B cells, or GBMs and glia. To compare the regulatory features of ASM to those of other allele-specific marks, we performed similar analyses for enrichment of ASM index SNPs in the sets of publicly available eQTLs [58] and ASB SNPs [24, 39] that were informative in our dataset.

### Tests for correlations of ASM with SNPs in TF and CTCF binding sites

To test for correlations of ASM with disruptive SNPs in TF binding motifs, we used position weight matrices (PWMs) of TF motifs from ENCODE ChIP-seq data [33, 36], as well as PWMs from the JASPAR database [37, 58]. We scored allele specific binding affinity at each index SNP using the atSNP R package [40], which computes the B-H corrected p-values (i.e. q-values) of the affinity scores for each allele and q-value of the affinity score differences between alleles. Motifs that contained SNPs affecting allele specific TF binding affinity were defined as motifs with a significant difference in binding affinity scores of the two alleles (q-value<0.05) and a significant binding affinity score in at least one allele (p-value<0.005). For each TF occurrence, the binding scores per allele were estimated using PWM scores calculated as described in our earlier study [8]. In addition, among the ASM index SNPs, we specifically annotated TF binding motifs that overlapped with cognate TF ChIP-seq peaks based on ENCODE data[66]. For each motif, we used data from Kheradpour and Kellis [33, 36] to define the cognate TF peaks, required a 10-fold enrichment of the motif among ASM loci compared to background, and filtered out TF peaks with less than 10 occurrences of the tested motif among ASM loci.

To test whether ASM index SNPs are enriched in variants that disrupt polymorphic TF binding motifs, we used logistic regressions to calculate O.R.s for each disrupted polymorphic motif. Enrichment was defined as an O.R.>1.5 and B-H corrected p-value <0.05. Since computing resources required to run atSNP for >2 million SNPs and > 2000 TF motifs are extremely large, we random sampled 40,000 non-ASM informative SNPs (1:3 ASM vs non-ASM SNP ratio) to estimate the random expectation of each TF motif occurrence. To test whether the disruption of TF binding sites could be a mechanism of ASM, we correlated the difference in PWM scores between alleles of each occurrence of a given TF motif disrupted by an ASM index SNP to the differences in methylation levels between the two alleles, using linear regression. Only TF motifs with more than 10 disrupted occurrences in ASM regions were analyzed. For index SNPs showing ASM in multiple samples, we used the average methylation difference between the two alleles. For each TF motif, a significant correlation of ASM with predicted TF binding affinity differences between the two alleles was defined as FDR<0.05 and R^2^ >0.4.

To ask whether the correlations between ASM and predicted TF binding affinity differences between alleles might be similar for ASM loci found only in cancers compared to ASM loci that were observed in at least one normal sample, and to ask this same question for chromatin desert ASM vs non-desert ASM regions, we used a multivariate mixed model with random slope and intercept, with pooling of TF motifs at this step to reach sufficient power (number of occurrences used for the regression). TF motifs with less than 10 occurrences total, or less than 3 occurrences in any ASM class, were filtered out. TF motifs included in the final mixed models for the four classes of ASM loci were pre-selected from the bivariate model (performed without distinction of ASM class; requiring FDR<0.05 and R^2^ >0.4). To not bias the analysis toward TF motifs without any ASM class effect (which might be overrepresented in the set of significant TF motifs identified in the bivariate analyses), we also screened each TF motif, including CTCF motifs, using separated multivariate linear fixed models to include any motifs showing no correlation overall but a correlation trend only in one of the ASM classes (FDR<0.05 for at least one of the ASM classes, multivariate model adjusted R^2^>0.4).

We defined chromatin deserts as 1 kb genomic windows, centered on ASM index SNPs, which contained no DNAse peaks or only one DNAse peak among the 122 ENCODE cell lines and tissues, and no strong active promoter/enhancer, poised, or insulator chromatin state in any ENCODE sample. The multivariate mixed model accounts for both intra- and inter-TF motif error terms and includes the predicted TF binding affinity difference, either for two classes of ASM loci (non-cancer ASM and cancer ASM) or 4 classes of ASM loci (non-cancer ASM in non-desert regions, non-cancer ASM in desert regions, cancer ASM in non-desert regions, and cancer ASM in desert regions), the interaction between ASM class and binding affinity as fixed explanatory covariates for the methylation difference, and the TF motif as a random covariate. Marginal effects from predictions of the mixed model and Bonferroni-corrected p-values were then computed to compare the correlation between ASM classes. The variation due to the TF motif was considered as a random effect, under the assumption that each TF motif might have a different intercept and slope. The interaction terms reflect the difference in the methylation to binding affinity correlation between each ASM class compared to the reference class, which we defined as non-cancer ASM for the 2-calss analysis and non-cancer ASM in non-desert region for the 4-class analysis. Analysis after excluding ASM loci that showed switching behavior gave similar results. TF motifs with significant correlations of disruptive SNPs with ASM for at least one of the 2 or 4 ASM classes (FDR <0.05 and R^2^ >0.4) were then pooled to be tested in the final mixed model, such that the model was run using a total of 178 TF motifs with 16,609 motif occurrences disrupted by 3,394 ASM SNPs for the 2 ASM-class analysis and a total of 62 TF motifs with 10,709 motif occurrences disrupted by 1,967 ASM SNPs for the 4 ASM-class analysis. To assess ASB in the GM12878 cell line, ChiP-seq data for 154 TFs available for this cell line were downloaded from ENCODE. For each TF, SNP genotyping and allele specific read count were performed using the ChiP-seq alignment data for the set of high confidence ASM SNPs found in our GM12878 data and compared to data from WGBS. ASB SNPs were defined as SNPs showing homozygous genotype in the ChiP-seq data (but heterozygous in WGBS) with a significant allele-specific occupancy bias (FDR <0.05, Fisher exact test). All analyses were performed using R and STATA statistical software.

### Associations of ASM with GWAS peaks

GWAS traits and associated supra and subthreshold SNPs (p<10^-6^) were downloaded from the NHGRI GWAS catalog [42, 67]. We defined haplotype blocks using 1000 Genomes phase 3 data [67] based on the method of Gabriel et al. for scoring linkage disequilibrium (LD) with emphasis on D-prime values [45] in PLINK [68]. To identify GWAS peaks in moderate LD with ASM index SNPs, we used relaxed criteria of D-prime confidence intervals (0.60-0.84) and historical recombination (0.55) but set the maximum haplotype block size at 200 kb to minimize large block calling in genomic regions lacking haplotype block structure. The blocks so defined have a median size of 46 kb. To identify ASM SNPs in strong LD with GWAS peak SNPs, we utilized the default parameters of Gabriel et al. for haplotype block calling [45]. The blocks so defined have a much smaller median size of 5 kb. Finally, we computed pairwise R^2^ between our ASM SNPs and all GWAS SNPs within 200kb. SNPs with high R^2^ represent a subtype of SNPs in high LD where not only a non-random association (high D’) is observed but where these SNPs can essentially be considered as proxies of each other. Statistical association between a GWAS SNP and trait can be directly imputed to any SNPs with very high R^2,^ so such SNPs are obvious candidates for post-GWAS analyses. However, SNPs showing high D’ but low R^2^ with the GWAS SNP (which occurs when a rare SNP is in high LD with a more frequent SNP) might also contribute biologically to disease associations. We annotated each ASM index SNP for localization within these haplotype blocks, and for precise co-localization with a GWAS peak SNP or high R^2^ (>0.8), and tested for enrichment of ASM SNPs within these blocks, as well as among GWAS peak SNPs, using the same approach as described above for other genomic features.

## Author Contributions

CD and BT designed the research. CD, ED, MS, AC, HM, LM, AJ, PLN and KC, generated the data. CD, ED and BT carried out data analysis. CD, ED and SM set up the cloud computing platform. AMC, SL, CT, KM, LH, CM, SG, GK, DS and BT obtained the research funding. CD, ED and BT wrote the manuscript, with suggestions from AC, HM, LM, AJ, SCD, NPI, GB, SL, AMC, SM, PHRG, RF, CT, KM, LH, CM, AG, KC, SG, GK and DS. All authors read and approved the final manuscript.

## Ethics approval and consent to participate

Peripheral blood samples were obtained with informed consent. Primary tumor samples were collected by the JTCC Tissue Biorepository and transferred to the laboratory in a de-identified manner under I.R.B.- approved protocols.

## Availability of Data

The Agilent SureSelect and WGBS data have been submitted to NCBI/GEO. Examples of custom genome browser tracks with annotated ASM loci can be viewed at a UCSC browser session hosted by our laboratory (https://bit.ly/tycko-asm).

## Supplemental Figure Legends

**Figure S1. Flow charts of computational and analytical approaches in this study.**

**A**, (Left panel) Steps for ASM calling and ranking, including ASM definition and criteria (See Methods). Our ASM definition incorporates both individual CpG and DMR-wide (multiple CpG) statistical criteria. (Right panel) Analytical pipeline for post-calling annotation and analyses to test ASM mechanisms, comparing ASM sub-classes (cancer vs non-cancer; desert vs non-desert), and overlaying that information with public GWAS data to nominate disease associated rSNPs and disrupted TF binding motifs. **B,** diagram of ASM criteria utilized in this study.

**Figure S2. Summary of sample types and numbers and yield of informative SNPs**

**A**, Summary of samples sequenced by Agilent SureSelect and WGBS. Additional information is in **Table S1**. Since our high confidence ASM set required ASM in at least 2 samples, the final informative SNP set used for downstream analyses corresponds to the 2,485,759 SNPs that were informative in at least two samples. The Venn diagrams are schematic, not drawn to scale. The percentages are calculated on the union of Agilent and WGBS SNPs. In other words, 1% of the total number of informative SNPs in our dataset were found only in SureSelect samples, 3% are found in both WGBS and SureSelect and 96% only in WGBS samples, which means that 4% (3 + 1) of the SNPs were informative in SureSelect and 99% (96 + 3) in WGBS. **B**, Map of a region of chromosome 20, showing an increased yield of ASM SNPs in WGBS compared to SureSelect, as expected based on genomic coverage and a greater number of samples and therefore informative heterozygotes, with consistent findings in regions covered by both methods.

**Figure S3. PCA of the combined WGBS and SureSelect methyl-seq data and series-wide overlap between ASM loci detected by the two methods**

**A**, PCA performed using net methylation values of CpGs on chromosome 20. Only CpGs informative (>10X) in both Agilent SureSelect and WGBS were used. The PCA shows clear clustering by cell/tissue and cancer type. Similar results were found using methylation data from other autosomes. **B**, Pie chart showing the proportion of high confidence ASM SNPs found in more than two biological samples, identified by WGBS and by SureSelect methyl-seq. Numbers of ASM SNPs are in parenthesis. **C**, Venn diagram showing a cross-platform comparison, with the percentage of high confidence ASM index SNPs that were identified series-wide in both assays. Only SNPs informative in both assays (adequate sequence coverage and heterozygous genotype calls) were considered for this comparison.

**Figure S4. Distribution of ASM shows a high proportion of rare or private ASM in both cancer and normal samples and a significant increase in per-sample ASM in the cancers.**

**A**, Most of the ASM calls are found only in one sample. While many might be genuine ASM associated with rare SNPs or with inter-individual variability, all downstream analyses in the current study are focused on recurrent ASM detected in at least two samples. **B** and **C**, Approximately one third of the ASM DMRs were identified only in cancer samples (referred to here as “cancer-only ASM”). Given that our study included more non-cancer than cancer samples, this high proportion of ASM SNPs found only in the cancers is significantly increased compared to random expectation.

**Figure S5. Comparison of results from individual DNA samples analyzed by two WGBS library construction kits at two sequencing facilities, or by SureSelect and WGBS.**

**A,** Cross platform and library correlations between net methylation showing a good R^2^ coefficient ranging from 0.94 to 0.96 for the 9 samples assessed by two WGBS library preparation protocols or two platforms (SureSelect, WGBS), and improve in regions with deeper coverage. **B**, Cross platform and cross-library correlations between allelic methylation differences, showing that most of the ASM found in only one facility or one platform shows the same methylation difference trend (i.e. “direction” of the allelic methylation bias) in both facilities, but is sub-threshold in the data from one facility, in terms of p-value and/or number of ASM CpGs.

**Figure S6. Example of a chromosome region illustrating consistency between SureSelect methyl-seq, WGBS, and targeted bisulfite sequencing**

**A**, Map of the region on chromosome 20 containing the ASM index SNP rs2427290. When covered in both SureSelect and WGBS, the net methylation is consistent between both assays, and shows low methylation “wells” at CpG islands, as expected. ASM dictated by index SNP rs2427290 is detected in both assays, with additional ASM SNPs found by WGBS, as expected. **B**, Primary sequencing data from WGBS, Agilent SureSelect methyl-seq, and targeted bis-seq, showing consistent findings of ASM in T cell samples. Rows represent sequence reads, and columns CpG sites in these reads. All samples are heterozygous, and the reads are separated by allele. Methylated CpGs are black circles, and unmethylated CpGs are white circles.

**Figure S7. Validations of ASM DMRs in disease-associated chromosomal regions: rs1041163 and multiple sclerosis**

**A**, Map showing the ASM region associated with rs1041163 in a putative tissue-specific promoter/enhancer region of the *CCDC155* gene, downstream of the *DKKL1* gene on chromosome 19. ENCODE chromatin state tracks suggest dynamic regulation, with active or quiescent marks depending on the cell type. Bisulfite PCR amplicons were designed to overlap the ASM and flanking SNPs, and to include at least 3 CpGs. The SNP is in high LD and R^2^ (R^2^ =0.98) with the rs2303759 GWAS peak SNP associated with multiple sclerosis, and another amplicon was designed to assess possible ASM at this position (which did not show ASM in our genome-wide data). The ASM index SNP disrupts a C-rich EGR1 TF binding motif and an EGR1 ChIP-seq peak found in K562 cells, supporting rs1041163 as a strong candidate rSNP. **B**, Targeted bis-seq reads, validating a discrete ASM regions (amplicon 2 and part of amplicon 3) spanning ∼700 bp in T cells and brain. Number of ASM samples, informative tissue types and additional annotations are in **Table S2**. The targeted bis-seq showed no evidence of ASM at the GWAS peak SNP, as expected. Rows represent sequence reads, and columns CpG sites in these reads. All samples are heterozygous, and the reads are separated by allele. Methylated CpGs are black circles, and unmethylated CpGs are white circles.

**Figure S8. Validation of ASM DMRs in disease-associated chromosomal regions: rs2427290 and colorectal cancer**

**A**, Map showing the ASM region tagged by SNP rs2427290 in the *LAMA5* gene on chromosome 20. The same region is shown, for a different purpose, in **Figure S5**. This region has an active promoter/enhancer state (color-coded red and yellow) in the GM12878 LCL (asterisk) but is in a Txn state (coded green) without promoter characteristics in other ENCODE cell lines. The ASM index SNP is close to (6.5 kb distance) and in moderate LD (lenient haplotype block; D’=0.8) with GWAS peak SNP rs4925386 associated with colorectal cancer. The ASM index SNP is in a region of open chromatin (DNAse hypersensitivity) and disrupts an ENCODE discovery motif for CCNT2 TF binding, located in a cluster of other TF binding sites and ENCODE ChIP-seq signals. The relatively hypomethylated allele is the one with the higher predicted CCNT2 binding affinity. For this occurrence, the motif maps the negative strand and is reported from 3’ to 5’ as per atSNP convention. **B**, Targeted bis-seq reads validating a discrete ASM regions (amplicon 2) spanning ∼400 bp in T cells and colonic mucosa. ASM is not found at the GWAS peak SNP. Numbers of ASM samples in each cell and tissue type, and additional annotations, are in **Table S2**. Rows represent sequence reads, and columns CpG sites in these reads. All samples are heterozygous, and the reads are separated by allele. Methylated CpGs are black circles, and unmethylated CpGs are white circles.

**Figure S9. Validation of ASM DMRs in disease-associated chromosomal regions: rs2283639 and non-small cell lung carcinoma**

**A**, Map showing the ASM region tagged by index SNP rs2283639, located in a promoter/enhancer region (color-code yellow and red) of the *ETS2* gene on chromosome 21. The SNP is very close to (4 kb distance) and in partial LD (lenient haplotype block; D’=.96) with GWAS peak SNP rs1209950, associated with survival after treatment of non-small cell lung carcinoma. The ASM index SNP disrupts an ENCODE-discovery motif for SMC3 (cohesion complex component), and it co-localizes with a CTCF ChIP-seq peak and a and weak SMC3 ChIP-seq peak, in a cluster of multiple TF binding sites and ENCODE ChIP-seq signals. There is also disruption of an ETS1 motif, which could play a role as negative or positive feedback on *ETS2* gene expression. For this occurrence, these two motifs map to the negative strand and are reported from 3’ to 5’. Three amplicons were designed for targeted bis-seq of the ASM region. **B**, Graphical representation of the targeted bis-seq results, validating a discrete ASM regions (amplicon 2) spanning ∼600 bp in T cells and lung. The relatively hypomethylated allele is the one with higher predicted SMC3 binding affinity. Numbers of ASM samples in each tissue and cell type, and additional annotations, are in **Table S2**. Rows represent sequence reads, and columns CpG sites in these reads. All samples are heterozygous, and the reads are separated by allele. Methylated CpGs are black circles, and unmethylated CpGs are white circles.

**Figure S10. Validations of ASM DMRs spanning a range of ASM ranks**

Targeted bis-seq showing validation of additional ASM regions (others in **Figures S7-S9**), with ASM index SNPs that have high, middle or low overall ranks. The results of all validations (27 ASM DMRs tested) are summarized in **Table S6**. Samples are heterozygous, and the reads are separated by allele. Methylated CpGs are black circles, and unmethylated CpGs are white circles. In each illustrated case, the relatively hypermethylated allele (REF or ALT) in the targeted bis-seq data is consistent with the relatively hypermethylated allele detected in the primary SureSelect or WGBS data (**Table S2**).

**Figure S11. Kernel density plots of methylation levels showing global hypomethylation and decrease in the percentage of high methylated CpGs in cancers**

Distribution of the averaged percentage of net methylation genome wide for all informative CpGs in cancer and lineage matched normal samples, by Kernel density estimation. As expected, CpG methylation has a bimodal distribution with a large major mode of high methylated CpGs (>80%) and a small minor mode of low methylated CpGs (<5% methylation; corresponding to CpG-islands) in the non-neoplastic cell type (B cells and glia). In multiple myeloma and lymphoma, prominent global hypomethylation with the loss of the high methylated CpGs peak is observed. Global hypomethylation is present, but milder, in GBMs.

**Figure S12. Replication of the major findings of this study using WGBS data from a single sequencing facility**

Gains of ASM are strongly correlated with genome-wide DNA hypomethylation, and ASM correlates with allele-specific binding affinities of specific classes of CTCF and TF recognition motifs in both cancer and normal samples, using WGBS data from a single sequencing facility (RPCI), which utilized a single library construction method. **A,** Data from WGBS performed at the RPCI facility, showing the relationship between global DNA methylation and the percentage of informative SNPs that reveal ASM in each sample. The results confirm a strong inverse correlation between per sample ASM frequencies and global methylation levels. Cancer samples (color-coded in red scale) have higher ASM frequencies than nearly all the non-cancer samples (color-coded in blue scale), except for overlap of the GBM data points with normal placental trophoblast and cultured bladder epithelial cells. When compared to lineage-matched normal cell types, there is a 7 to 12-fold increase in ASM frequencies in the three types of cancers. The EBV-immortalized but euploid GM12878 LCL shows global hypomethylation and a high frequency of ASM. **B,** Graphs showing significant correlations between allelic TF binding affinity scores and ASM in each of the 4 classes of ASM loci, using data from the RPCI sequencing facility. The left panel shows the fitted ASM difference on PWM score using a multivariate mixed model. The fitted line and its 95-confidence intervals (area) are shown for each ASM class; slopes were calculated by the marginal effects of the interaction term between PWM score and ASM class and were significantly different from zero. The correlations are similar in cancer ASM (in both non-desert and desert) compared to non-cancer ASM, with small differences in the slopes for each class. The right panel shows pairwise comparisons of the correlations in each of the 4 classes of ASM loci, Bonferroni-adjusted for multiple testing. While all the slopes are in a similar range, the correlations in the mixed model are weakest for cancer-only ASM loci, with a modest but statistically significant difference between the cancer vs non-cancer ASM classes, but not between desert and non-desert ASM loci. N: number of occurrences included in the mixed model. **C,** X-Y plots showing examples of TF motifs with strong correlations between predicted allele-specific binding site affinities (estimated by PWM scores) and methylation differences across all occurrences showing ASM, using data from the RPCI sequencing facility. Each data point represents one occurrence of the motif overlapping an ASM index SNP in cancer (orange) or non-cancer samples (blue). For occurrences showing ASM in multiple samples, allelic methylation differences were averaged across samples by sample type. R2 and B-H corrected p-values (FDR) were calculated using linear regression.

**Figure S13. Allele-specific losses of methylation leading to ASM in cancers**

Graphs showing the fitted values of the percentage methylation in cancer (red) for myeloma, lymphoma and glioblastoma versus the lineage-matched non-neoplastic cell types (B cells for myeloma and lymphoma and glia for glioblastoma) for regions where ASM was found only in cancer. In non-neoplastic cells, on average, the methylation levels in these regions were high or intermediate on both alleles and ASM in cancer reflects losses of methylation on one of the alleles. The average fractional methylation was estimated using a linear mixed model with random intercept and random slope (Methods). The light lines represent the fitted values for each locus and the bold line the average fit. The slope between low and high methylation estimates the ASM magnitude. The non-significant and small slope in non-neoplastic cells reflects the absence of significant ASM in these regions.

**Figure S14. Kernel density plots of methylation level distributions showing statistically enriched instances of allele-specific gains of methylation leading to ASM in cancers.**

These Kernel density plots show the distribution of methylation values in non-neoplastic cells comparing methylation at loci where ASM was found neither in the non-neoplastic cells nor in the matched cancer samples vs loci where ASM was found only in the matched cancer. These graphs show that allele-specific loss of methylation (LOM), which represents the most common scenario for cancer-only ASM, is numerically predominant but is nonetheless relatively under-represented compared to random expectation in the globally hypomethylated genomic background, while the less frequent allelic-specific gains of methylation (GOM) are over-represented relative to this background. As shown in Figure 3, these instances of GOM in the cancers often map to regions of poised chromatin.

**Figure S15. Shared ASM loci in cancer and non-cancer have similar ASM magnitude**

Graphs and diagrams showing the fitted values of the average percent methylation of the low and high methylated alleles in cancer (RED) for multiple myeloma (MM), lymphoma, and glioblastoma multiforme (GBM) vs cell lineage-matched non-neoplastic cell types (BLUE), namely B cells for MM and lymphoma and glia for GBM, for DMRs where ASM was found both in cancer and non-cancer. The average fractional methylation of each allele and in each cancer or normal sample class (middle panels) was estimated using a linear mixed model with random intercept and random slope (Methods).On the left panels, the light lines represent the fitted values for each locus and the bold line the average fit. The slopes between low and high methylated alleles estimate the ASM magnitude and are similar (parallel) in cancer and non-cancer samples, with a non-significant statistical interaction between cancer vs normal status and ASM magnitude. The right panels show primary WGBS data for representative examples, with sequence reads separated by allele. Methylated CpGs are black circles, and unmethylated CpGs are white circles.

**Figure S16. Correlations between allelic TF binding affinity scores and ASM cancer versus non cancer and specific examples of TF binding motifs, showing significant correlations between predicted allele-specific binding site affinities and ASM amplitude in both normal and cancer samples.**

**A,** Graphs showing significant correlations between allelic TF binding affinity scores and ASM in each of the 2 classes of ASM loci. The left panel shows the fitted ASM difference on PWM score using a multivariate mixed model. The fitted line and its 95-confidence intervals (area) are shown for each ASM class. The slopes of the fitted lines were calculated by the marginal effects of the interaction term between PWM score and ASM class and were significantly different from zero. The correlations are similar in cancer ASM compared to non-cancer ASM, with slightly weaker slope in cancer. The right panel shows the pairwise comparison of the correlations in each of the 2 classes of ASM loci with a significant difference between the cancer vs non-cancer ASM classes. N: number of occurrences included in the mixed model. **B**, These examples were selected requiring at least 3 occurrences per ASM classes. The X-Y plots show ASM magnitude vs differences in predicted allele-specific binding affinities (PWM scores) for the EHF_1, SPI1_3, SPIB_2 and ETV6_1 motifs. All 4 classes of ASM loci show similar anti-correlations, but there is a slight decrease in the slope for cancer compared to normal ASM. Desert vs non-desert classes of ASM loci show essentially identical slopes. Regression lines were not plotted if there were less than 3 occurrences within the ASM class (non-cancer/desert for SPIB_2 and cancer/desert for ETV6_1)

**Figure S17. Examples of ASM DMRs in chromatin deserts**

**A**, Map showing the ASM DMR tagged by index SNP rs2272697 in the body of the *MANBA* gene. ENCODE chromatin state data show that this region is marked only by a non-regulatory chromatin state (Txn, color-coded green), and the index SNP has a weak regulomeDB score (5), this SNP in fact disrupts multiple ETS-family binding motifs, suggesting that it could have a regulatory role via an ETS-family transcriptional pathway at some stage of cellular differentiation. This SNP is in high LD (R2>0.9) with multiple GWAS peak SNPs (rs5026472, rs1054037 and rs7665090) associated lymphocyte count, liver cirrhosis, and multiple sclerosis. **B,** Map showing the ASM DMR tagged by index SNP rs13097644 in the intergenic region upstream of the *SETMAR* gene. This region is flagged as quiescent by ENCODE chromatin state (color-coded light gray), consistent with the low regulomeDB score (6) for this SNP. However, the index ASM SNP disrupts an ASM-correlated Erg TF binding motif, suggesting that rs13097644 might act as a regulatory genetic variant at some stage of cell differentiation.

**Figure S18. Models for inter-individual variability and allele-switching at ASM loci**

**A**, ASM is not present, with high methylation on both alleles, when either the TFBS is not accessible (closed chromatin) or the TF is not sufficiently expressed (left panel). For accessible TFBS, the magnitude of ASM depends on the level of free TFs (middle panels). When the TF level is low, binding occurs on allele A (with high binding affinity) but stochastically in only a subset of DNA molecules. The overall proportion of low methylated reads (bound TFBS) reflects the steady state between dissociation and binding rates, defined by the concentration of the TF. At the other end of the concentration curve (right panel), strongly overexpressed TFs can bind both high and low binding affinity sites, leading to protection of both alleles against methylation and a loss of ASM. **B**, Inter-individual variability and allele-switching at ASM loci can be explained by a haplotype effects, in which multiple SNPs rather than a single SNP, or a dominant SNP in weak LD with the scored index SNP, dictate the ASM. This situation is “pseudo-switching”. **C,** Since most ASM SNPs found in this study can potentially disrupt multiple TF motifs, a TF competition model can explain bona fide allele-switching. This model appears to apply more often in cancer cells, which show a high frequency of ASM allele-switching in this study and are known to frequently over-express oncogenic TFs (e.g. c-MYC; Fig. 6).

**Figure S19. The percentage of ASM loci that show switching behavior in cancers is smaller when considering only loci for which ASM is also detected in non-cancer samples**

Graph showing the percentage of switching ASM loci in cancer and non-cancer samples as a function of the number of non-cancer samples where ASM is seen. For ASM loci in cancer, the x=0 data point corresponds to the percentage of switching among cancer-only ASM loci, while the subsequent data points show a decrease in switching among ASM loci found in both cancer and normal as the number of non-cancer samples (in addition to the cancer samples) showing ASM increases. As a comparison, the percentage of switching among ASM loci found in non-cancer samples is low and independent of the total number of samples showing ASM. This finding supports a working model that postulates two classes of binding motifs: one group in which disruptive SNPs show strong correlations with ASM, independently of the neoplastic cell phenotype, and stably bind their cognate factors, mitigating against allele switching; and another group of motifs with more labile TF binding, which are sensitive to global increases in chromatin accessibility and changes in intracellular levels of their cognate factors, leading to allele switching via “TF competition”.

**Figure S20. Examples of haplotype blocks defined by stringent and lenient parameters**

Example of haplotype blocks on chromosome 5 using the Gabriel et al. approach based on confidence interval of D’ values, with stringent (top) and lenient parameter (bottom). The lenient parameters, with relaxed D-prime confidence intervals and historical recombination rate (Methods), lead to haplotype blocks with larger sizes. Graphs were generated using Haploview using 1000 Genome data.

**Figure S21. Utility of D’ and R-square parameters for assessing candidate disease-associated rSNPs**

Example of D’ (left) and R^2^ (right) values between GWAS SNP rs710987 and all SNPs within 200 kb. The GWAS SNP is in red and ASM SNPs are in blue. The lenient haplotype block borders are shown in dashed green. The D’ graph confirms that most of the SNPs within the block (including the ASM SNPs) exhibit high D’ with the GWAS SNP and in this regard are in strong LD with it. The R^2^ graph of the same window and SNPs shows that only a small subset of the SNPs in LD also exhibits high R^2^ values, because even among SNPs in perfect LD only those with similar allele frequencies are expected to have high R^2^ values. A complete understanding of disease associations, including possible effects of more than one rSNP in the same haplotype block, requires extending the identification of rSNPs to those in strong LD with the GWAS peak SNP based on D’, even without high R^2^ values.

**Figure S22. Additional examples of mechanistically informative disease associated ASM index SNPs: autoimmune and neuropsychiatric disorders**

**A**, Map of the ASM region tagged by index SNP rs6603785 and located in an active enhancer region (color coded dark yellow) downstream of the *UBE2J2* gene on chromosome 1. ASM was observed in multiple blood cell types, including B cells. The ASM index SNP coincides with a GWAS peak SNP, associated with SLE (p=9.0×10^-6^; O.R.=1.11) and hypothyroidism (p=2.0×10^-9^; O.R. not listed). The SNP disrupts a MYC motif, with lower binding affinity and hypermethylation on the ALT allele, as predicted by the TF binding site occupancy model. **B**, Map of the ASM region tagged by index SNP rs2710323 and located in an active enhancer region (color coded in dark yellow) in the gene body of *ITIH1* on chromosome 3. ASM was observed in multiple blood cells, including T cells. The ASM index SNP coincides with a supra-threshold GWAS peak SNP for BMI measurements and multiple neuropsychiatric phenotypes including feeling nervous measurement, anxiety measurement, schizoaffective disorder, schizophrenia, and bipolar disorder (p-values and O.R. or Beta values in **Table S12**). The SNP disrupts an ELF1 motif, with lower binding affinity and higher methylation, as predicted, on the REF allele. For this occurrence, the motif maps the negative strand and is reported from 3’ to 5’.

**Figure S23. Additional examples of mechanistically informative disease associated ASM index SNPs: breast cancer and lymphoma**

**A**, Map of the ASM region tagged by index SNP rs61837215 and located in the active promoter region (color coded red) of the *SEPT7P9* pseudogene (nearest coding gene, *ZNF37A*) on chromosome 10. ASM is observed in multiple myeloma cells and in normal B cells. The index SNP is in strong LD with GWAS peak SNP rs2754412 associated with breast cancer (p=6.0×10^-7^; Beta=+.031). The ASM index SNP disrupts an ELF1_2 motif, with lower binding affinity and higher methylation on the REF allele, as predicted by the TF binding site occupancy model. **B**, Map of the ASM region tagged by SNP rs3806624 and located in a poised promoter region (color coded purple) of the *EOMES* gene on chromosome 3. ASM was observed in DLBCL and in GBM, with allele switching between the two cancer types. The ASM index SNP coincides with a GWAS peak SNP for Hodgkin lymphoma (p=1.0×10^-12^; O.R.=1.26) and is in strong LD with GWAS peak SNP rs9880772 associated with chronic lymphocytic leukemia (p=3.0×10^-11^; O.R.=1.19), as well as with multiple myeloma (**Table S11**). The SNP disrupts multiple motifs, including a BATF motif (lower binding affinity and higher methylation on the ALT allele) and a MAZ motif with opposite disruption of the binding affinity (lower binding affinity and higher methylation on the REF allele).

**Figure S24. ASM loci displayed as annotated genome browser tracks**

Track of high confidence ASM are provided in UCSC browser format (see Availability of Data). The detailed bed file is provided as supplemental data and can be uploaded to the UCSC Genome Browser. ASM is color coded in blue scale for negative direction ASM (hypomethylation of ALT allele compared to REF allele on average across all ASM samples) and positive direction ASM in red scale (hypermethylation of ALT allele compared to REF allele). Information about the index ASM SNP is displayed by clicking on the SNP (box). Reported information includes sample-aggregated information on the ASM-DMR and the index SNP, sample-specific information on ASM strength (p-value and methylation difference), the two classes of polymorphic motifs disrupted by the index SNP (i.e. enriched among ASM and/or with binding affinity-methylation correlation). Motif logo and sequences of the 2 alleles at the motif occurrences, generated using atSNP, are displayed by clicking on the motif name. Additional annotations of ASM index SNPs are in **Table S2**.

## Supplemental Table Legends

**Table S1. Biological samples analyzed in this study**

List of genomic DNA samples sequenced by Agilent SureSelect methyl-seq and by WGBS. Cell/tissue type, normal or cancer status, and diagnoses of the cancer patients from which the samples were collected are listed. Multiple biological samples from the same subjects can be identified by subject ID. Sequencing depth and QC information are also reported.

**Table S2. ASM index SNPs and DMRs identified in this study and annotated for multiple relevant parameters**

ASM loci, excluding known imprinted chromosomal regions (Methods), are listed using index SNPs as unique identifiers. Thus, some ASM DMRs are listed more than once, when multiple ASM index SNPs lie in the same DMR. Information relative to samples with and without ASM are aggregated, with the samples listed as concatenated entries. However, information at the single sample level can be retrieved from the UCSC-format detailed bed file: (https://genome.ucsc.edu/s/TyckoLab/High%20Confidence%20ASM). Relevant annotations were selected to characterize ASM SNPs based on their potential regulatory functions and disease associations. Briefly, the ASM index SNPs and DMRs are ranked for ASM strength and confidence (Methods), annotated using information about chromatin states, TF binding motifs, and allele-specific marks (ASB, eQTLs) from public databases, with some parameters being calculated using analytical procedures described in the Methods. ASM index SNPs are also annotated for their haplotype block locations and LD with GWAS peak SNPs.

**Table S3. Definitions of the terms in Table S2**

Description and definition of the columns in table S2. The order by row corresponds to the column order.

**Table S4. Imprinted regions with known ASM detected in this study**

ASM DMRs detected within 75 kb of known and validated imprinted genes (i.e. 150 kb windows) was considered as likely due to imprinting and was therefore excluded from our downstream analyses but listed in this table. The detection of these instances of ASM, with the expected high rate of allele switching (due to parent of origin dependence) in imprinted regions serves as a positive internal control for the initial steps of the ASM calling pipeline.

**Table S5. New candidate imprinted regions and previously provisional imprinted loci with ASM detected in this study**

**Table S6. ASM loci tested for validations by targeted bisulfite sequencing** Primers for bisulfite PCR were designed using MethPrimer, with the resulting amplicons spanning the indicated genomic coordinates.

**Table S7. Complete list of polymorphic CTCF and TF binding motifs found to be significantly enriched among ASM loci, requiring that the motif be disrupted by the ASM index SNP**

Results are from testing for enrichment of motif occurrences in which the motif is disrupted by the ASM index SNP, with a significant difference in affinity score between the two alleles. Significant enrichment among ASM loci was defined as FDR<0.05 and OR>2 (no depletion was observed). Background number of polymorphic occurrences for each motif (random expectation) was computed by screening a random sample of 40,000 occurrences from the list of non-ASM heterozygous (informative) SNPs in our study.

**Table S8. Complete list of CTCF and TF binding motifs that show significant correlations between allelic PWM scores and magnitude of ASM**

Significance was defined as FDR<0.05 and model R^2^ >0.4. Results from the linear regressions with and without controlling for the motif CpG content (when the number of occurrences was >3 in each group) are reported.

**Table S9. CTCF and TF binding motifs that show strong correlations of PWM scores with ASM and are also significantly enriched among ASM loci.**

Subset of enriched motifs with significant correlation of predicted allele-specific binding affinities with ASM magnitude. Results of the enrichment analysis by ASM classes (non-desert-non-neoplastic, desert-non-neoplastic, non-desert-cancer and non-desert-cancer) are reported.

**Table S10. ASM index SNPs in strong LD or precisely coinciding with GWAS peak SNPs for immune-related diseases and phenotypes**

The R^2^ cutoff was set at 0.8. GWAS peak SNPs associated with immune related diseases were identified using EFO parent-terms mapped to the GWAS reported traits (provided by the GWAS catalog) with additional manual curation.

**Table S11. ASM index SNPs in strong LD or precisely coinciding with GWAS peak SNPs for cancer susceptibility**

The R^2^ cutoff was set at 0.8. GWAS peak SNPs associated with cancer susceptibility were identified using EFO parent-terms mapped to the GWAS reported traits (provided by the GWAS catalog) with additional manual curation.

**Table S12. ASM index SNPs in strong LD or precisely coinciding with GWAS peak SNPs for brain-related diseases and phenotypes**

The R^2^ cutoff was set at 0.8. GWAS peak SNPs associated with neuropsychiatric disorders and traits, or with neurodegenerative diseases, were identified using EFO parent-terms mapped to the GWAS reported traits (provided by the GWAS catalog) with additional manual curation.

