## Supplemental Figures for "Allele-specific DNA methylation is increased in cancers and its dense mapping in normal plus neoplastic cells increases the yield of disease-associated regulatory SNPs"

### Figure S1A

#### Identifying and ranking ASM DMRs

**Sequence alignment to the reference methylome**

- BisMark - default settings with PE mode (WGBS, Agilent) and SE mode (Agilent unpaired reads after trimming).

**Heterozygous SNP calling on bisulfite-seq**

- BisSNP - with Quality Score recalibration and maximum coverage less then 200x.
- Non-G/A SNP coverage > 5x per allele (total coverage > 10x)
- Coverage of allele B between 20% and 80% of total coverage
- Filter out false calls : SNPs with multiple alignment, > 2 alleles with AF>0.01, indels, no AF (UCSC browser annotation of dbSNP147)
- Filter out false calls: SNPs with in HW disequilibrium (exact FDR<0.05) and het. freq > expected het. freq (dbSNP147)

**Identification of CpGs with ASM**

- CpG coverage > 5X per allele
- Filter out CpGs destroyed by common SNPs (> 5% MAF)
- Filter out CpGs within 10 bp of PE read 2 for Nextera ("fill-in" region) and 7 bp of both reads for TruSeq
- Fisher exact test comparing methylation on allele A vs B ($p < 0.05$)
- Check predicted differences in methylation in AA vs. BB homozygotes using an mQTL-like approach

**Identification & ranking of ASM DMRs**

- Estimate DMR border (first and last ASM CpG) and count the # of significant CpGs in the DMR
- Compare methylation between alleles across the DMR: avg methylation across all covered CpGs between the first and last ASM CpG of the same DMR.
- **Final criteria for calling ASM : DMR difference >20% and BH-corrected Wilcoxon p-value < 0.05, and at least 3 ASM CpGs including at least 2 consecutive ASM CpGs (overlapping DMRs merged) – see Fig. S1B**
- Exclude DMRs in known imprinted chromosomal regions
- Rank DMRs by absolute methylation difference, number and percentage of ASM CpGs
- Independent validations by targeted bis-seq on a set of ASM loci with strong and weak ranks

#### Testing mechanisms in normal and cancer ASM; nominating disease-associated rSNPs

**Functional annotation and enrichment analyses of features in ASM DMRs**

- eQTLs; DNAse-HS, TF binding (ChIP-seq)
- Disrupted TFBS motif occurrences overlapping a cognate ChIP-seq peak (200 bp window)
- TF peaks for which the motif is enriched (≥10 fold ) compared to background based on ENCODE ChIP-seq

**Identification of TFBS motifs with disruptive SNPs that correlate with ASM**

- Identify polymorphic TF motifs occurrences using ENCODE and JASPAR PWM and AtSNP software (require DNAse-HS peaks)
- Test for enrichment of disrupted vs non-disruptive polymorphic TF occurrences among ASM DMRs
- Test for correlations of ASM strength with PWM scores

**Assess mechanistic similarities and differences between cancer and non-cancer ASM**

- Multivariate analysis to compute odds ratios of finding ASM DMRs from cancer and non-cancer samples in specific chromatin states and associated with SNPs that disrupt specific TF binding motifs
- Assess rates of allele switching in cancer vs non-cancer ASM loci
- Assess frequencies of chromatin desert locations for cancer vs non-cancer ASM loci

**Identification of hap-ASM DMRs in haplotype blocks that contain GWAS peaks**

- Determine stringent and lenient LD and haplotype blocks
- Require distance between ASM and GWAS SNP < 200 kb
- Annotate ASM index SNPs for associations with immune/inflammatory, neuropsychiatric and neurodegenerative, neoplastic, and cardiometabolic diseases and traits

**Creation of genome browser tracks for visualization and prioritization of candidate disease-associated rSNPs**

- Custom tracks of ASM for each chromosome in UCSC Genome Browser format; tracks provide multiple annotations of each ASM index SNP.
- Annotations include ranking based on ASM strength and mechanistically relevant features including the identities of enriched and correlated TF and CTCF binding motifs that are disrupted by each ASM index SNP

**Figure S1B**

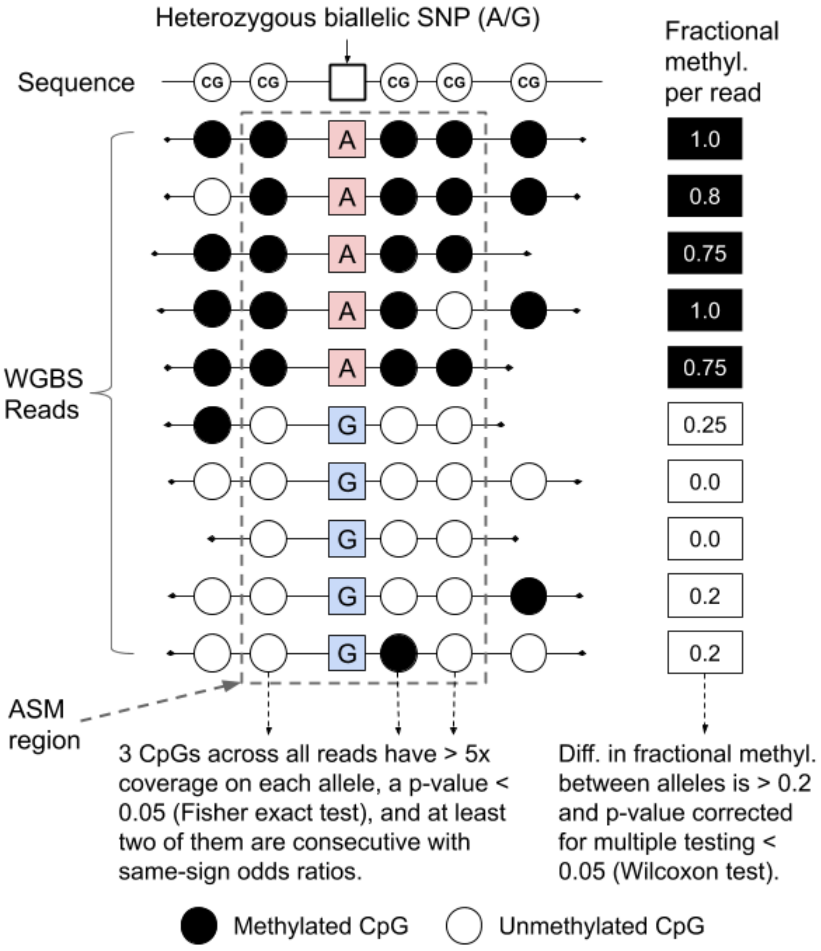

Heterozygous biallelic SNP (A/G)
Fractional methyl. per read
Sequence
CG
WGBS Reads
A
G
1.0
0.8
0.75
1.0
0.75
0.25
0.0
0.0
0.2
0.2
ASM region
3 CpGs across all reads have > 5x coverage on each allele, a p-value < 0.05 (Fisher exact test), and at least two of them are consecutive with same-sign odds ratios.
Diff. in fractional methyl. between alleles is > 0.2 and p-value corrected for multiple testing < 0.05 (Wilcoxon test).
Methylated CpG
Unmethylated CpG

#### Figure S2

**A**

**Agilent SureSelect (25 samples)**

- 1 lymphoblastoid cell line
- 24 primary non cancer tissues/cells

*9 brain cortex, 1 fetal lung, 2 fetal hearts, 1 fetal placenta, 2 livers, 3 PBL, 6 T cells*

**WGBS (81 samples)**

- 5 normal cell types from explants grown in tissue culture (*2 bladder epith. cell lines, 3 mammary epith. cell lines)*
- 1 lymphoblastoid cell line
- 59 primary non-neoplastic tissues or purified cell types (3 *brain cortex, 4 glia, 5 NeuN+ neurons, 9 B cells, 13 CD3+ T cells, 2 CD4+, 2 CD8+, 2 macrophage preps, 7 monocytes, 2 PBL, 1 LN, 1 fetal side placenta, 1 fetal side CTB, 1 maternal side CTB, 1 maternal side EVT, 3 livers, 2 breast tissue*)
- 16 primary cancers (7 *multiple myeloma, 2 DBCL, 1 FL, 6 GBM)*

Informative SNPs

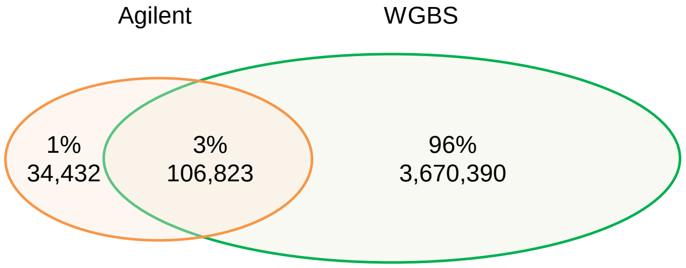


Informative SNPS in ≥ 2 biological samples

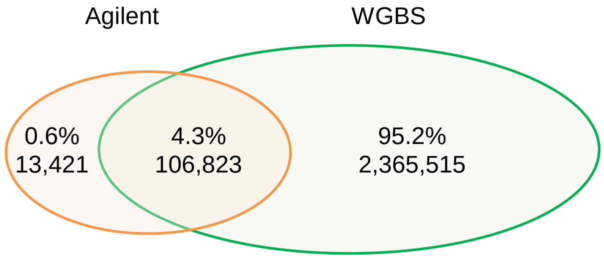


NB: The percentage are calculation on the union of Agilent and WGBS SNPs

**B**

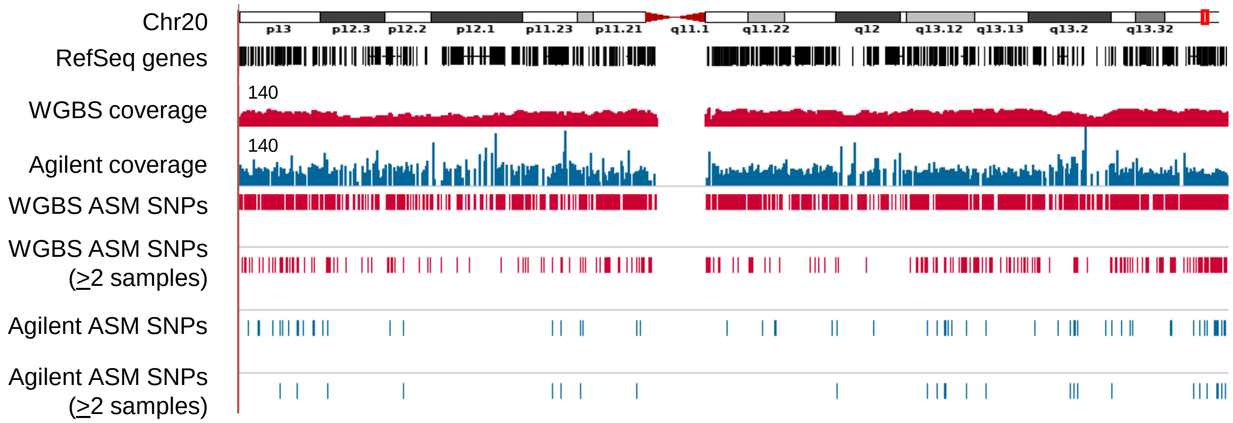

#### Figure S3

A

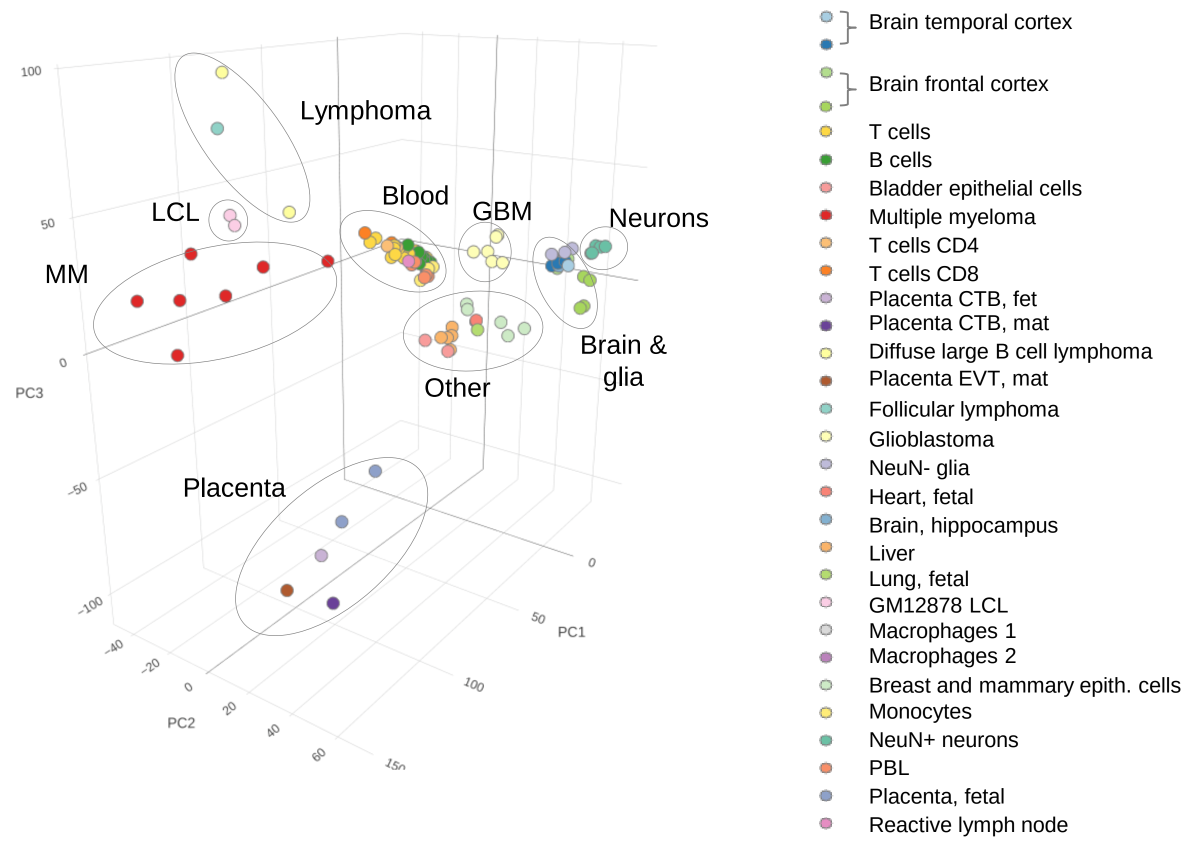


B

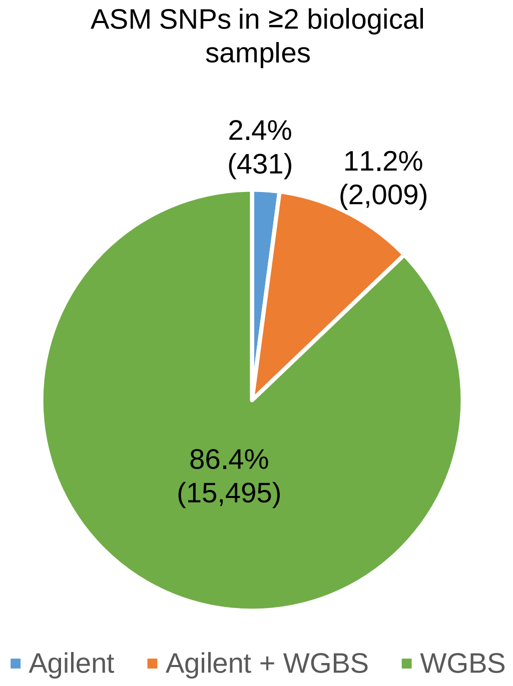


C

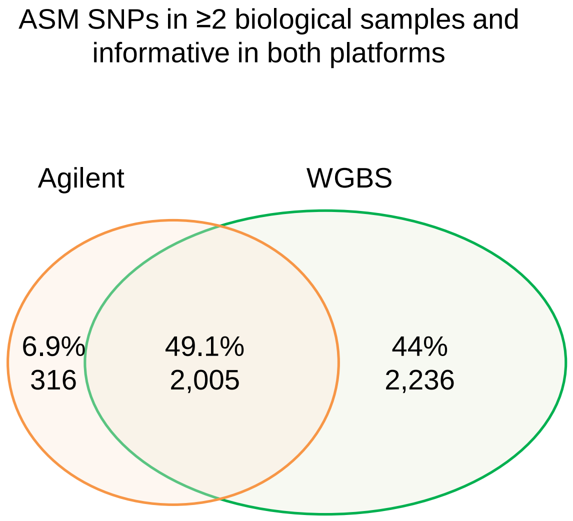

**Figure S4**

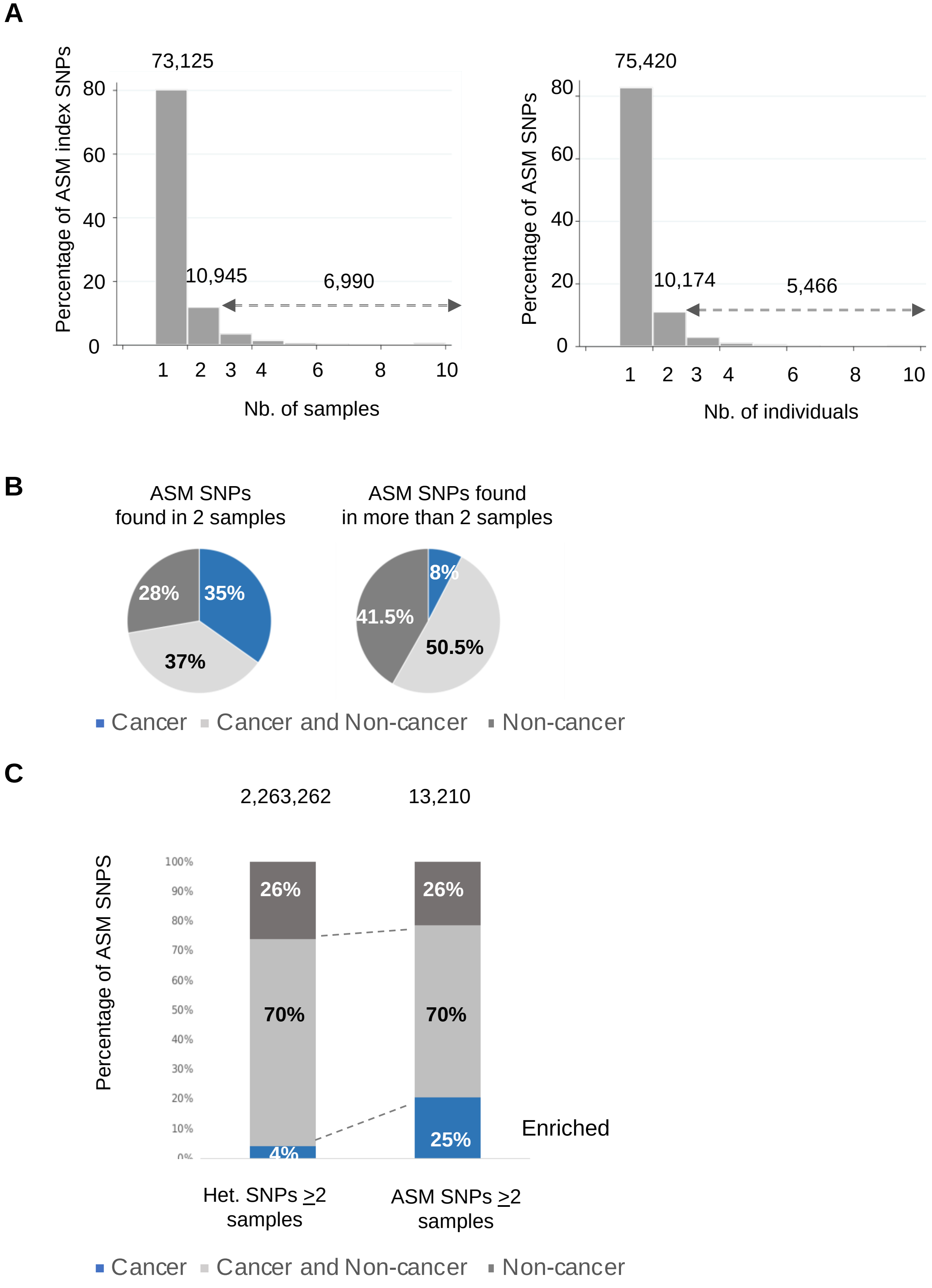
A
73,125
Percentage of ASM index SNPs
80
60
40
20
0
10,945
6,990
1 2 3 4 6 8 10
Nb. of samples
75,420
Percentage of ASM SNPs
80
60
40
20
0
10,174
5,466
1 2 3 4 6 8 10
Nb. of individuals
B
ASM SNPs
found in 2 samples
28%
35%
37%
ASM SNPs found
in more than 2 samples
8%
41.5%
50.5%
Cancer
Cancer and Non-cancer
Non-cancer
C
2,263,262
13,210
Percentage of ASM SNPS
100%
90%
80%
70%
60%
50%
40%
30%
20%
10%
0%
26%
70%
4%
26%
70%
25%
Enriched
Het. SNPs ≥2 samples
ASM SNPs ≥2 samples
Cancer
Cancer and Non-cancer
Non-cancer

#### Figure S5

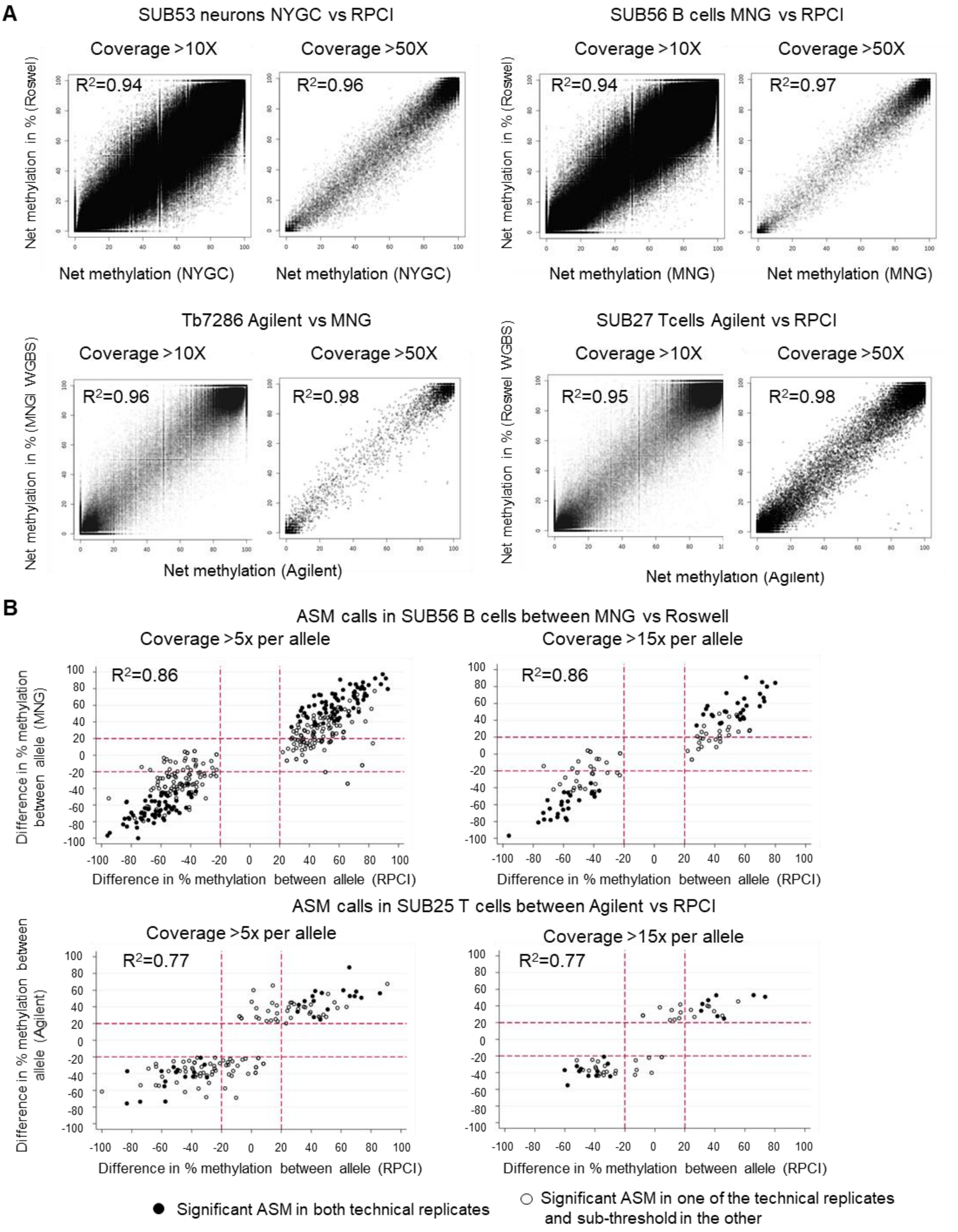

A
SUB53 neurons NYGC vs RPCI
SUB56 B cells MNG vs RPCI
Coverage >10X
Coverage >50X
R2=0.94
R2=0.96
R2=0.94
R2=0.97
Net methylation in % (Roswel)
Net methylation (NYGC)
Net methylation (MNG)
Tb7286 Agilent vs MNG
SUB27 Tcells Agilent vs RPCI
R2=0.96
R2=0.98
R2=0.95
R2=0.98
Net methylation in % (MNGI WGBS)
Net methylation in % (Roswel WGBS)
Net methylation (Agilent)
B
ASM calls in SUB56 B cells between MNG vs Roswell
Coverage >5x per allele
Coverage >15x per allele
R2=0.86
R2=0.86
Difference in % methylation between allele (MNG)
Difference in % methylation between allele (RPCI)
ASM calls in SUB25 T cells between Agilent vs RPCI
R2=0.77
R2=0.77
Difference in % methylation between allele (Agilent)
Significant ASM in both technical replicates
Significant ASM in one of the technical replicates and sub-threshold in the other

### Figure S6

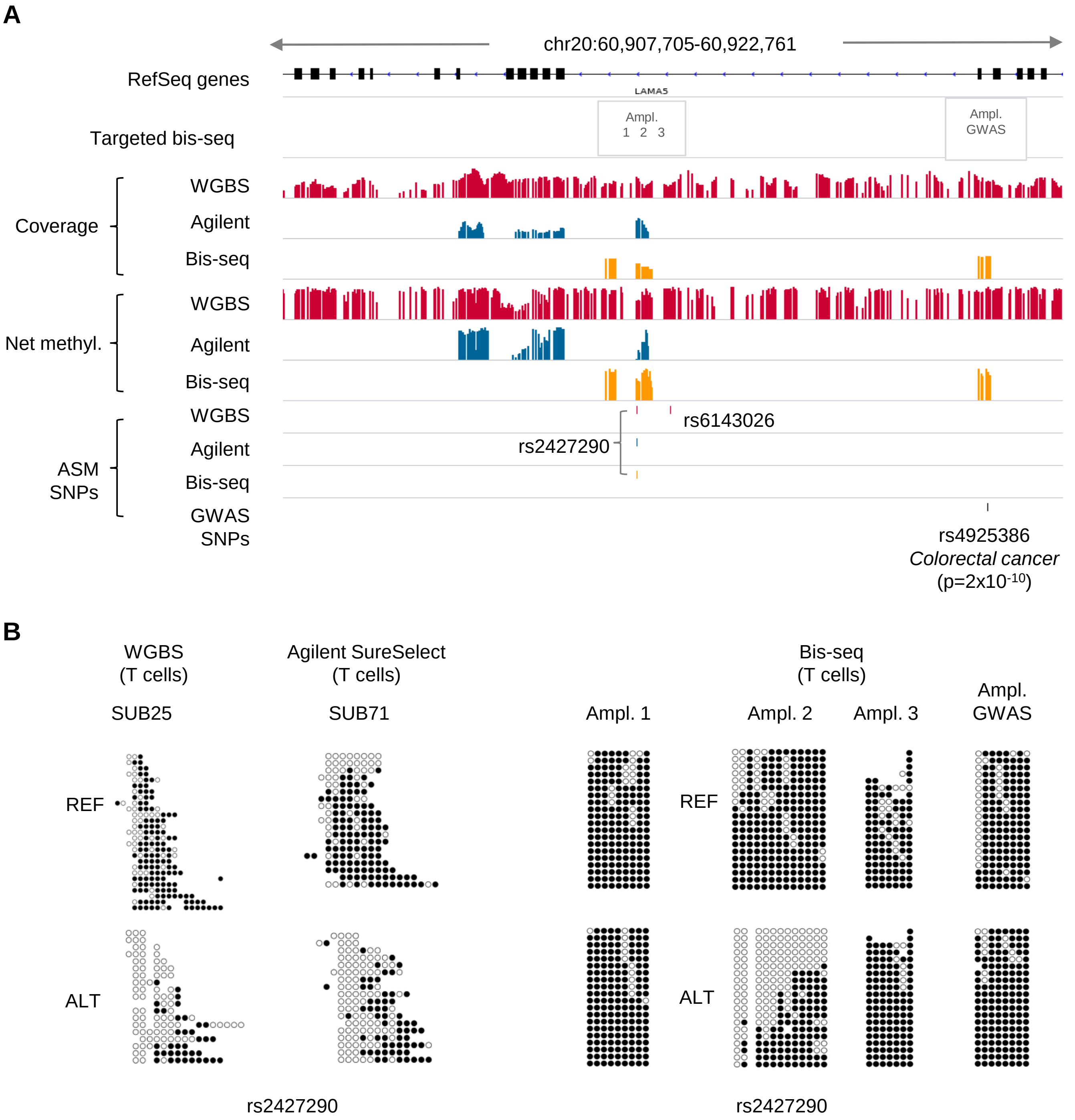
A
chr20:60,907,705-60,922,761
RefSeq genes
LAMA5
Targeted bis-seq
Ampl.
1 2 3
Ampl.
GWAS
Coverage
WGBS
Agilent
Bis-seq
Net methyl.
WGBS
Agilent
Bis-seq
ASM SNPs
WGBS
Agilent
Bis-seq
GWAS SNPs
rs6143026
rs2427290
rs4925386
Colorectal cancer
(p=2x10-10)
B
WGBS
(T cells)
SUB25
Agilent SureSelect
(T cells)
SUB71
Bis-seq
(T cells)
Ampl. 1
Ampl. 2
Ampl. 3
Ampl.
GWAS
REF
ALT
REF
ALT
rs2427290
rs2427290

**Figure S7**

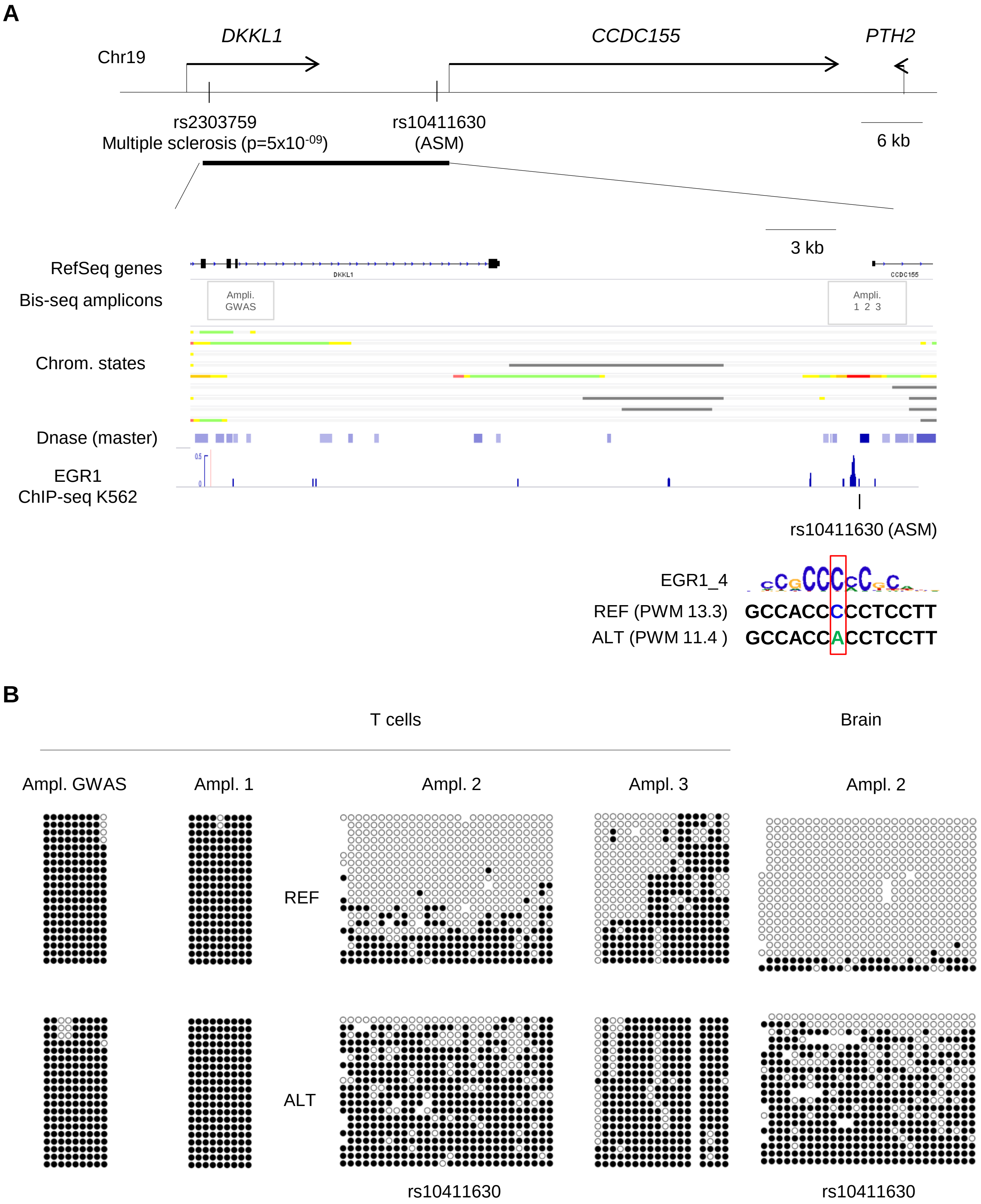
A
DKKL1
CCDC155
PTH2
Chr19
rs2303759
Multiple sclerosis (p=5x10-09)
rs10411630
(ASM)
6 kb
3 kb
RefSeq genes
DKKL1
CCDC155
Bis-seq amplicons
Ampli. GWAS
Ampli. 1 2 3
Chrom. states
Dnase (master)
EGR1
ChIP-seq K562
rs10411630 (ASM)
EGR1_4
REF (PWM 13.3)
GCCACCCCCTCCTT
ALT (PWM 11.4 )
GCCACCACCTCCTT
B
T cells
Brain
Ampl. GWAS
Ampl. 1
Ampl. 2
Ampl. 3
Ampl. 2
REF
ALT
rs10411630
rs10411630

#### Figure S8

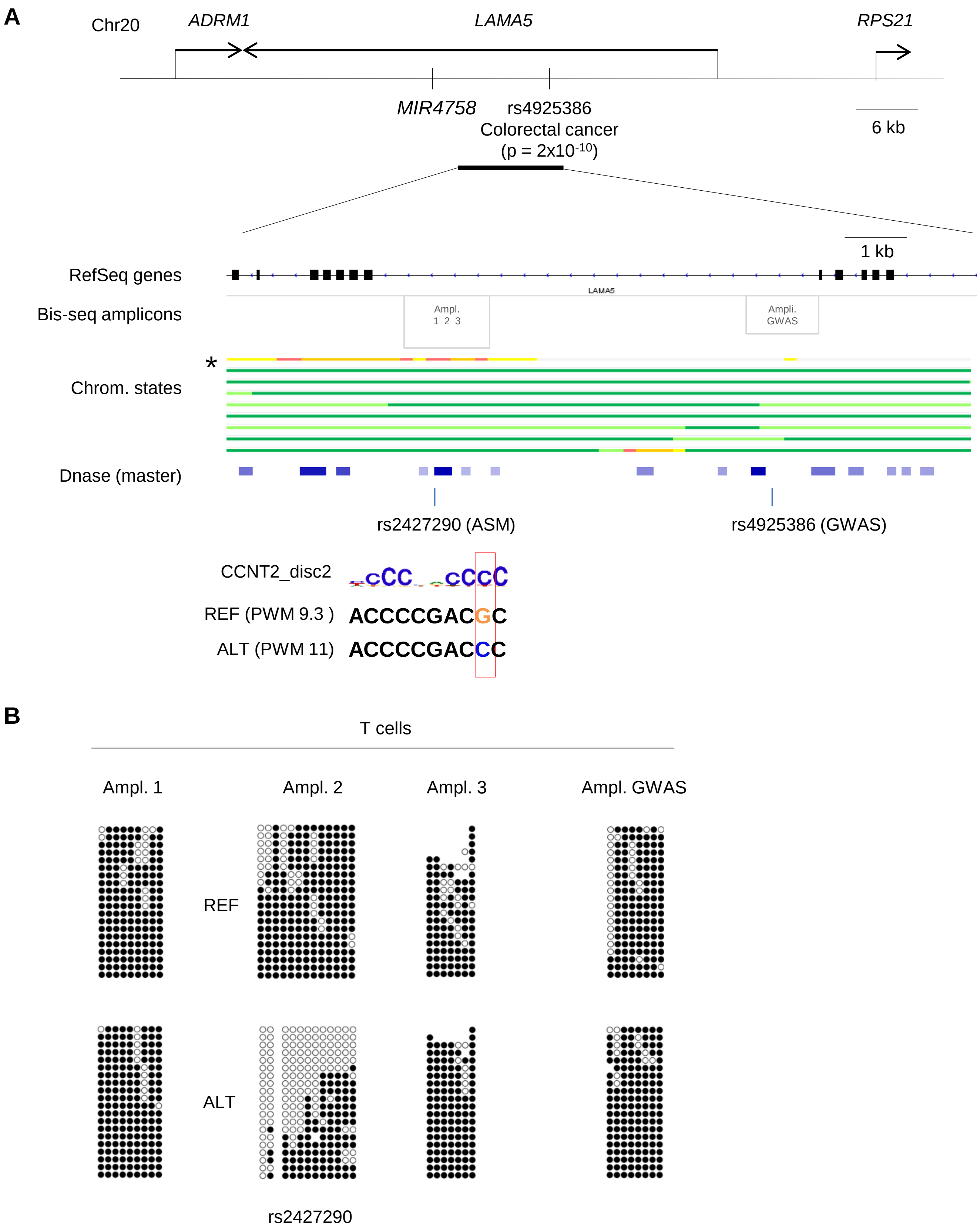
A
Chr20
ADRM1
LAMA5
RPS21
MIR4758
rs4925386
Colorectal cancer
(p = 2x10-10)
6 kb
1 kb
RefSeq genes
LAMA5
Bis-seq amplicons
Ampl. 1 2 3
Ampli. GWAS
*
Chrom. states
Dnase (master)
rs2427290 (ASM)
rs4925386 (GWAS)
CCNT2_disc2
REF (PWM 9.3 )
ACCCCGACGC
ALT (PWM 11)
ACCCCGACCC
B
T cells
Ampl. 1
Ampl. 2
Ampl. 3
Ampl. GWAS
REF
ALT
rs2427290

#### Figure S9

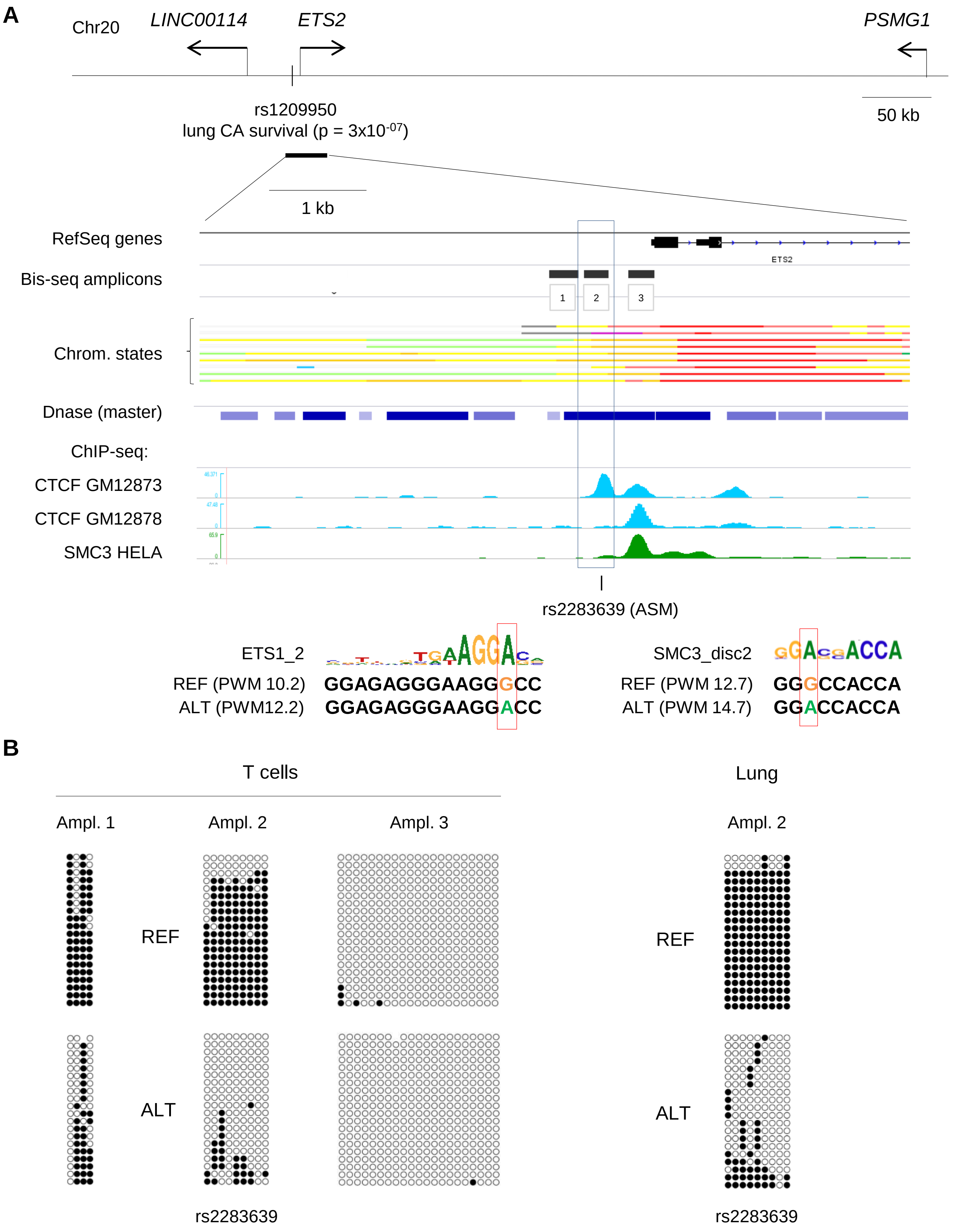

A
Chr20
LINC00114
ETS2
PSMG1
rs1209950
lung CA survival (p = 3x10-07)
50 kb
1 kb
RefSeq genes
ETS2
Bis-seq amplicons
1
2
3
Chrom. states
Dnase (master)
ChIP-seq:
CTCF GM12873
CTCF GM12878
SMC3 HELA
rs2283639 (ASM)
ETS1_2
REF (PWM 10.2)
GGAGAGGGAAGGGCC
ALT (PWM12.2)
GGAGAGGGAAGGACC
SMC3_disc2
REF (PWM 12.7)
GGGCCACCA
ALT (PWM 14.7)
GGACCACCA
B
T cells
Lung
Ampl. 1
Ampl. 2
Ampl. 3
Ampl. 2
REF
ALT
REF
ALT
rs2283639
rs2283639

### Figure S10

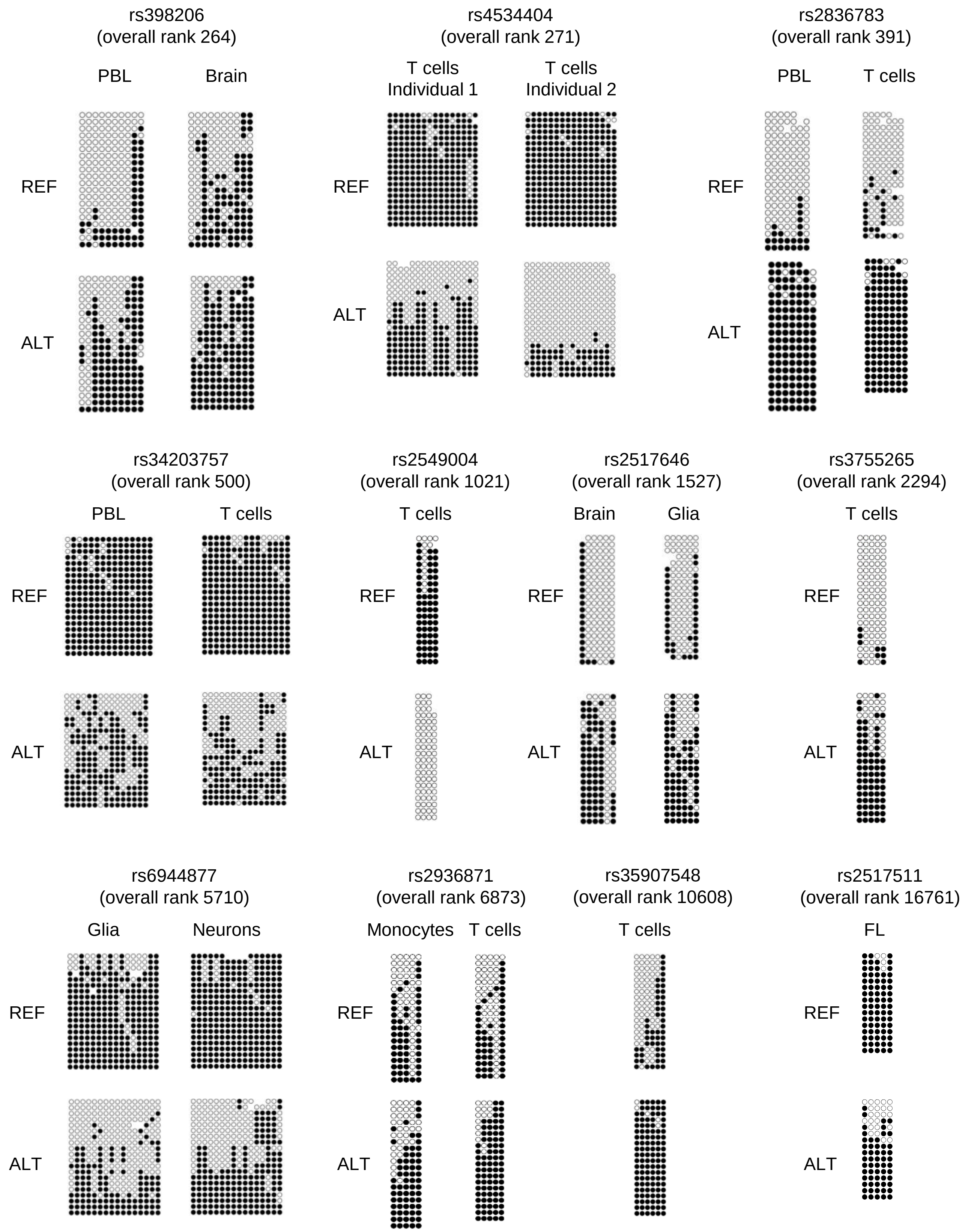


Complete list of tested loci is in Table S6

### Figure S11

Distribution of net methylation in B cells
vs Myeloma

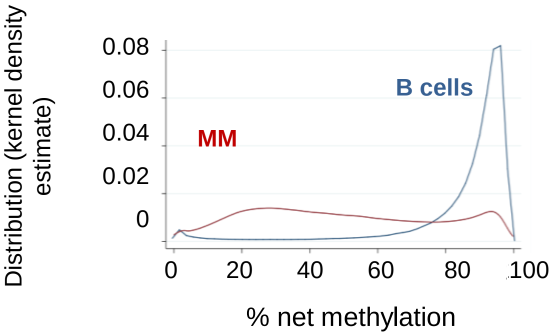


Distribution of net methylation in
B cells vs lymphoma

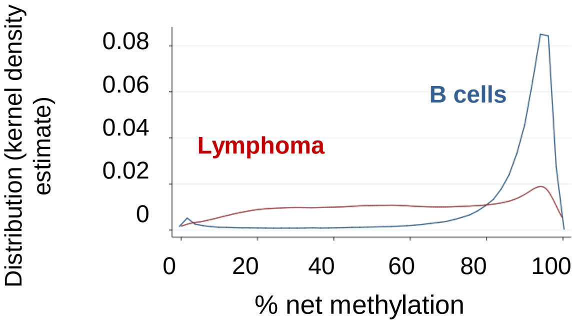


Distribution of net methylation in glia
vs GBM

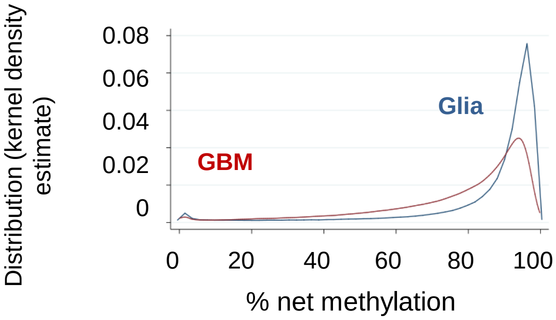

**Figure S12**: results based on data from a single sequencing facility (RPCI), using a single library construction method.

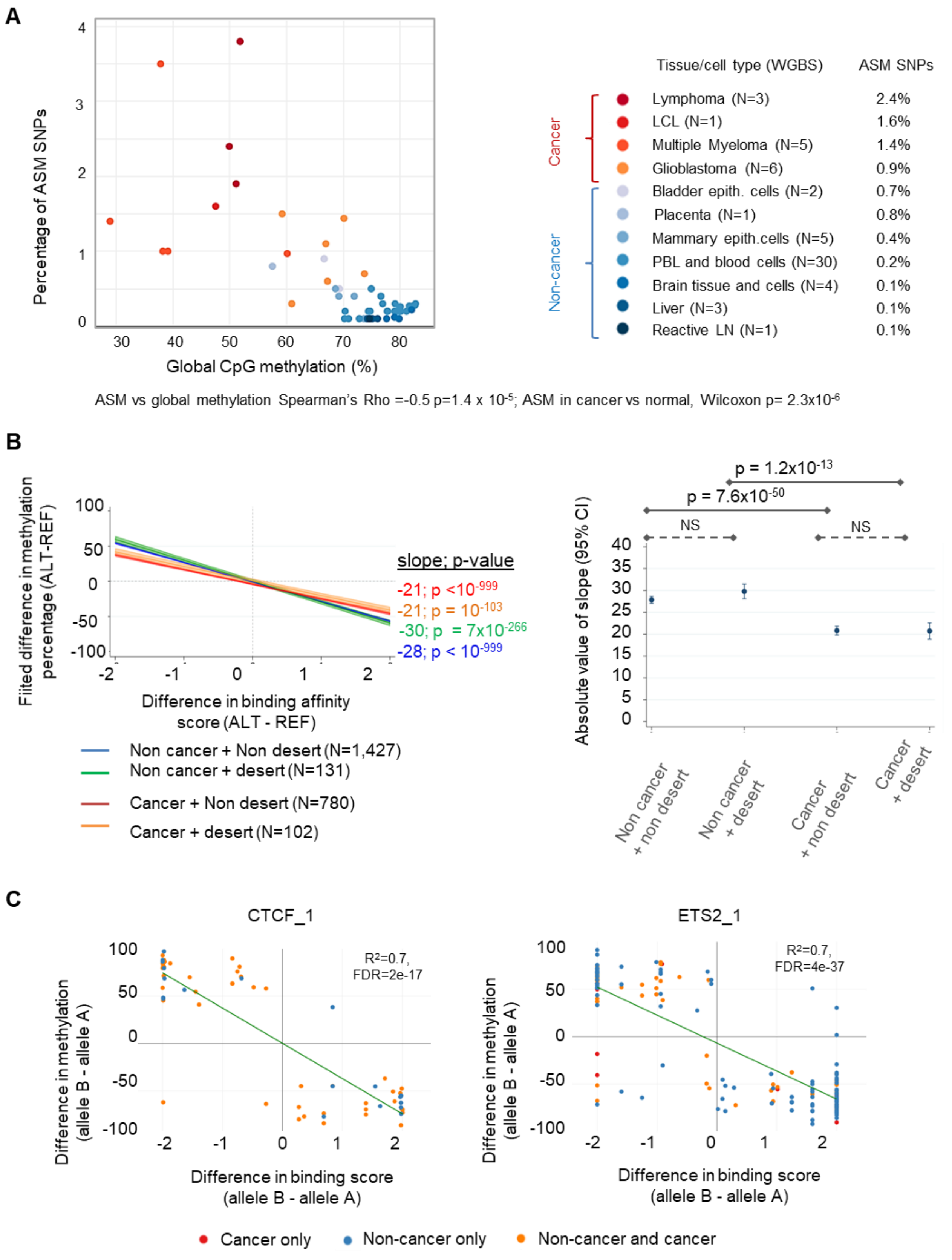

**Figure S13**

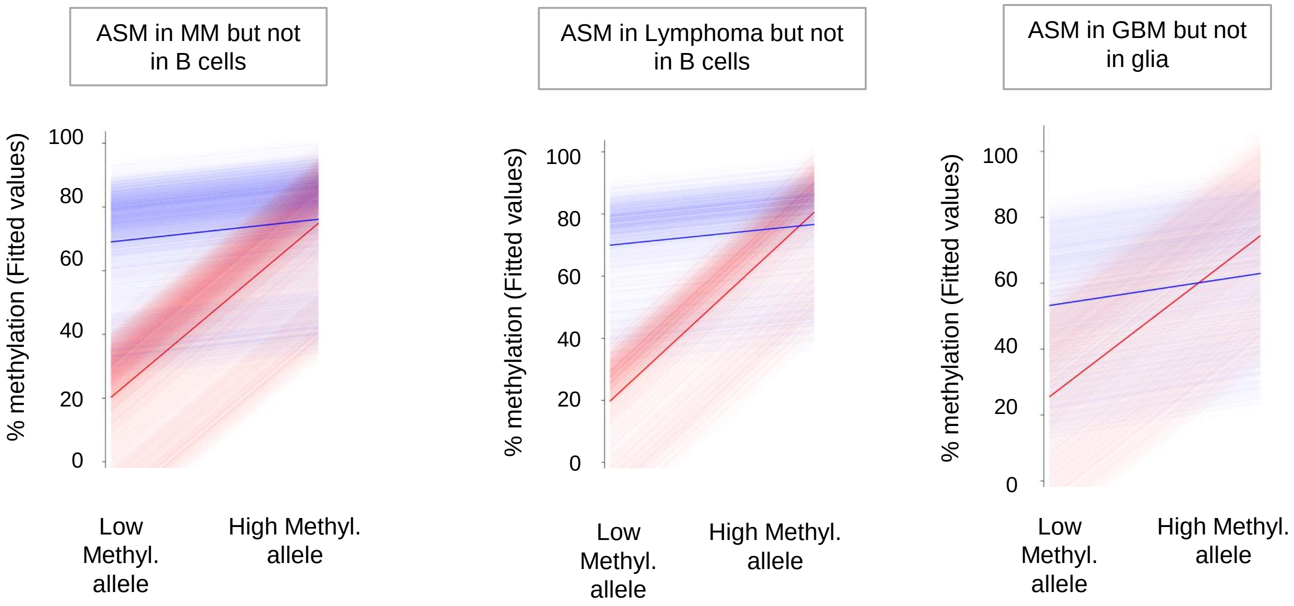
ASM in MM but not in B cells
% methylation (Fitted values)
100
80
60
40
20
0
Low Methyl. allele
High Methyl. allele
ASM in Lymphoma but not in B cells
% methylation (Fitted values)
100
80
60
40
20
0
Low Methyl. allele
High Methyl. allele
ASM in GBM but not in glia
% methylation (Fitted values)
100
80
60
40
20
0
Low Methyl. allele
High Methyl. allele

**Figure S14**

Distribution of net methylation in B cells - specifically for loci that show ASM in myeloma (BLUE) vs loci without ASM (RED)

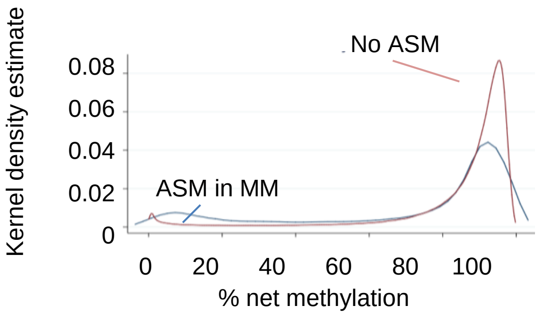


Over-representation of low methylation (OR: 4.1, $p< 10^{-999}$)

Under-representation of high methylation (OR: 0.3, $p=6x10^{-290}$)

Distribution of net methylation in B cells - specifically for loci that show ASM in lymphoma (BLUE) vs loci without ASM (RED)

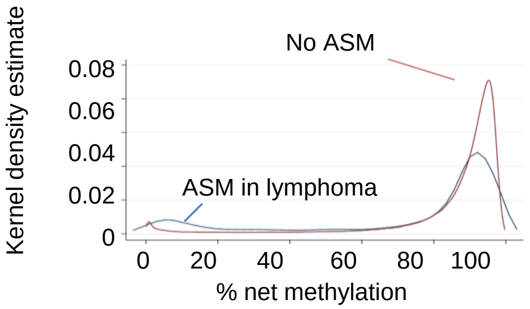


Over-representation of low methylation (OR: 3.9, $p=2x10^{-146}$)

Under-representation of high methylation (OR: 0.3, $p= 10^{-133}$)

Distribution of net methylation in glia - specifically for loci that show ASM in GBM (BLUE) vs loci without ASM (RED)

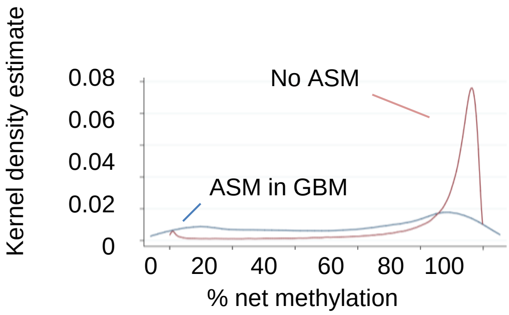


Over-representation of low methylation (OR: 6.9, $p<10^{-999}$)

Under-representation of high methylation (OR: 0.1, $p<10^{-999}$)

#### Figure S15

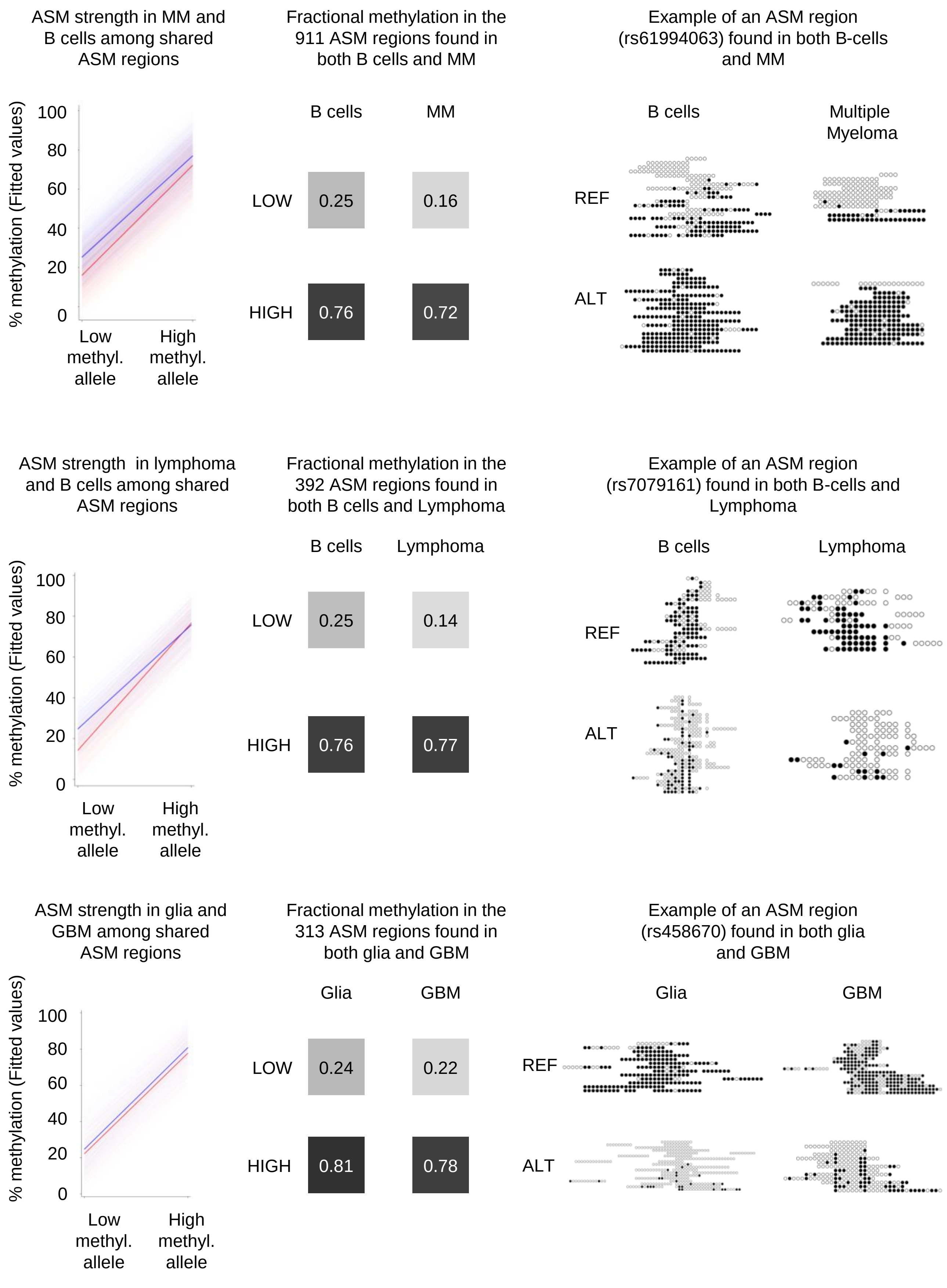
ASM strength in MM and B cells among shared ASM regions
% methylation (Fitted values)
100
80
60
40
20
0
Low methyl. allele
High methyl. allele
Fractional methylation in the 911 ASM regions found in both B cells and MM
B cells
MM
LOW
0.25
0.16
HIGH
0.76
0.72
Example of an ASM region (rs61994063) found in both B-cells and MM
B cells
Multiple Myeloma
REF
ALT
ASM strength in lymphoma and B cells among shared ASM regions
% methylation (Fitted values)
100
80
60
40
20
0
Low methyl. allele
High methyl. allele
Fractional methylation in the 392 ASM regions found in both B cells and Lymphoma
B cells
Lymphoma
LOW
0.25
0.14
HIGH
0.76
0.77
Example of an ASM region (rs7079161) found in both B-cells and Lymphoma
B cells
Lymphoma
REF
ALT
ASM strength in glia and GBM among shared ASM regions
% methylation (Fitted values)
100
80
60
40
20
0
Low methyl. allele
High methyl. allele
Fractional methylation in the 313 ASM regions found in both glia and GBM
Glia
GBM
LOW
0.24
0.22
HIGH
0.81
0.78
Example of an ASM region (rs458670) found in both glia and GBM
Glia
GBM
REF
ALT

**Figure S16**

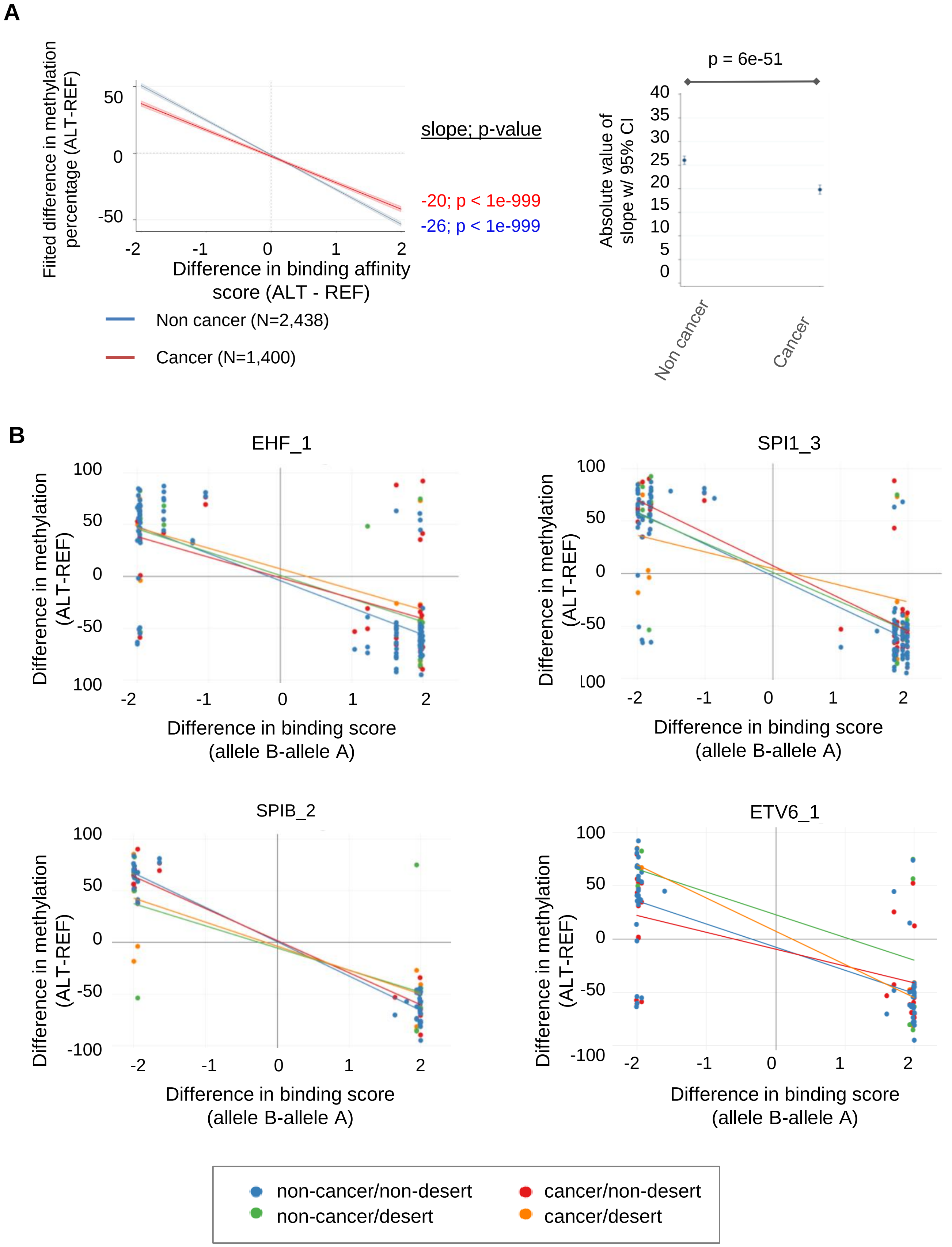
A
Fitted difference in methylation percentage (ALT-REF)
50
0
-50
-2
-1
0
1
2
Difference in binding affinity score (ALT - REF)
slope; p-value
-20; p < 1e-999
-26; p < 1e-999
Non cancer (N=2,438)
Cancer (N=1,400)
p = 6e-51
Absolute value of slope w/ 95% CI
40
35
30
25
20
15
10
5
0
Non cancer
Cancer
B
EHF_1
SPI1_3
SPIB_2
ETV6_1
Difference in methylation (ALT-REF)
100
50
0
-50
-100
Difference in binding score (allele B-allele A)
non-cancer/non-desert
non-cancer/desert
cancer/non-desert
cancer/desert

### Figure S17

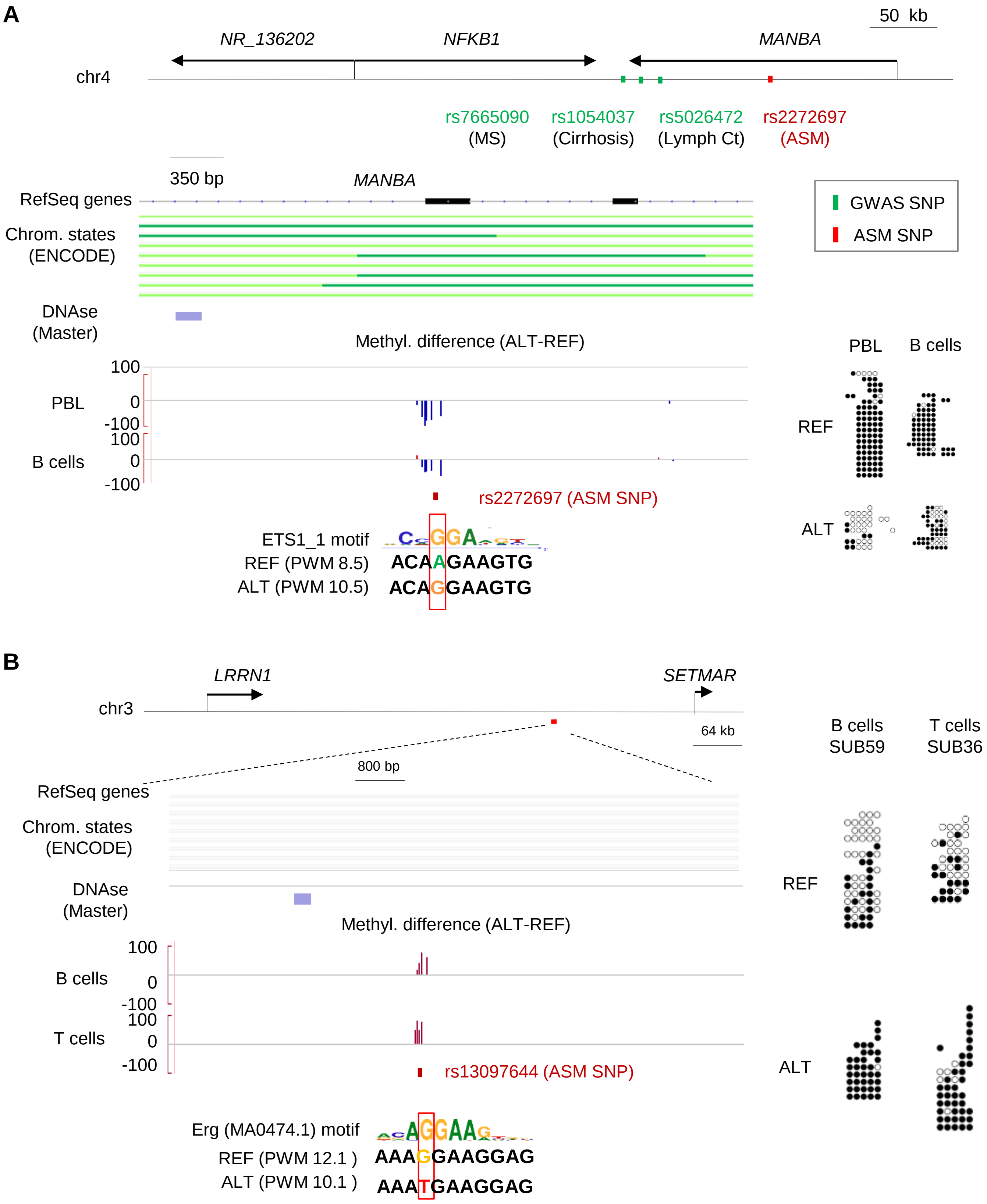
A
50 kb
NR_136202
NFKB1
MANBA
chr4
rs7665090
(MS)
rs1054037
(Cirrhosis)
rs5026472
(Lymph Ct)
rs2272697
(ASM)
350 bp
MANBA
RefSeq genes
Chrom. states
(ENCODE)
GWAS SNP
ASM SNP
DNAse
(Master)
Methyl. difference (ALT-REF)
PBL
B cells
100
0
-100
rs2272697 (ASM SNP)
REF
ALT
ETS1_1 motif
REF (PWM 8.5)
ALT (PWM 10.5)
ACAAGAAGTG
ACAGGAAGTG
B
LRRN1
SETMAR
chr3
64 kb
800 bp
B cells
SUB59
T cells
SUB36
RefSeq genes
Chrom. states
(ENCODE)
DNAse
(Master)
Methyl. difference (ALT-REF)
B cells
T cells
100
0
-100
rs13097644 (ASM SNP)
REF
ALT
Erg (MA0474.1) motif
REF (PWM 12.1 )
ALT (PWM 10.1 )
AAAGGAAGGAG
AAATGAAGGAG

**Figure S18**

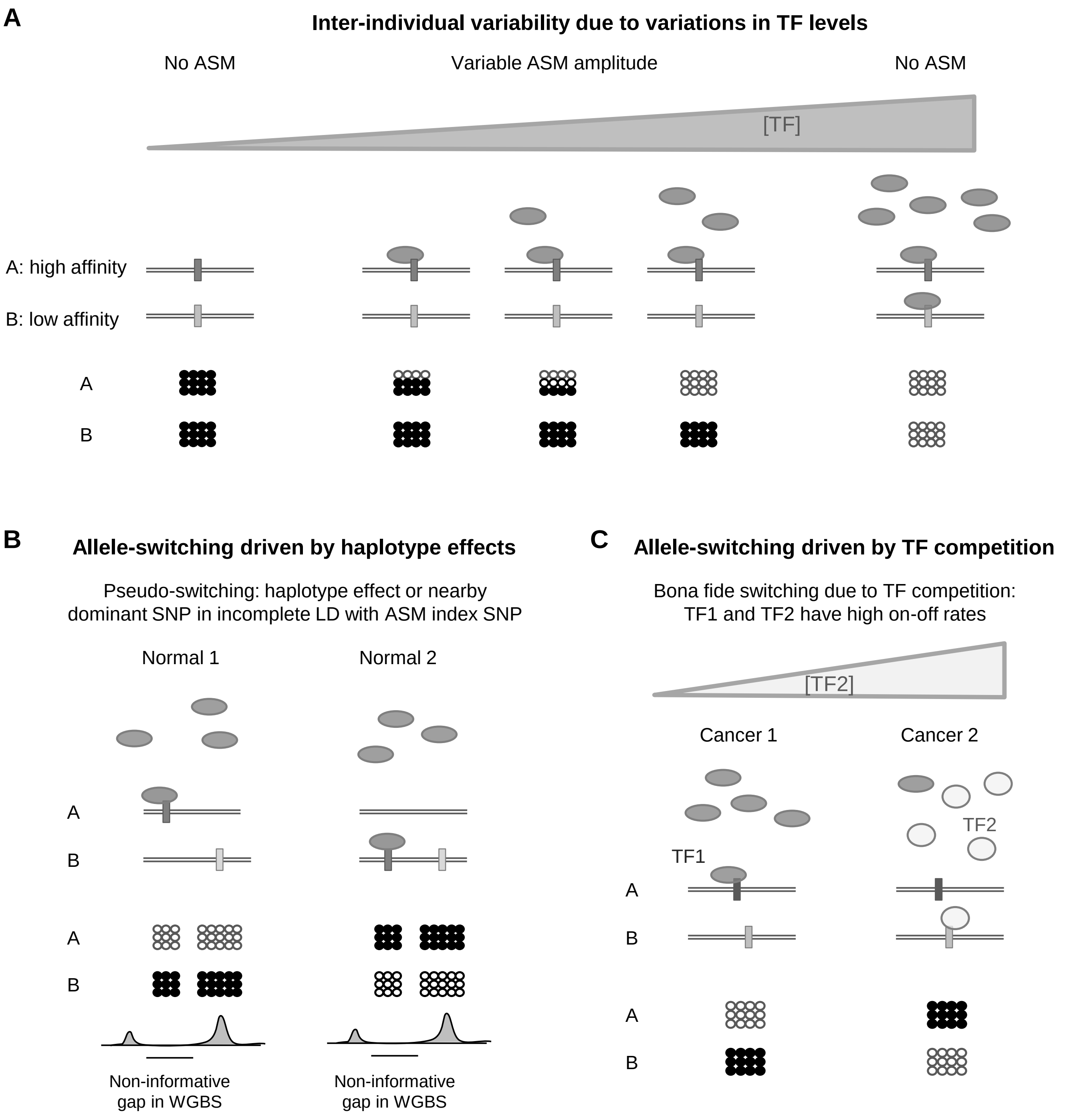
A
Inter-individual variability due to variations in TF levels
No ASM
Variable ASM amplitude
No ASM
[TF]
A: high affinity
B: low affinity
A
B
B
Allele-switching driven by haplotype effects
Pseudo-switching: haplotype effect or nearby dominant SNP in incomplete LD with ASM index SNP
Normal 1
Normal 2
A
B
A
B
Non-informative gap in WGBS
Non-informative gap in WGBS
C
Allele-switching driven by TF competition
Bona fide switching due to TF competition: TF1 and TF2 have high on-off rates
[TF2]
Cancer 1
Cancer 2
TF2
TF1
A
B
A
B

**Figure S19**

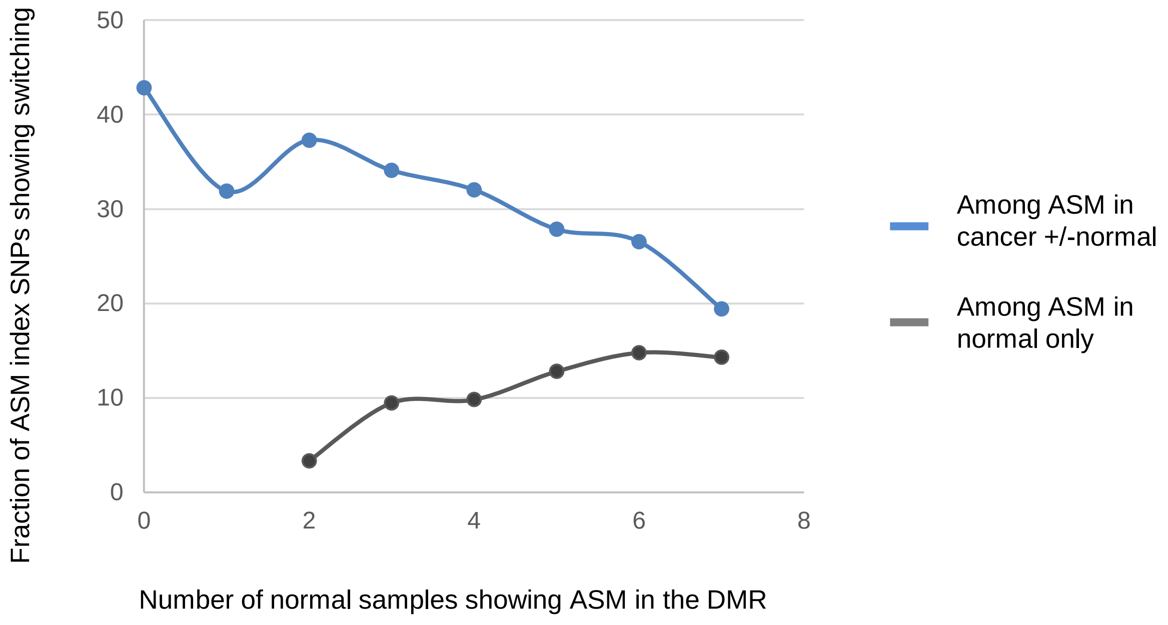

Fraction of ASM index SNPs showing switching
50
40
30
20
10
0
0
2
4
6
8
Number of normal samples showing ASM in the DMR
Among ASM in cancer +/-normal
Among ASM in normal only

#### Figure S20

Gabriel et al. (stringent) criteria

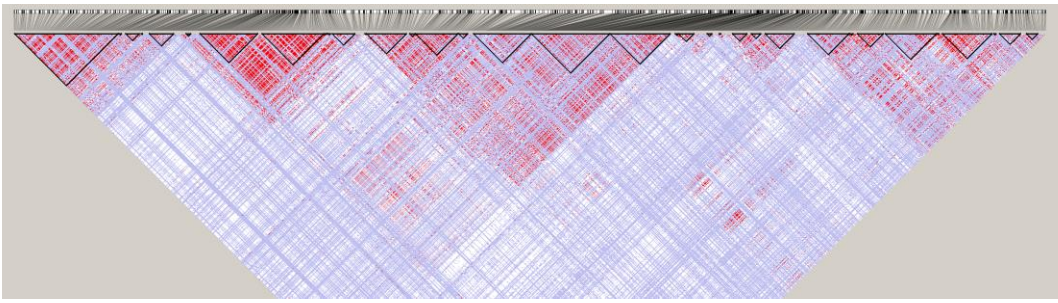

“Relaxed” criteria

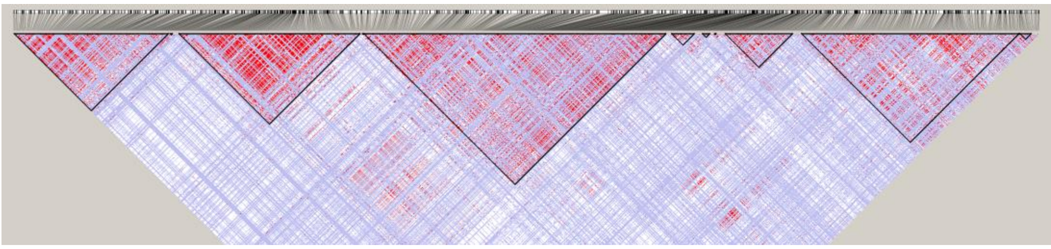

#### Figure S21

**A**

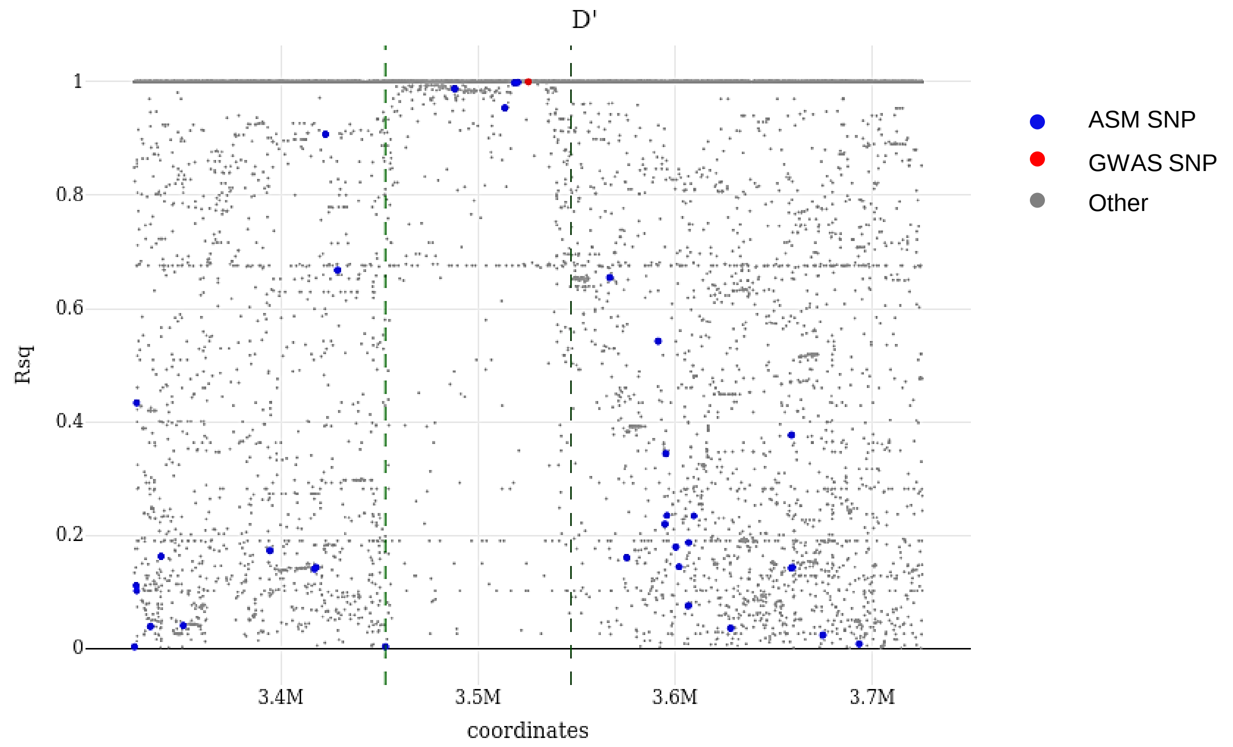


**B**

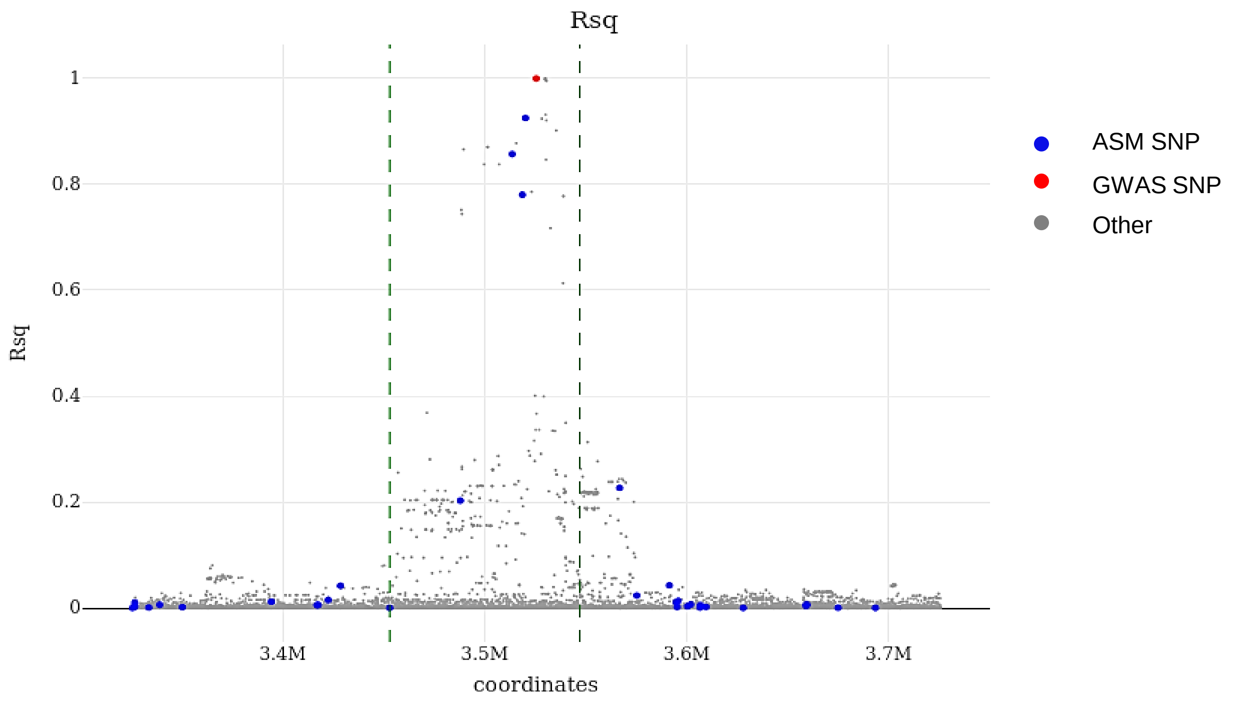

#### Figure S22

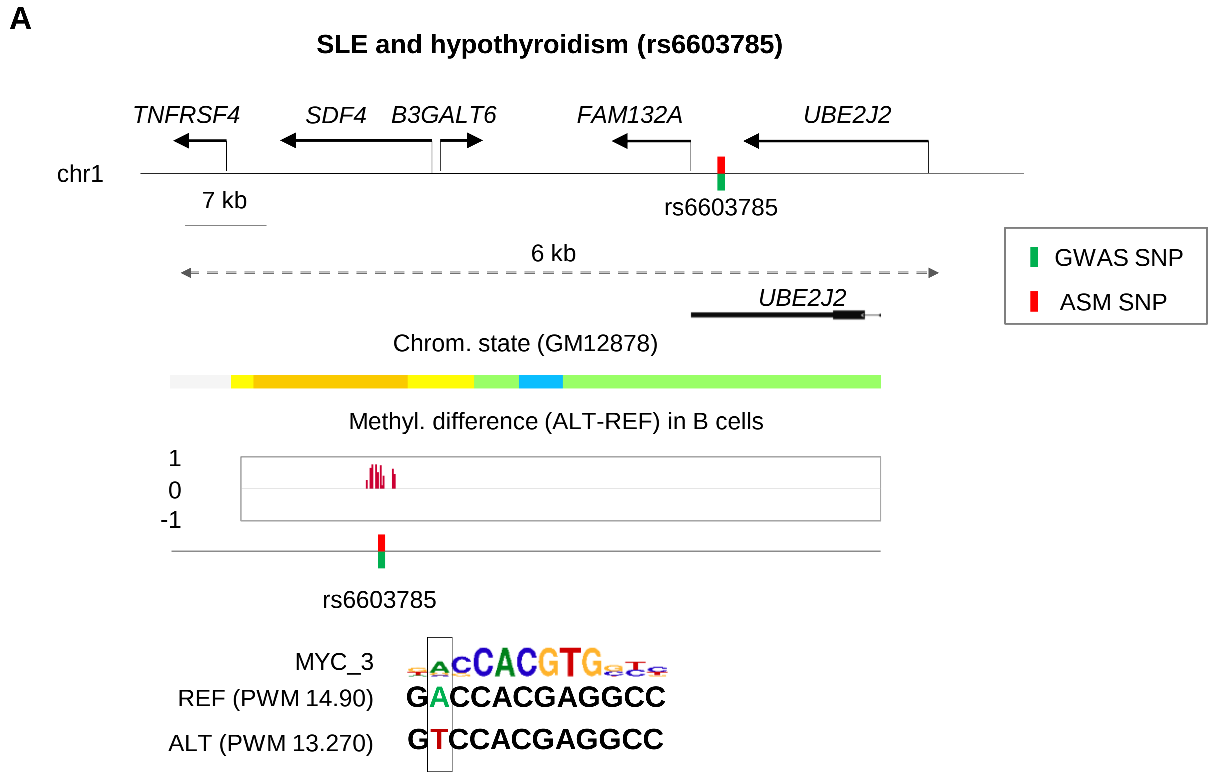
A
SLE and hypothyroidism (rs6603785)
TNFRSF4
SDF4
B3GALT6
FAM132A
UBE2J2
chr1
7 kb
rs6603785
6 kb
UBE2J2
GWAS SNP
ASM SNP
Chrom. state (GM12878)
Methyl. difference (ALT-REF) in B cells
1
0
-1
rs6603785
MYC_3
REF (PWM 14.90) GACCACGAGGCC
ALT (PWM 13.270) GTCCACGAGGCC

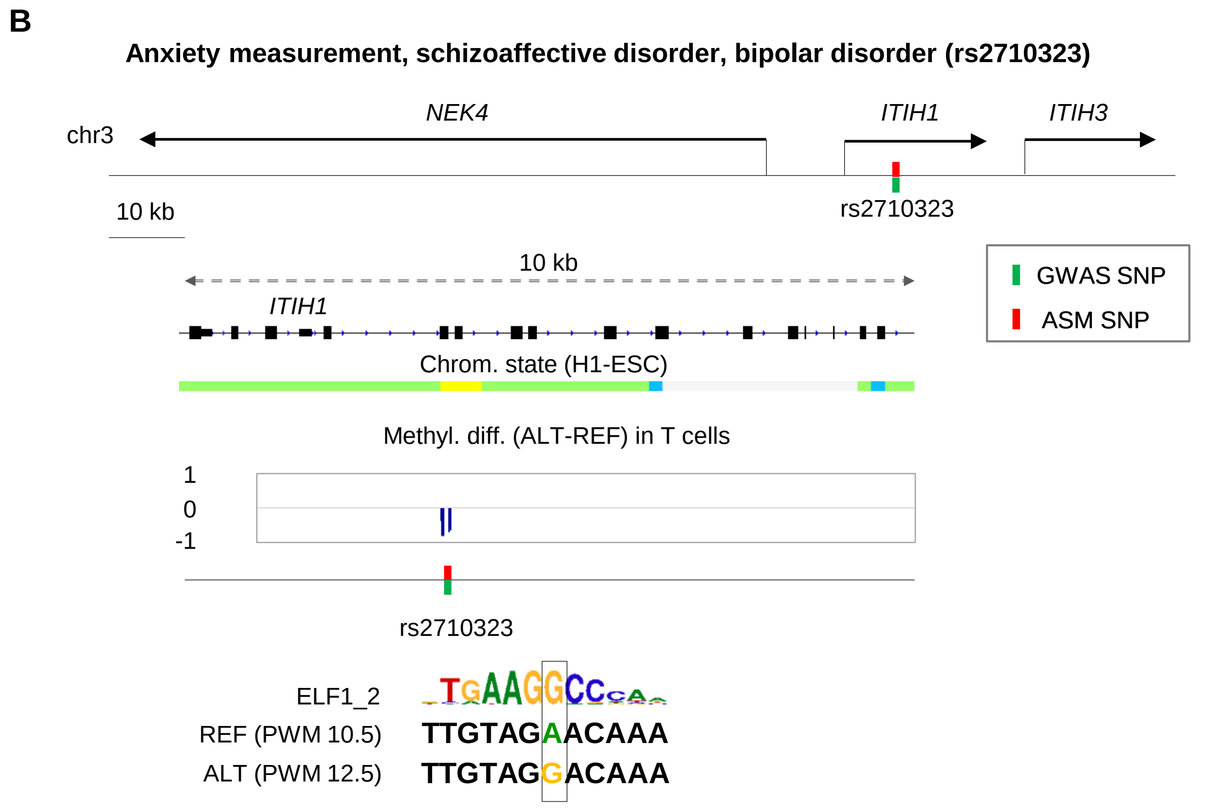
B
Anxiety measurement, schizoaffective disorder, bipolar disorder (rs2710323)
NEK4
ITIH1
ITIH3
chr3
10 kb
rs2710323
10 kb
ITIH1
GWAS SNP
ASM SNP
Chrom. state (H1-ESC)
Methyl. diff. (ALT-REF) in T cells
1
0
-1
rs2710323
ELF1_2
REF (PWM 10.5) TTGTAGAACAAA
ALT (PWM 12.5) TTGTAGGACAAA

#### Figure S23

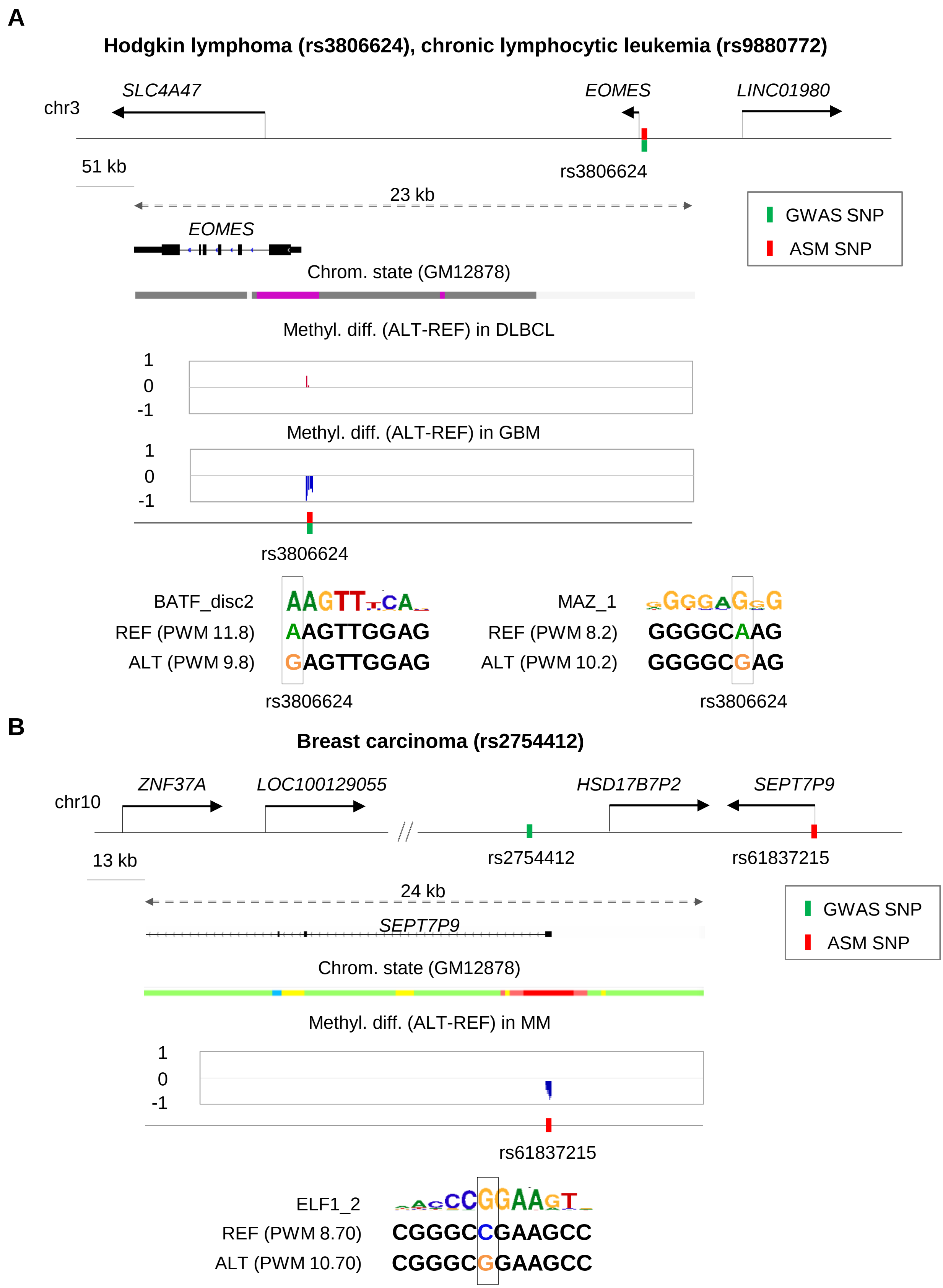
A
Hodgkin lymphoma (rs3806624), chronic lymphocytic leukemia (rs9880772)
chr3
SLC4A47
EOMES
LINC01980
51 kb
rs3806624
23 kb
GWAS SNP
ASM SNP
EOMES
Chrom. state (GM12878)
Methyl. diff. (ALT-REF) in DLBCL
1
0
-1
Methyl. diff. (ALT-REF) in GBM
1
0
-1
rs3806624
BATF_disc2
REF (PWM 11.8) AAGTTGGAG
ALT (PWM 9.8) GAGTTGGAG
rs3806624
MAZ_1
REF (PWM 8.2) GGGGCAAG
ALT (PWM 10.2) GGGGCGAG
rs3806624
B
Breast carcinoma (rs2754412)
chr10
ZNF37A
LOC100129055
HSD17B7P2
SEPT7P9
13 kb
rs2754412
rs61837215
24 kb
SEPT7P9
GWAS SNP
ASM SNP
Chrom. state (GM12878)
Methyl. diff. (ALT-REF) in MM
1
0
-1
rs61837215
ELF1_2
REF (PWM 8.70) CGGGCCGAAGCC
ALT (PWM 10.70) CGGGCGGAAGCC

#### Figure S24

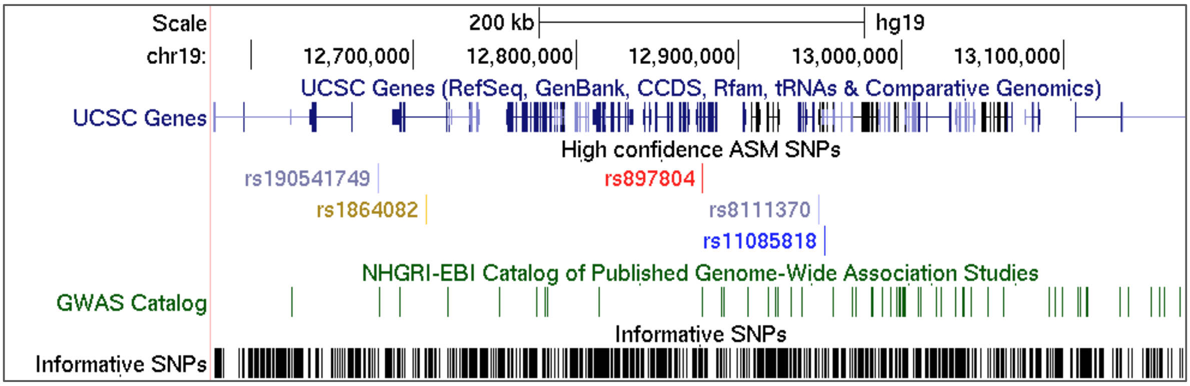


**Custom Track: ASM SNPs**

##### High confidence ASM SNPs

**Item:** rs897804
**Score:** 884
**Position:** chr19:12876964-12876964
**Band:** 19p13.2
**Genomic Size:** 1
View DNA for this feature (hg19/Human)

**ID:** chr19:12876798-12877138

General information about the ASM index SNP

#### ASM information for index SNP : rs897804

##### DMR infomation

- **DMR coordinates :** chr19:12876798-12877138
- **DMR overall rank :** 1
- **DMR strength rank :** 5
- **DMR confidence rank :** 45
- **SNP associated with DMR :** rs897804

Information about the ASM DMR

##### SNP level infomation

- **SNP overall rank :** 1
- **SNP strength rank :** 5
- **SNP confidence rank :** 45
- **Nb. samples with ASM :** 21
- **Nb. heterozygous samples :** 21
- **Switching ASM :** No

Ranking of the ASM index SNP

- **SNP color code :**

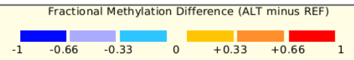


Color code for fractional methylation difference

| Sample ID | Cancer status | Cell/Tissue | Methylation Difference | FDR | Nb. CpGs with ASM | Nb. covered CpGs | Sequencing platform |
|---|---|---|---|---|---|---|---|
| Sample 101 | Cancer | Multiple Myeloma | .9 | 6.6e-04 | 15 | 17 | WGBS |
| Sample 12 | Non-Cancer | B Cells | 1 | 5.6e-05 | 18 | 18 | WGBS |
| Sample 17 | Non-Cancer | Bladder Epith Cells | 1 | 2.5e-04 | 17 | 17 | WGBS |
| Sample 22 | Non-Cancer | Brain Frontal Cortex | .7 | 1.2e-19 | 25 | 26 | WGBS |

Samples with ASM

#### Polymorphic motifs for rs897804

##### Enriched polymorphic motif

| Motif name | PWM score ALT allele | PWM score REF allele | Difference in PWM score | FDR for the difference in PWM score |
|---|---|---|---|---|
| ABF1_MA0570_1 | 10.6 | 12.5 | -1.9 | 6.8e-03 |
| BCL_disc10 | 4.6 | 4.1 | .5 | <5e-324 |
| CREB3L1_2 | 10.7 | 12.7 | -2 | 2.3e-02 |

##### Polymorphic motifs with methylation-binding affinity correlation

| Motif name | PWM score ALT allele | PWM score REF allele | Difference in PWM score | FDR for the difference in PWM score |
|---|---|---|---|---|
| CTCF_1 | 14.8 | 15.7 | -.8 | 4.5e-03 |
| CTCF_MA0139.1 | 14.8 | 15.6 | -.8 | 4.8e-03 |

Go to ASM SNPs track controls

**Data last updated:** 2019-07-26

Polymorphic TF binding motifs enriched among ASM loci and disrupted by the ASM index SNP
